# MIRA: Joint regulatory modeling of multimodal expression and chromatin accessibility in single cells

**DOI:** 10.1101/2021.12.06.471401

**Authors:** Allen W. Lynch, Christina V. Theodoris, Henry Long, Myles Brown, X. Shirley Liu, Clifford A. Meyer

## Abstract

Rigorously comparing gene expression and chromatin accessibility in the same single cells could illuminate the logic of how coupling or decoupling of these mechanisms regulates fate commitment. Here, we present MIRA: Probabilistic Multimodal <u>M</u>odels for <u>I</u>ntegrated <u>R</u>egulatory <u>A</u>nalysis, a comprehensive methodology that systematically contrasts transcription and accessibility to infer the regulatory circuitry driving cells along developmental trajectories. MIRA leverages topic modeling of cell states and regulatory potential modeling of individual gene loci. MIRA thereby represents cell states in an efficient and interpretable latent space, infers high fidelity lineage trees, determines key regulators of fate decisions at branch points, and exposes the variable influence of local accessibility on transcription at distinct loci. Applied to epidermal maintenance differentiation and embryonic brain development from two different multimodal platforms, MIRA revealed that early developmental genes were tightly regulated by local chromatin landscape whereas terminal fate genes were titrated without requiring extensive chromatin remodeling.

## Main

Profiling both expression and chromatin accessibility in the same single cells^1–5^ opens an unprecedented opportunity to understand the interaction of transcriptional and epigenetic mechanisms driving cells along developmental continuums. While many computational methods analyze expression and accessibility separately, several recent algorithms have adopted joint analysis where the cells are projected onto a shared latent space based on both data modalities, which better captures the biological structure of the data^6–11^. However, the field lacks tools that go beyond visualization and clustering to rigorously contrast transcription and accessibility in each single cell to illuminate the complex regulatory circuitry driving developmental fate decisions.

Integrated analysis of global transcriptional and accessibility states across developmental trajectories would enable discovery of key regulators controlling fate decisions at lineage branch points. At the gene level, examining the dynamics of transcription versus chromatin accessibility proximal to the gene locus may reveal how these mechanisms interact to regulate distinct gene modules. Certain genes may be regulated by cis-regulatory elements that are simultaneously activated as they become accessible, whereas others may be regulated by elements whose accessibility and activation are decoupled^12,13^. Determining the logic of which genes are regulated by each of these distinct mechanisms may provide insight into the patterns of pathways that demand tight spatiotemporal regulation versus signal responsivity.

Here, we present MIRA: Probabilistic Multimodal <u>M</u>odels for <u>I</u>ntegrated <u>R</u>egulatory <u>A</u>nalysis, a comprehensive methodology that systematically contrasts transcription and accessibility to determine the regulatory circuitry driving cells along developmental continuums. MIRA leverages topic modeling of cell states and regulatory potential (RP) modeling of individual gene loci. MIRA thereby represents cell states in an efficient and interpretable latent space, infers high fidelity lineage trees, determines key regulators of fate decisions at branch points, and exposes the variable influence of local accessibility on transcription at distinct loci. We applied MIRA to an epidermal maintenance differentiation^3^ and brain developmental system^14^ assayed by multimodal single cell RNA-sequencing (scRNA-seq) and Assay for Transposase-Accessible Chromatin-sequencing (scATAC-seq) data from two different platforms (SHARE-seq and 10x Genomics). In each system, MIRA constructed a high fidelity developmental trajectory and determined the regulatory factors driving key fate decisions at lineage branch points. Furthermore, MIRA distinguished early developmental genes that were tightly spatiotemporally regulated by local chromatin landscape from terminal fate genes that were permitted to remain accessible while titrated by factors with minimal impact on local chromatin, revealing how variable regulatory circuitry coordinates fate commitment and terminal identity.

## Results

### MIRA leverages topic modeling and RP modeling to reveal the circuitry regulating developmental trajectories

MIRA leverages topic modeling and RP modeling of expression and chromatin accessibility in single cells to determine the regulatory mechanisms driving key fate decisions within developmental continuums (Fig. 1a-b, Methods). Probabilistic topic modeling has been employed in natural language understanding to elucidate the abstract topics that shape the meaning of a given collection of text^15^. Recently, topic modeling has been applied to scRNA-seq and scATAC-seq separately to describe either transcriptional or epigenetic cell states^16,17^ as “thematic” groups of co-regulated genes or cis-regulatory elements, respectively.

**Fig. 1.**
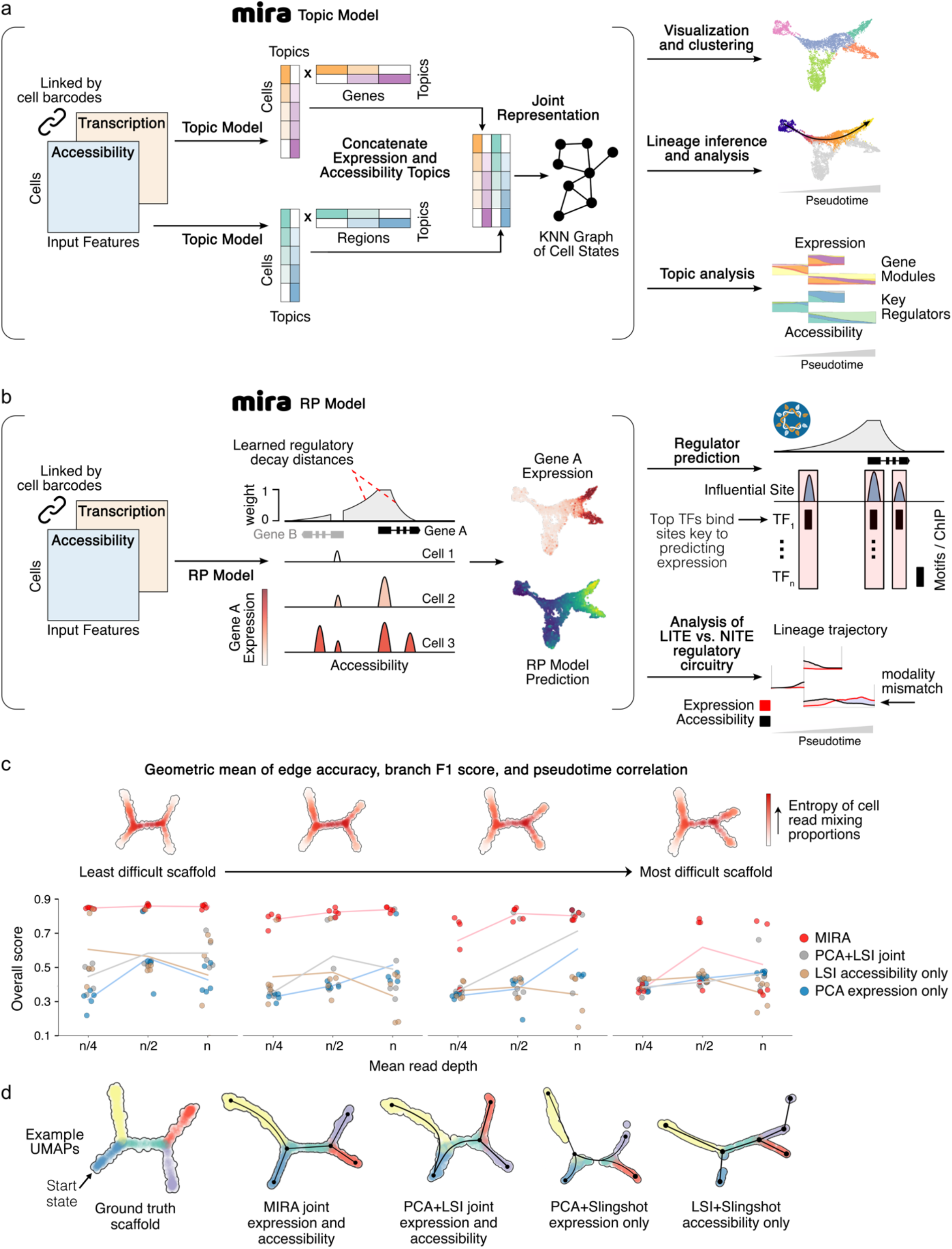
Schematic of MIRA’s cell-level topic and gene-level RP models for integrated analysis of single cell multimodal transcription and accessibility data. **a,** MIRA harnesses a variational autoencoder^18^ approach to model the transcription and chromatin accessibility topics defining each cell’s identity with a sparsity constraint to ensure topics are coherent and interpretable^19^. MIRA’s hyperparameter tuning scheme learns the appropriate number of topics needed to comprehensively yet non-redundantly describe each dataset. MIRA next combines the expression and accessibility topics into a joint representation used to calculate a k-nearest neighbors (KNN) graph. This output can then be leveraged for visualization and clustering, construction of high fidelity lineage trajectories, and rigorous topic analysis to determine regulators driving key fate decisions at lineage branch points. **b,** MIRA’s RP model integrates transcriptional and chromatin accessibility data at each gene locus to determine how regulatory elements surrounding each gene influence its expression. Regulatory influence of enhancers is modeled to decay exponentially with genomic distance at a rate learned by the MIRA RP model from the joint multimodal data. MIRA learns independent upstream and downstream decay rates and includes parameters to weigh upstream, downstream, and promoter effects. MIRA quantifies the regulatory influence of local chromatin state by contrasting the observed gene expression to that which would be predicted based solely on the local accessibility state (LITE model) or an expanded model that also includes the genome-wide accessibility state encoded by MIRA accessibility topics (NITE model). This quantification enables MIRA to distinguish genes primarily regulated by local chromatin remodeling (LITE genes) versus those more heavily influenced by non-local signals (NITE genes) reflected in the genome-wide accessibility topics with minimal impact on local chromatin landscape. MIRA furthermore predicts key regulators at each locus by examining transcription factor motif or occupancy (from ChIP-seq) enrichment within elements predicted to highly influence transcription at that locus via a pISD approach. **c,** Benchmarking results comparing MIRA joint lineage trajectory inference to standard methodology of Seurat principal component analysis (PCA)+Slingshot on expression data only, Seurat latent semantic indexing (LSI)+Slingshot on accessibility data only, or joint model combining Seurat PCA on expression data and LSI on accessibility data followed by Slingshot. Overall score is the geometric mean of edge accuracy, branch F1 score, and pseudotime correlation metrics. Performance was tested on four different ground truth scaffolds, which are computationally synthesized by mixing reads from distinct populations of single cells from a 10x Genomics dataset^47^ of peripheral blood mononuclear cells (PBMCs). Scaffold views calculated by Cartesian projection of per-cell read mixing proportions. Scaffold difficulty increases from left to right, where more difficult scaffolds contain lineages where mixture components are more similar. Line plots indicate performance in each of the four scaffold difficulties with trials for three different mean read depths (lower read depth further increases the difficulty of solving the topology). For expression data, mean read depth n=4000; for accessibility data, mean read depth n=14000. For each trial, 5 replicates were tested. *Edge accuracy* measures the accuracy of the inferred edges compared to ground truth (dynverse’s edge flip score^48^). *Branch F1 score*^48^ measures the precision and recall of the inferred branches compared to ground truth. *Pseudotime correlation*^48^ measures the correlation between inferred versus ground truth pseudotime for each cell. **d,** Example UMAPs for each modeling approach for the least difficult ground truth scaffold. Black edges show lineage parsing from each algorithm. Cells colored by ground truth branch assignment.

MIRA’s topic model uses a variational autoencoder^18^ approach, intersecting deep learning with probabilistic graphical models, to learn expression and accessibility topics defining each cell’s identity (Extended Data Fig. 1). MIRA accounts for the distinct statistical properties of each modality by using different generative distributions for overdispersed scRNA-seq counts and sparse scATAC-seq data. A sparsity constraint is employed to ensure cells’ topic compositions are coherent and interpretable^19^. MIRA’s Bayesian hyperparameter tuning scheme finds the appropriate number of topics needed to comprehensively yet non-redundantly describe each dataset.

MIRA next combines the expression and accessibility topics into a joint representation used to calculate a k-nearest neighbors (KNN) graph. The KNN graph is then leveraged to construct a lineage tree using a new method we developed to define the branch points between lineages where the probabilities of differentiating into one terminal state diverges from another (Extended Data Fig. 2). A benchmarking comparison of MIRA’s lineage tree construction demonstrated consistently better performance than standard alternatives (Fig. 1c-d, Extended Data Fig. 3-5). MIRA then contrasts the emergence of expression and accessibility topics mapped on this lineage tree to elucidate the key regulators driving fate decisions at the inferred lineage branch points.

Next, MIRA leverages RP modeling^20,21^ to integrate transcription and accessibility at the resolution of individual gene loci to determine how regulatory elements surrounding each gene influence its expression (Fig. 1b). While correlation between chromatin accessibility and expression is confounded by coordinated genome-wide changes ascribed to cell state, genomic proximity suggests a mechanistic regulatory relationship between cis-regulatory elements and transcription. Thus, the perceived influence of cis-regulatory elements is modeled to decay exponentially with genomic distance upstream or downstream of a transcriptional start site (TSS) at independent rates learned by MIRA from the multimodal data. Each gene’s RP is scored as the sum of the contribution of individual regulatory elements. MIRA predicts key regulators at each locus by examining transcription factor motif enrichment or occupancy (if provided chromatin immunoprecipitation-sequencing (ChIP-seq) data) within elements predicted to highly influence transcription at that locus by probabilistic *in silico* deletion (pISD).

Furthermore, MIRA quantifies the regulatory influence of local chromatin accessibility on gene expression by comparing the local RP model with a second, expanded model augmented with knowledge of genome-wide accessibility states encoded by MIRA’s accessibility topics. Genes whose transcription is sufficiently predicted by the RP model based on local accessibility alone (±600 kilobases from the TSS) are defined as local chromatin accessibility-influenced transcriptional expression (LITE) genes. Genes whose expression is significantly better described by the model with genome-wide scope are defined as non-local chromatin accessibility-influenced transcriptional expression (NITE) genes.

While LITE genes appear tightly regulated by local chromatin accessibility, the transcription of NITE genes appears to be titrated without requiring extensive local chromatin remodeling. MIRA defines the extent to which the LITE model over- or under-estimates expression in each cell as “chromatin differential”, highlighting cells where transcription is decoupled from shifts in local chromatin accessibility. MIRA examines chromatin differential across the developmental continuum to reveal how variable circuitry regulates fate commitment and terminal identity.

### MIRA topic modeling determined regulators driving key fate decisions in hair follicle differentiation

Applied to hair follicle maintenance differentiation assayed by SHARE-seq^3^, MIRA’s joint topic representation constructed a lineage map whose latent structure mimicked the follicle’s true spatial layout^22,23^ (Fig. 2a, Extended Data Fig. 6). MIRA’s inferred lineage tree reconstructed the ancestral hierarchy of follicular lineages, with outer root sheath cells leading to early matrix progenitors, which subsequently branched into descendant inner root sheath (IRS) followed by medulla and cortex lineages (Fig. 2b). Accurate lineage trees are a crucial prerequisite to determining the factors directing cell fate decisions at lineage branch points.

**Fig. 2.**
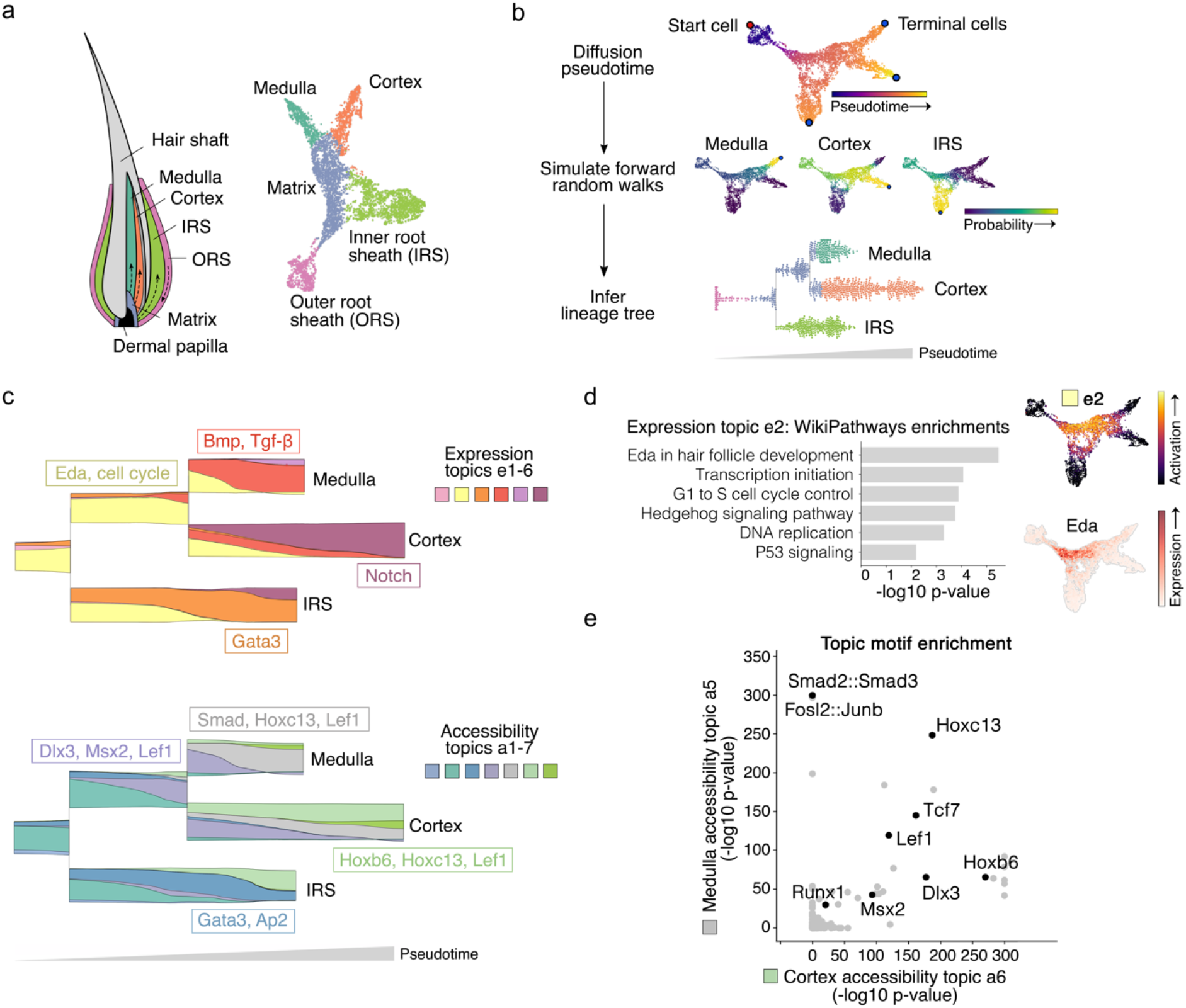
MIRA topic modeling determined regulatory factors driving key fate decisions in hair follicle differentiation. **a,** MIRA’s joint topic representation constructed a UMAP *(right)* whose structure mimicked the true spatial layout^22^ *(left)* of the progenitor matrix cells and descendant medulla, cortex, and IRS lineages in the hair follicle. Colors indicate cell types defined by fine Leiden clustering followed by agglomeration of clusters based on known marker gene expression. **b,** *(top)* Diffusion pseudotime through joint KNN graph representing differentiation progress. Terminal cells were identified using stationary states from a forward Markov chain model of differentiation. *(middle)* Each cell’s probability of reaching each terminal state. *(bottom)* Parsed bifurcating tree structure of lineage probabilities visualized as stream graph. Each point is an individual cell arranged as a swarm plot (arranged such that points do not overlap, resulting in larger spread where there are more points). Cells are colored by clusters in 2a, indicating that bifurcation points closely correspond to changes in cell identity as separately defined by markers for each cell type. **c,** Stream graph of window-averaged cell-topic compositions as cells progress through differentiation starting from matrix cell state (see Extended Data Fig. 7a for stream graph including outer root sheath (ORS); topics that comprise ≥3% of the total at any point are shown). Representative genes activated in expression topics and motifs enriched in accessibility topics are depicted in boxes corresponding with the color of the source topic. Accessibility topic a4 described a transitory accessibility state at the branch point between the medulla and cortex lineages without a corresponding expression topic, suggesting global chromatin remodeling in progenitor matrix cells preceded transcriptional alterations specifying each downstream lineage. **d,** *(left)* Gene set enrichment for progenitor matrix cell expression topic e2. *(right)* Expression topic e2 activation or Eda expression on UMAP of joint topic representation. **e,** Comparison of motif enrichment in top peaks of medulla versus cortex accessibility topics (a5 and a6, respectively).

MIRA contrasts the flow of expression and accessibility topics across the inferred lineage tree using stream graphs^24^ to expose the regulatory modules driving cell fates along distinct developmental paths (Fig. 2c; Extended Data Fig. 7a-b, Table 1-2). Stream graphs enable high-dimensional, multimodal comparisons along continuums. Expression topic e2 captured the transcriptional state governing progenitor matrix cells, including cell proliferation^25^ and Eda and Shh signaling^22,26^ (Fig. 2d). Thereafter, expression topic e6 described cortex specification corresponding with activation of Notch-associated factors^27^. Conversely, expression topic e4 characterized medulla specification, containing Bmp/Tgf-β-associated factors^22^ aligned with enrichment of Smad5/Smad2/3 motifs in medulla-specific accessibility topic a5 (Fig. 2e, Extended Data Fig. 7c). Comparison with cortex-specific accessibility topic a6 showed both lineages were enriched for motifs bound by canonical hair shaft regulators Lef1 and Hoxc^22^, with expression implicating the influence of Hoxc13 (Extended Data Fig. 8a).

Contrasting modalities, Wnt-driven accessibility topic a4 described a transitory accessibility state at the branch point between the medulla and cortex lineages without a corresponding expression topic (Fig. 2c, Extended Data Fig. 8b-c). Cell-level chromatin remodeling in progenitor matrix cells thus preceded transcriptional alterations specifying each downstream lineage.

### MIRA RP modeling distinguished LITE versus NITE genes in the hair follicle

While most genes in the hair follicle exhibited LITE regulation with local accessibility increasing synchronously with transcription, expression diverged from that predicted by the LITE model for genes such as Krt23 (Fig. 3a-d, Extended Data Fig. 8d-e). Although local chromatin accessibility was poorly predictive of Krt23 expression, its transcription was lineage-specific and closely aligned with activation of accessibility topic a5, encoding a medulla genome-wide pattern of accessibility. Consistently, Krt23 expression was more closely predicted by the NITE model which includes these genome-wide accessibility states as features (Fig. 3e). LITE genes are thus tightly regulated by local chromatin remodeling, whereas NITE genes are titrated without requiring extensive local chromatin remodeling, decoupling transcription from local accessibility (Fig. 3f).

**Fig. 3.**
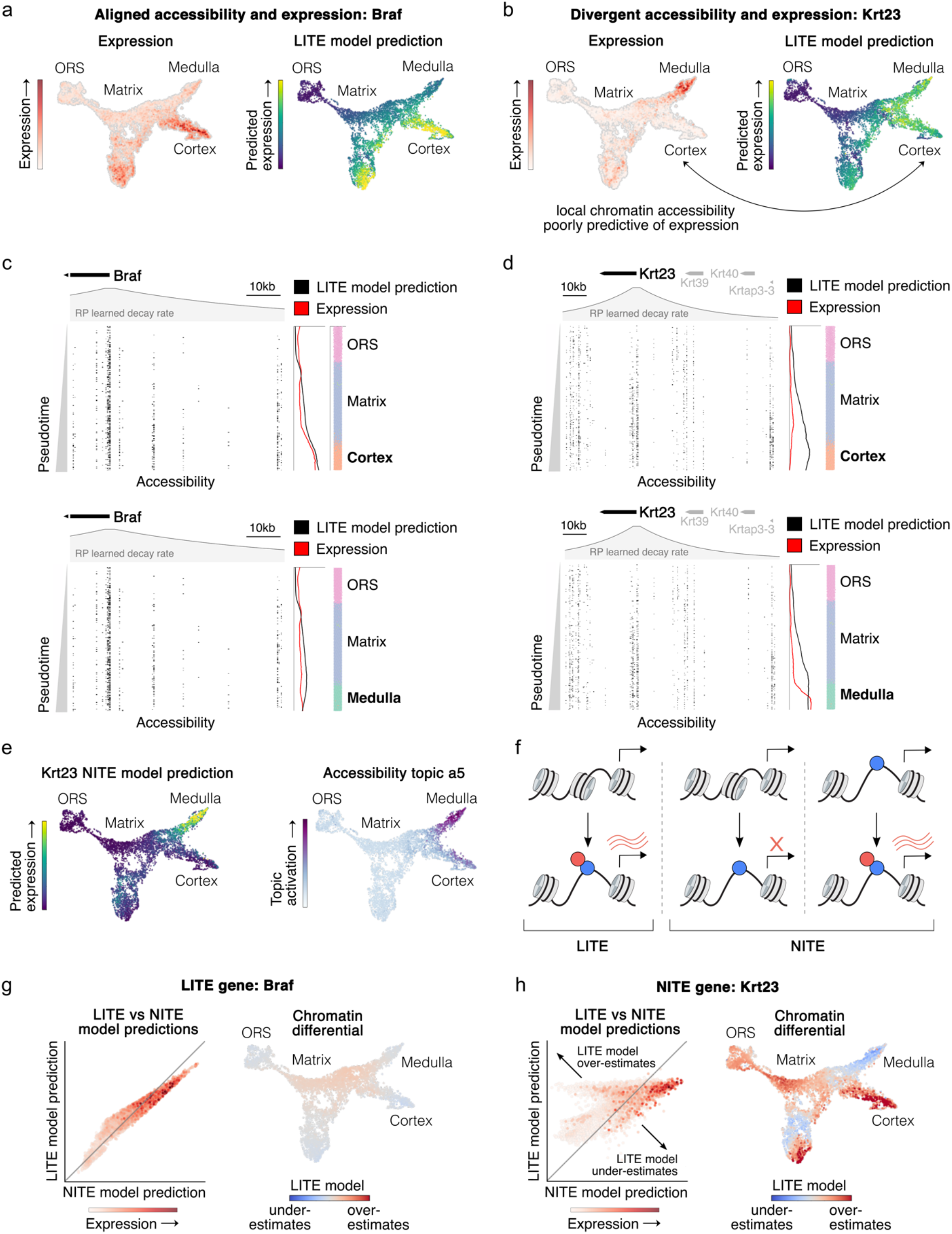
MIRA RP modeling identified genes for which changes in expression were insufficiently explained by local chromatin accessibility. **a,** LITE gene Braf or **b,** NITE gene Krt23 expression or local-only RP model predictions (LITE model) on joint representation UMAP. **c,** LITE gene Braf or **d,** NITE gene Krt23 locus’s chromatin accessibility fragments across pseudotime (moving downwards) in trajectories from ORS to matrix to cortex *(top)* or medulla *(bottom)*. Colored bars on the right indicate the identity of cells (colored by clusters in 2a) within each bin reflected by each row of accessibility fragments. Line plots across pseudotime depict the indicated gene’s observed expression (red) and LITE model prediction of expression (black), which is informed by the local accessibility reflected in the fragment plot. While the observed expression and LITE model prediction align for LITE gene Braf, they diverge for NITE gene Krt23. **e,** Joint representation UMAP colored by *(left)* Krt23 NITE model prediction or *(right)* medulla accessibility topic a5 capturing a genome-wide chromatin state. NITE model predictions were more closely aligned with Krt23 expression shown in 3b. **f,** Proposed mechanism of LITE versus NITE regulation. In LITE regulation, expression is tightly regulated by chromatin remodeling. In NITE regulation, binding of an additional factor is required to enact transcription. **g,** LITE gene Braf or **h,** NITE gene Krt23 LITE versus NITE model predictions (cells colored by gene expression) and “chromatin differential” (relative prediction of LITE versus NITE models). In chromatin differential plots, red indicates LITE model overestimates expression while blue indicates LITE model under-estimates expression relative to NITE model.

MIRA’s “chromatin differential” mapped the extent to which local accessibility was decoupled from transcription across the developmental trajectory (Fig. 3g-h). Although Krt23 local accessibility increased at the branch point between the medulla and cortex lineages and remained elevated in both, it was ultimately only highly expressed in the medulla, causing high chromatin differential that over-estimated its expression in the cortex. Krt23’s lineage-specific expression despite accessibility in both lineages suggests its activation requires addition of a factor that does not primarily impact transcription via remodeling local accessibility.

### MIRA analysis of NITE regulation elucidated hair follicle fate commitment mechanism

At the cell level, gene expression in terminally-differentiated medulla and cortex cells exhibited significantly more NITE regulation than gene expression earlier in hair follicle differentiation (p<0.05, Wilcoxon) (Fig. 4a, Extended Data Fig. 8f). Often, accessibility of terminally-expressed genes increased before fate commitment and was maintained in both subsequent lineages, but expression activated in a lineage-specific manner only after the branch point between medulla and cortex (Fig. 4b-c, Extended Data Fig. 8g). We used chromatin differential at the branch point to identify genes with these “branch-primed” dynamics. While priming suggests the inevitability of expression, these genes indicate subsequent expression at primed loci can be conditional, a pattern detected as strong NITE regulation.

**Fig. 4.**
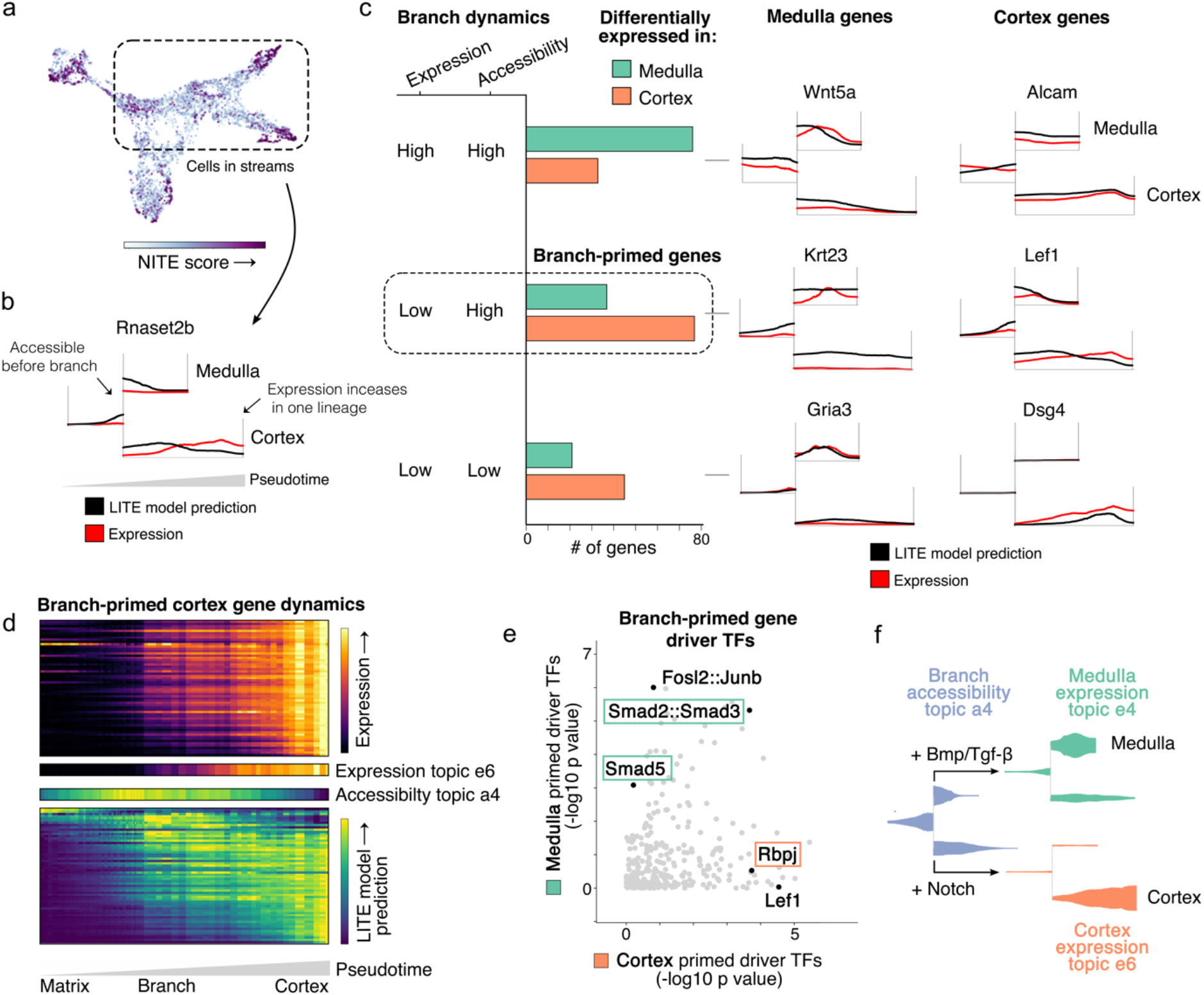
Gene-level and cell-level analysis of NITE gene regulation in the hair follicle elucidated regulatory mechanisms of fate commitment. **a,** NITE regulation score of each cell. Genome-wide expression in darker cells appeared to be more highly influenced by factors with minimal impact on local chromatin. **b,** Stream graph contrasting expression versus LITE model prediction (which represents local accessibility) of NITE gene Rnaset2b. Unlike LITE genes where expression and local accessibility (and therefore the LITE model prediction) change over time in a coordinated manner, NITE genes such as Rnaset2b show a divergence between observed expression and LITE model prediction in the stream graphs. When the lines diverge, this indicates that observed expression or local accessibility is changing in a way that is not coordinated with the other, drawing attention to genes whose expression is regulated by mechanisms not solely determined by local chromatin accessibility. **c,** *(left)* Regulatory classifications of medulla or cortex terminally expressed genes based on expression and local chromatin accessibility at the branch point between medulla and cortex lineages. Classifications colored green or orange by whether the genes were significantly upregulated in the medulla or cortex cells, respectively. The High Expression-High Accessibility group is composed of medulla- or cortex-specific genes that are already highly expressed and accessible at the branch. The Low Expression-High Accessibility group, referred to as “branch-primed genes”, are medulla- or cortex-specific genes that are more accessible at the branch than would be expected based on their expression at the branch. They subsequently increase in expression levels after the branch in one of the two lineages. The Low Expression-Low Accessibility group, referred to as “terminal genes”, are medulla- or cortex-specific genes that are not yet expressed nor accessible at the branch. Only after the cells have committed to one of the two fates do these genes become expressed and accessible in that lineage. *(right)* Example of each classification. **d,** Interaction between gene-level regulation and cell-level topics. *(top)* Expression of branch-primed cortex genes increased after branch, correlating with expression topic e6. *(bottom)* LITE model prediction (local chromatin accessibility) of branch-primed genes increased before cortex commitment, correlating with accessibility topic a4. **e,** Driver transcription factor analysis of branch-primed medulla versus cortex genes. Branch-primed cortex genes showed regulation by Rbpj, effector of Notch signaling, while branch-primed medulla genes showed regulation by Smad5 and Smad2/3, effectors of Bmp and Tgf-β signaling. **f,** Model for regulation of fate commitment in hair follicle depending on activation of distinct signaling pathways. Accessibility topic a4 opens chromatin around branch-primed genes at branch point between lineages. Depending on signal, branch-primed lineage-specific genes are expressed, enforcing lineage commitment.

Cell-level topic modeling also supported the pattern of primed accessibility preceding fate commitment. For example, the dynamics of genes ultimately expressed in the cortex whose accessibility was primed at the preceding branch point were described by cortex-specific expression topic e6 and branch-spanning accessibility topic a4 (Fig. 4d). As previously noted, accessibility topic a4 described a cell-wide change in chromatin state that did not correspond with a synchronous change in expression topic influence.

Branch-primed genes that were subsequently conditionally expressed in medulla or cortex appeared to respond to a regulator of medulla or cortex fate commitment. MIRA pISD implicated Notch effector Rbpj as a top regulator of branch-primed cortex genes and Bmp/Tgf-β-induced Smad5/Smad2/3 as regulators of branch-primed medulla genes (Fig. 4e, Extended Data Fig. 8h), consistent with expression of genes associated with these factors’ induction (Extended Data Fig. 7b). Thus, MIRA determined that cells at the branch point have a chromatin state permissible to multiple fates, described by transitory accessibility topic a4, ultimately driven to medulla or cortex through the subsequent addition of a fate-defining signal, namely Bmp/Tgf-β or Notch^22,27,28^ (Fig. 4f).

Overall, leveraging MIRA to systematically contrast expression and accessibility at single cell and locus resolution in the hair follicle revealed the fate commitment mechanism regulating the medulla and cortex lineages.

### MIRA captured two distinct spatiotemporal axes of differentiation in the interfollicular epidermis

We next applied MIRA to a separate system in the same dataset^3^, the interfollicular epidermis (IFE). Two spatial axes of differentiation specify the IFE, one controlling the differentiation of basal stem cells into increasingly superficial epidermal layers (epidermal stratification axis) and another controlling basal cell invagination and follicular formation (follicular axis)^29^ (Fig. 5a). The latent structure of MIRA’s joint topic representation again mimicked the spatial layout of this differentiation system, reconstructing the two axes of differentiation (Fig. 5b, Extended Data Fig. 9-10).

**Fig. 5.**
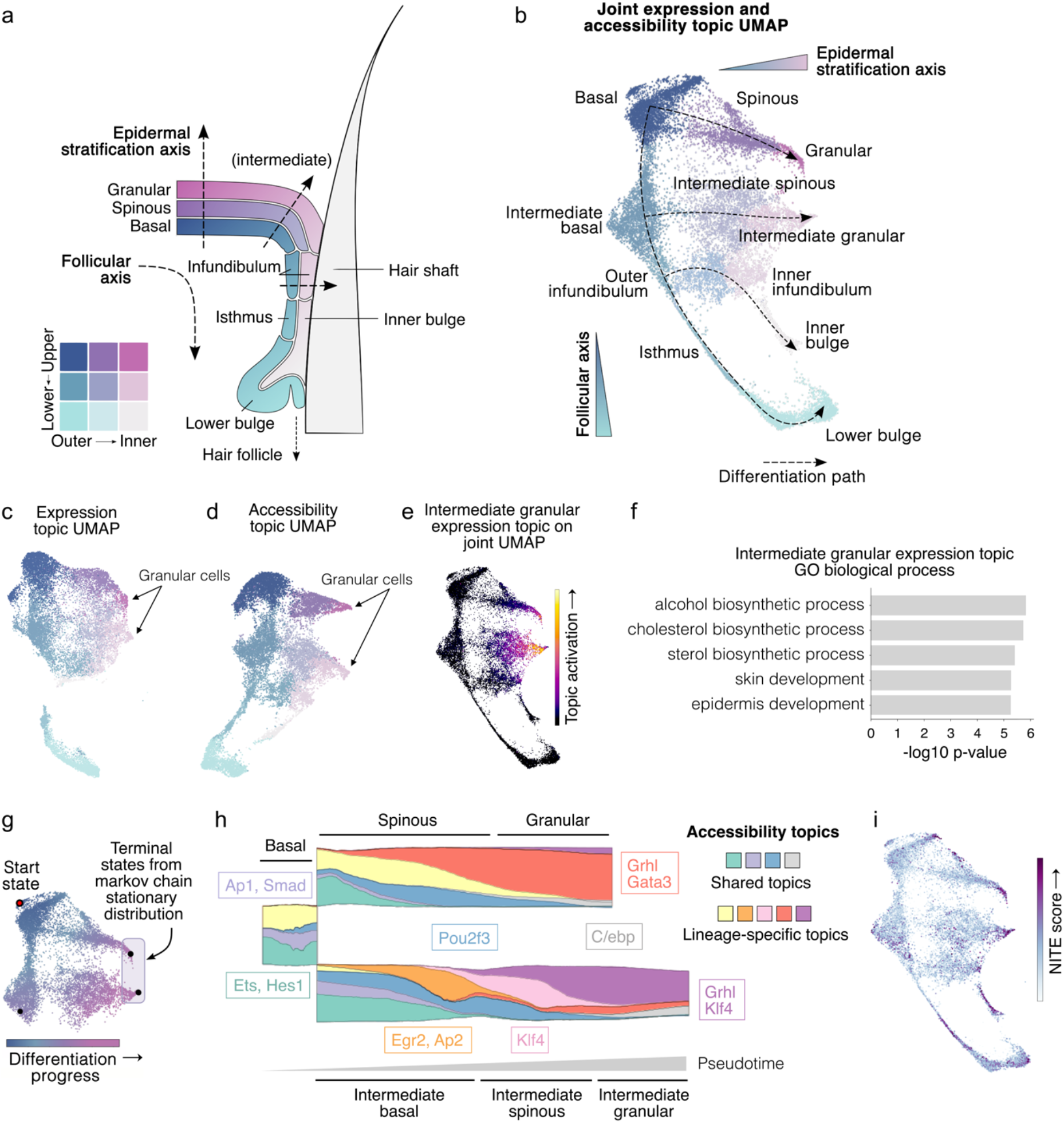
MIRA joint representation reconstructed complex multi-axis differentiation in the IFE. **a,** Anatomical model of mouse keratinocyte differentiation along epidermal and follicular axes. **b,** UMAP calculated from MIRA joint expression and accessibility topic representation. Dotted lines show constructed lineage structure resulting from two axes of differentiation. **c,** UMAP calculated from MIRA expression topics alone. **d,** UMAP calculated from MIRA accessibility topics alone. **e,** Activation of intermediate granular expression topic e8 on joint representation UMAP. **f,** Gene Ontology (GO) enrichment of top genes from intermediate granular expression topic e8. **g,** Two separate terminal states were identified from the Markov chain model of differentiation starting from basal cells labeled “Start state”. **h,** Stream graph of accessibility topic compositions of basal-spinous-granular *(top)* and intermediate basal-spinous-granular *(bottom)* lineages. Top enriched factors shown in boxes with color indicating source topics. **i,** NITE regulation score of each cell in the IFE.

Furthermore, unlike prior reported analysis of this dataset^3^ that did not jointly model expression and accessibility, MIRA identified two distinct basal-spinous-granular trajectories. One trajectory, labeled “intermediate”, was more transcriptionally and epigenetically similar to upper hair follicle structures, suggesting these cells were spatially proximal to the hair follicle and subject to more pro-follicular regulation. These “intermediate” basal cells showed activation of Egr2 expression and motifs, previously implicated in epidermal proliferation and wound healing^30^ (Extended Data Fig. 11a-b). By contrast, basal cells distant from the hair follicle showed stronger expression of Thbs1, consistent with prior work^29^ that identified two distinct populations of basal cells with Thbs1 marking those distant from the hair follicle. Each of these two distinct basal cell niches produced their own columns of epidermal strata, which was captured by MIRA joint modeling.

Notably, the UMAP projection based only on expression obfuscated these distinct trajectories (Fig. 5c). RNA features were sufficient to distinguish the multi-stage transitions governing each basal-spinous-granular transformation but could not detect the lineage histories of each population. The accessibility-only representation, however, successfully aligned cells along lineages according to their distinct epigenetic characteristics (Fig. 5d). Projected together, the joint representation preserved the structure of the accessibility mode while integrating information of shared transcriptional identity from expression topics (Fig. 5b). In particular, expression topic e13 established cells with granular identities and captured co-upregulation of hallmark genes^29^ marking epidermal terminal differentiation (Fig. 5e-f).

The lineage tree inferred from the joint topic representation revealed both the shared and lineage-specific regulators shaping the spatial programs of the two basal-spinous-granular trajectories (Fig. 5g-h; Extended Data Fig. 11c, Table 3-4). Visualizing state changes through accessibility topics identified the shared regulatory influence of Hes on basal cells, followed by Pou2f3 on spinous cells and terminating in Grhl and C/ebp^23,31^ on granular cells in both trajectories. By contrast, lineage-specific accessibility topics distinguished the influence of Klf4 motifs^23^ in “intermediate” spinous and granular cells, as opposed to Gata3 influence^32^ in granular cells arising from Thbs1+ basal cells more distant from the hair follicle.

We observed that expression in terminal populations was significantly enriched for NITE regulation, especially in terminal genes differentially-expressed between lineages (p<0.05, Wilcoxon) (Fig. 5i, Extended Data Fig. 11d-f). Again, terminal fate chromatin accessibility appeared to specify the available cell states, while transcription ultimately depended on additional spatial or signaling queues. Overall, MIRA elucidated the shared and lineage-specific mechanisms of differentiation along two parallel trajectories with distinct spatial regulation within the IFE.

### MIRA elucidated regulators driving key fate decisions in embryonic brain development

We next applied MIRA to an E18 mouse embryonic brain dataset including cortex, hippocampus, and ventricular zone (assayed on a different platform, 10x Genomics Multiome)^14^ to determine the key factors driving astrocyte, excitatory neuron, and inhibitory neuron fates, the balance of which is critical for normal brain development. MIRA topic modeling constructed a joint representation with Pax6+ radial glia-like cells located centrally between the astrocyte, excitatory neuron, and inhibitory neuron branches (Fig. 6a-b). Pax6 marks both dorsal progenitors that give rise to astrocytes and excitatory neurons and the anatomically juxtaposed ventral progenitors in the lateral ganglionic eminence that give rise to inhibitory neurons^33,34^. Both of these progenitor populations were present within the 10x Genomics dataset and co-located within the joint representation due to their shared transcriptional state (Extended Data Fig. 12). MIRA topic analysis revealed the regulators driving the fate decision between astrocytes and excitatory neurons and furthermore identified regulators that contribute to the rise of excitatory versus inhibitory neurons from anatomically separated progenitors with a similar transcriptional state (Fig. 6c, Extended Data Fig. 13-16).

**Fig. 6.**
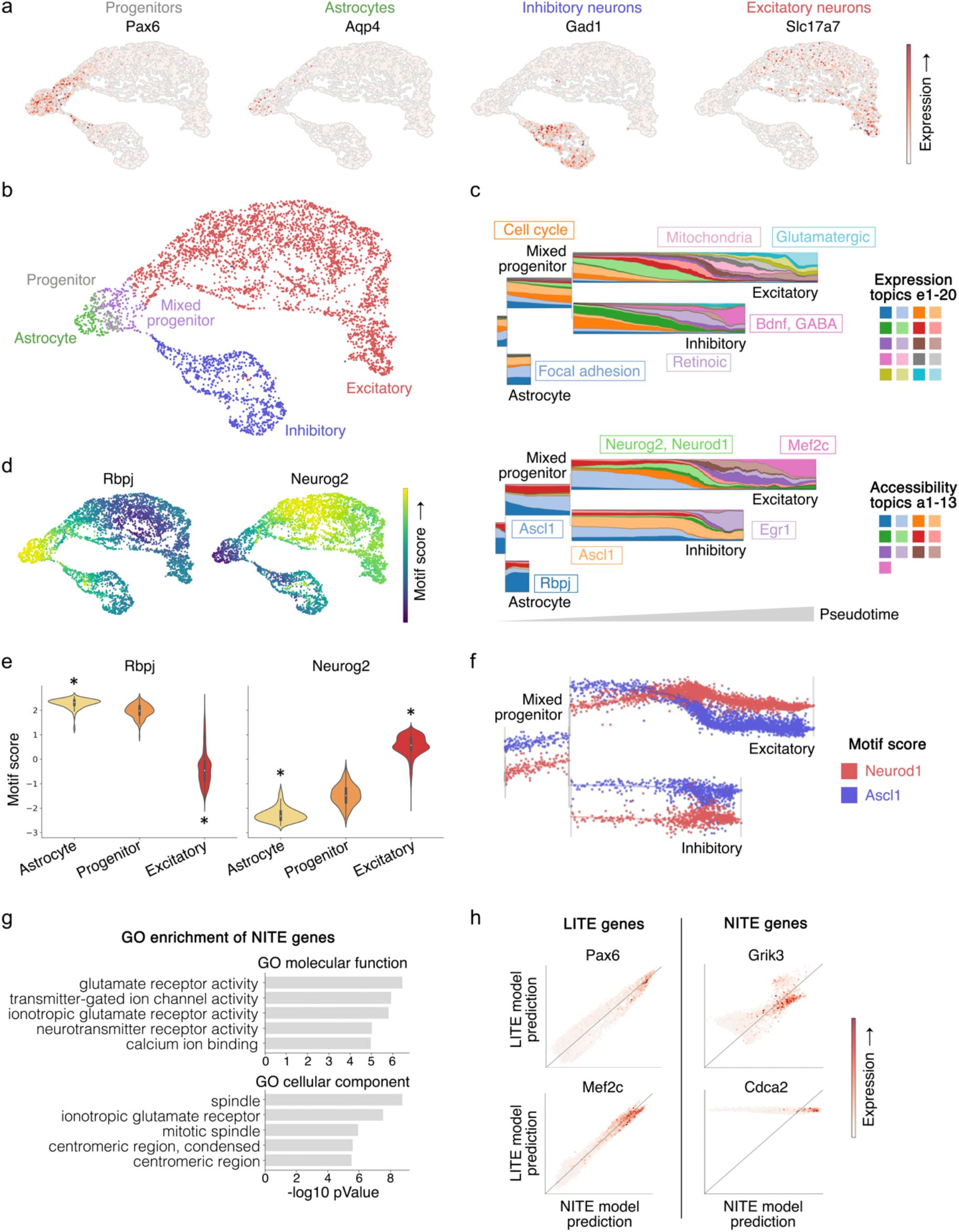
MIRA elucidated regulatory factors driving fate decisions in key developmental trajectories in the developing brain. **a,** Expression of marker genes for progenitor Pax6+ cells and terminal states of astrocytes, excitatory neurons, or inhibitory neurons. **b,** MIRA joint representation UMAP colored by inferred lineages. Mixed progenitor cells include both excitatory and inhibitory progenitors, which are transcriptionally similar. (See Extended Data Fig. 12). **c,** Stream graphs of expression and accessibility topic activation across pseudolineage trajectory. Pathways activated in expression topics and motifs enriched in accessibility topics are indicated by topic color. **d,** Motif score of Rbpj and Neurog2 on joint representation UMAP. **e,** Motif score of Rbpj and Neurog2 in the indicated lineages (*p<0.05, Wilcoxon compared to progenitors (including mixed progenitors), Benjamini-Hochberg-corrected). **f,** Activation level of Ascl1 versus Neurod1 motif scores in each single cell along pseudolineage trajectory. **g,** GO terms enriched in top 500 genes with NITE regulation where local chromatin accessibility state is insufficient to predict expression. **h,** Correlation of LITE versus NITE model predictions of expression of example genes with LITE versus NITE regulation.

MIRA analysis revealed that astrocytes were defined by accessibility topic a1, which was significantly enriched for Rbpj motifs associated with Notch signaling (Fig. 6c-e, Extended Data Table 5-6). Rbpj motifs were also enriched in Pax6+ progenitors, but significantly depleted from the excitatory neuron branch. Conversely, early excitatory accessibility topic a6 was significantly enriched for Neurog2 and Neurod1 motifs. These findings are consistent with prior developmental studies indicating that Notch signaling maintains progenitor multipotency and specification towards astrocytes while Neurog2 and Neurod1 commit cells to the excitatory neuron fate^35–37^.

We then investigated how anatomically separated progenitors with a shared transcriptional state differentially give rise to excitatory versus inhibitory neurons. The major expression topic e3 defining both trajectories was enriched for cell cycle genes, likely reflecting the expansion of progenitors prior to commitment to their terminal neuron fates^38^ (Extended Data Fig. 15, Table 5). Subsequent inactivation of cell cycle topic e3 aligned with activation of accessibility topic a2, which was enriched for motifs of Ascl1, a pioneering transcription factor in neural progenitors known to promote cell cycle exit and differentiation^38^ (Fig. 6c and f; Extended Data Fig. 15, Table 6). Ascl1 motifs were also enriched in the early inhibitory accessibility topic a4, consistent with Ascl1’s key role in inhibitory neuron differentiation^37^.

In the alternative trajectory towards the excitatory fate, early excitatory accessibility topic a6 demonstrated depletion of Ascl1 motifs coordinated with increased Neurod1 motifs (Fig. 6c and f, Extended Data Table 6). Ascl1 and Neurod1 belong to separate subgroups of basic helixloop-helix transcription factors^39^; and Neurod1 promotes differentiation of induced pluripotent stem cells into excitatory neurons, while Ascl1 specifies inhibitory neurons^37^.

MIRA topics contrasted the temporal progression of specification initiated by inhibitory-driving Ascl1 or excitatory-driving Neurod1. The inhibitory trajectory activated inhibitory maturation-driving Bdnf signaling^40^, culminating in the activation of GABA synapse components that define the terminal inhibitory fate (topic e13) (Fig. 6c; Extended Data Fig. 15, Table 5-6). Consistently, aligned terminal inhibitory accessibility topic a10 was enriched for Egr1 motifs, a downstream Bdnf effector that directly activates GABAergic neurotransmission genes^41^.

The diverging excitatory branch first activated mitochondrial components important for supporting neuronal metabolic demands^42^ (topic e14) followed by terminal activation of glutamatergic synapse machinery, including glutamate transporters which uniquely distinguish excitatory neurons (topic e20) (Fig. 6c; Extended Data Fig. 16a, Table 5). Aligned terminal excitatory accessibility topic a13 was enriched for Mef2 motifs attributable to Mef2c given its expression in the excitatory branch, consistent with its known role in maintaining the excitatory/inhibitory balance by promoting excitatory differentiation^43^ (Fig. 6c; Extended Data Fig. 16b, Table 6).

In summary, contrasting expression and accessibility topics on MIRA’s joint representation identified regulators driving key cell fate decisions in the developing brain and demonstrated the temporal progression of specification into inhibitory or excitatory neuronal fates.

### MIRA revealed LITE and NITE genes in the embryonic brain

To determine LITE and NITE genes in the embryonic brain, we trained MIRA RP models for the genes defining each expression and accessibility topic (Fig. 6g-h). Notable LITE genes included fate-driving transcription factors with tight spatiotemporal regulation such as progenitor gene *Pax6* and excitatory-promoting *Mef2c*. Conversely, NITE genes were enriched for cell cycle machinery as well as neuronal differentiation gene batteries composed of neurotransmitter and ion channel genes. Local chromatin landscape has been previously reported to have limited contribution to the activation of cell cycle genes^3^, consistent with NITE regulation. This may reflect a requirement for titration of genes governing each cell cycle stage that would be incompatible with the time needed to remodel the local chromatin landscape. Similarly, synaptic maintenance and plasticity may require fast-response regulation of neurotransmitter and ion channel genes, reflected as NITE regulation.

Analogously to the hair follicle and IFE, expression topics describing progenitors were significantly enriched for LITE regulation, whereas after commitment to the excitatory or inhibitory fate, topics were significantly enriched for NITE regulation (p<0.05, Wilcoxon) (Extended Data Fig. 16c). Progenitor and early inhibitory regulator Ascl1 is known to be a pioneering transcription factor that remodels the chromatin landscape to regulate its targets^38,44^. By contrast, terminal inhibitory regulator Egr1 was previously reported to have non-pioneer-like properties^45^. Notably, targets predicted by MIRA pISD to be downstream of Ascl1 demonstrated significantly stronger LITE regulation than predicted Egr1 targets, potentially reflective of local chromatin remodeling by pioneering Ascl1 driving their expression (p<0.05, Wilcoxon) (Extended Data Fig. 16d).

## Discussion

In sum, MIRA leverages cell-level topic modeling and gene-level RP modeling to rigorously contrast the spatiotemporal dynamics of single cell transcription versus chromatin accessibility to reveal how these mechanisms interact to orchestrate key fate decisions in developmental trajectories. MIRA demonstrated the power of topic modeling of expression and accessibility data to infer high fidelity lineage trees that consistently outperformed standard alternatives in benchmarking. Mapping expression and accessibility topics onto MIRA’s joint lineage tree illuminated the key regulators driving fate decisions at pivotal lineage branch points.

MIRA contrasted the dynamics of transcription and local chromatin accessibility to define the chromatin differential at each gene locus, revealing discrete gene modules regulated by primarily LITE or NITE mechanisms. Intriguingly, in all three systems that we tested from the skin^3^ and brain^14^ datasets, earlier-expressed genes were enriched for LITE regulation. LITE regulation of earlier-expressed genes may reflect the importance of strict regulation requiring extensive chromatin remodeling for their expression followed by strong silencing in fates where their aberrant expression would have devastating consequences. Conversely, gene batteries important for maintaining terminal cell function were less reliant on local chromatin remodeling for their regulation, suggesting larger influence by mechanisms such as cell signaling that allow titration of transcription to fulfill fluctuating cell needs.

Among NITE-regulated genes, we also noted genes with primed accessibility at lineage branch points that showed subsequent lineage-specific activation in response to a fate-defining force such as signaling, presumably via binding or activation of a factor with minimal impact on local accessibility. In these cases, accessibility appeared to reflect a plastic cell identity encoding the available transcriptional states of the cell where ultimate transcription ensued in response to the cell’s spatial or signaling niche. Future work is warranted to further determine the logic of when cells employ LITE versus NITE mechanisms to regulate distinct cellular processes.

Of note, as with all approaches that model lineage trajectories from single cell data, proper interpretation of the results requires an understanding of the biological system and the underlying model assumptions and limitations^46^. Cells that are transcriptionally and epigenetically similar or following convergent developmental paths are assumed by the model to be nearby within the trajectory, even if they originate from disparate anatomical locations. Experimental approaches that retain the information of cells’ anatomical origins will be important to avoid co-location of cells that appear similar by the multimodal measurements presented to the model although they arise from different locations. Additionally, increased resolution of the data with more cells and more detected genes or accessibility peaks may also reveal previously undetected cell states that will improve the accuracy and resolution of lineage inference. Finally, basic biological knowledge of the system of interest will ensure that the origin point of the trajectory is properly defined so that the directionality reflects the true biological progression through cell states. Future advances in experimental approaches and data resolution will thus further enhance the analyses made possible by MIRA.

In conclusion, MIRA leverages principled probabilistic cell-level topic modeling and gene-level RP modeling to precisely contrast the spatiotemporal dynamics of transcription and local chromatin accessibility at unprecedented resolution. MIRA thereby exposes the key regulators driving fate decisions at lineage branch points and reveals the distinct circuitry regulating fate commitment versus terminal identity. MIRA thus represents a much-needed computational tool for deeply integrated analysis of the rapidly expanding wealth of multimodal data in the single cell field. Moving beyond visualization, MIRA enables rigorous interrogation of the transcriptional and epigenetic mechanisms interacting to drive dynamic biological systems.

## Methods

Complete methods available in Extended Data.

## Acknowledgements

We thank the X. Shirley Liu Lab members, Matthew Oser, and Kai Wucherpfennig for helpful scientific discussions. This work was supported by the National Institutes of Health (NIH) grant U24 CA237617 to CAM. CVT was supported by NIH T32GM007748.

## Author information

AWL developed MIRA, designed analyses, and analyzed the SHARE-seq dataset. CVT co-developed MIRA, designed analyses, and analyzed the 10x dataset. HL and MB contributed to analysis design. XSL and CAM designed analyses and supervised the work. AWL, CVT, XSL, and CAM wrote the manuscript. AWL and CAM originated the work. All authors edited and approved the manuscript.

Competing interests: MB is a consultant to and receives sponsored research support from Novartis. MB serves on the SAB of H3 Biomedicine, Kronos Bio, and GV20 Oncotherapy. XSL is a cofounder, board member, SAB member, and consultant of GV20 Oncotherapy and its subsidiaries; is a stockholder of BMY, TMO, WBA, ABT, ABBV, and JNJ; and received research funding from Takeda, Sanofi, and Novartis.

## Code availability

MIRA is available as a Python package at https://github.com/cistrome/MIRA.

## References

1. Chen, S., Lake, B. B. & Zhang, K. High-throughput sequencing of the transcriptome and chromatin accessibility in the same cell. Nat. Biotechnol. 37, 1452–1457 (2019).

2. Cao, J. et al. Joint profiling of chromatin accessibility and gene expression in thousands of single cells. Science 361, 1380–1385 (2018).

3. Ma, S. et al. Chromatin Potential Identified by Shared Single-Cell Profiling of RNA and Chromatin. Cell 183, 1103–1116.e20 (2020).

4. Zhu, C. et al. An ultra high-throughput method for single-cell joint analysis of open chromatin and transcriptome. Nat. Struct. Mol. Biol. 26, 1063–1070 (2019).

5. Duren, Z., Chen, X., Xin, J., Wang, Y. & Wong, W. H. Time course regulatory analysis based on paired expression and chromatin accessibility data. Genome Res. 30, 622–634 (2020).

6. Gayoso, A. et al. scvi-tools: a library for deep probabilistic analysis of single-cell omics data. bioRxiv 2021.04.28.441833 (2021) doi:10.1101/2021.04.28.441833.

7. Gong, B., Zhou, Y. & Purdom, E. Cobolt: Joint analysis of multimodal single-cell sequencing data. bioRxiv (2021).

8. Minoura, K., Abe, K., Nam, H., Nishikawa, H. & Shimamura, T. scMM: Mixture-of-experts multimodal deep generative model for single-cell multiomics data analysis. bioRxiv 2021.02.18.431907 (2021) doi:10.1101/2021.02.18.431907.

9. Chen, H., Ryu, J., Vinyard, M., Lerer, A. & Pinello, L. SIMBA: SIngle-cell eMBedding Along with features. bioRxiv 2021.10.17.464750 (2021) doi:10.1101/2021.10.17.464750.

10. Lin, Y. et al. scJoint: transfer learning for data integration of single-cell RNA-seq and ATAC-seq. bioRxiv 2020.12.31.424916 (2021) doi:10.1101/2020.12.31.424916.

11. Duren, Z. et al. Integrative analysis of single-cell genomics data by coupled nonnegative matrix factorizations. Proc. Natl. Acad. Sci. U. S. A. 115, 7723–7728 (2018).

12. Lara-Astiaso, D. et al. Chromatin state dynamics during blood formation. Science (2014) doi:10.1126/science.1256271.

13. Rada-Iglesias, A. et al. A unique chromatin signature uncovers early developmental enhancers in humans. Nature 470, 279–283 (2011).

14. 10x Genomics Datasets. https://support.10xgenomics.com/single-cell-multiome-atac-gex/datasets.

15. Blei, D. M. Probabilistic topic models. Commun. ACM 55, 77–84 (2012).

16. Zhao, Y., Cai, H., Zhang, Z., Tang, J. & Li, Y. Learning interpretable cellular and gene signature embeddings from single-cell transcriptomic data. Nat. Commun. 12, 5261 (2021).

17. Bravo González-Blas, C. et al. cisTopic: cis-regulatory topic modeling on single-cell ATAC-seq data. Nat. Methods 16, 397–400 (2019).

18. Kingma, D. P. & Welling, M. Auto-Encoding Variational Bayes. arXiv [stat.ML] (2013).

19. Blei, D. M. Latent Dirichlet Allocation. J. Mach. Learn. Res. 3, 993–1022 (2003).

20. Wang, S. et al. Target analysis by integration of transcriptome and ChIP-seq data with BETA. Nat. Protoc. 8, 2502–2515 (2013).

21. Qin, Q. et al. Lisa: inferring transcriptional regulators through integrative modeling of public chromatin accessibility and ChIP-seq data. Genome Biol. 21, 32 (2020).

22. Schneider, M. R., Schmidt-Ullrich, R. & Paus, R. The hair follicle as a dynamic miniorgan. Curr. Biol. 19, R132–42 (2009).

23. Blanpain, C. & Fuchs, E. Epidermal homeostasis: a balancing act of stem cells in the skin. Nat. Rev. Mol. Cell Biol. 10, 207–217 (2009).

24. L. Byron & M. Wattenberg. Stacked Graphs – Geometry & Aesthetics. IEEE Trans. Vis. Comput. Graph. 14, 1245–1252 (2008).

25. Soma, T., Ogo, M., Suzuki, J., Takahashi, T. & Hibino, T. Analysis of Apoptotic Cell Death in Human Hair Follicles In Vivo andIn Vitro. J. Invest. Dermatol. 111, 948–954 (1998).

26. Cui, C.-Y. et al. Ectodysplasin regulates the lymphotoxin-beta pathway for hair differentiation. Proc. Natl. Acad. Sci. U. S. A. 103, 9142–9147 (2006).

27. Pan, Y. et al. gamma-secretase functions through Notch signaling to maintain skin appendages but is not required for their patterning or initial morphogenesis. Dev. Cell 7, 731–743 (2004).

28. Genander, M. et al. BMP signaling and its pSMAD1/5 target genes differentially regulate hair follicle stem cell lineages. Cell Stem Cell 15, 619–633 (2014).

29. Joost, S. et al. Single-Cell Transcriptomics Reveals that Differentiation and Spatial Signatures Shape Epidermal and Hair Follicle Heterogeneity. Cell Syst 3, 221–237.e9 (2016).

30. Grose, R., Harris, B. S., Cooper, L., Topilko, P. & Martin, P. Immediate early genes krox-24 and krox-20 are rapidly up-regulated after wounding in the embryonic and adult mouse. Dev. Dyn. 223, 371–378 (2002).

31. Hildesheim, J. et al. The hSkn-1a POU transcription factor enhances epidermal stratification by promoting keratinocyte proliferation. J. Cell Sci. 114, 1913–1923 (2001).

32. Zeitvogel, J. et al. GATA3 regulates FLG and FLG2 expression in human primary keratinocytes. Sci. Rep. 7, 1–11 (2017).

33. Hernández-Miranda, L. R., Parnavelas, J. G. & Chiara, F. Molecules and mechanisms involved in the generation and migration of cortical interneurons. ASN Neuro 2, e00031 (2010).

34. La Manno, G. et al. Molecular architecture of the developing mouse brain. Nature 596, 92–96 (2021).

35. Di Bella, D. J. et al. Molecular logic of cellular diversification in the mouse cerebral cortex. Nature 595, 554–559 (2021).

36. Esther, L.-B. et al. Notch Signaling in the Astroglial Phenotype: Relevance to Glutamatergic Transmission. GABA And Glutamate: New Developments In Neurotransmission Research 25 (2018).

37. Yang, N. et al. Generation of pure GABAergic neurons by transcription factor programming. Nat. Methods 14, 621–628 (2017).

38. Raposo, A. A. S. F. et al. Ascl1 Coordinately Regulates Gene Expression and the Chromatin Landscape during Neurogenesis. Cell Rep. 10, 1544–1556 (2015).

39. de Martin, X., Sodaei, R. & Santpere, G. Mechanisms of Binding Specificity among bHLH Transcription Factors. Int. J. Mol. Sci. 22, (2021).

40. Porcher, C., Medina, I. & Gaiarsa, J.-L. Mechanism of BDNF Modulation in GABAergic Synaptic Transmission in Healthy and Disease Brains. Front. Cell. Neurosci. 12, 273 (2018).

41. Mo, J. et al. Early growth response 1 (Egr-1) directly regulates GABAA receptor α2, α4, and θ subunits in the hippocampus. J. Neurochem. 133, 489–500 (2015).

42. Sheng, Z.-H. & Cai, Q. Mitochondrial transport in neurons: impact on synaptic homeostasis and neurodegeneration. Nat. Rev. Neurosci. 13, 77–93 (2012).

43. Harrington, A. J. et al. MEF2C regulates cortical inhibitory and excitatory synapses and behaviors relevant to neurodevelopmental disorders. Elife 5, (2016).

44. Park, N. I. et al. ASCL1 Reorganizes Chromatin to Direct Neuronal Fate and Suppress Tumorigenicity of Glioblastoma Stem Cells. Cell Stem Cell 21, 411 (2017).

45. Chen, C.-H. et al. Determinants of transcription factor regulatory range. Nat. Commun. 11, 2472 (2020).

46. Tritschler, S. et al. Concepts and limitations for learning developmental trajectories from single cell genomics. Development 146, (2019).

47. 10x Genomics Datasets. https://www.10xgenomics.com/resources/datasets/pbmc-from-a-healthy-donor-granulocytes-removed-through-cell-sorting-10-k-1-standard-2-0-0.

48. Saelens, W., Cannoodt, R., Todorov, H. & Saeys, Y. A comparison of single-cell trajectory inference methods. Nat. Biotechnol. 37, 547–554 (2019).

