## Extended Data and Methods for "MIRA: Joint regulatory modeling of multimodal expression and chromatin accessibility in single cells"

**Extended Data Table 1.** Gene set enrichments of each MIRA expression topic in the hair follicle dataset.

**Extended Data Table 2.** Motif enrichments of each MIRA accessibility topic in the hair follicle dataset.

**Extended Data Table 3.** Gene set enrichments of each MIRA expression topic in the IFE dataset.

**Extended Data Table 4.** Motif enrichments of each MIRA accessibility topic in the IFE dataset.

**Extended Data Table 5.** Gene set enrichments of each MIRA expression topic in the embryonic brain dataset.

**Extended Data Table 6.** Motif enrichments of each MIRA accessibility topic in the embryonic brain dataset.

The above Extended Data Tables are available at the following link: https://github.com/AllenWLynch/MIRA_supplementary_tables

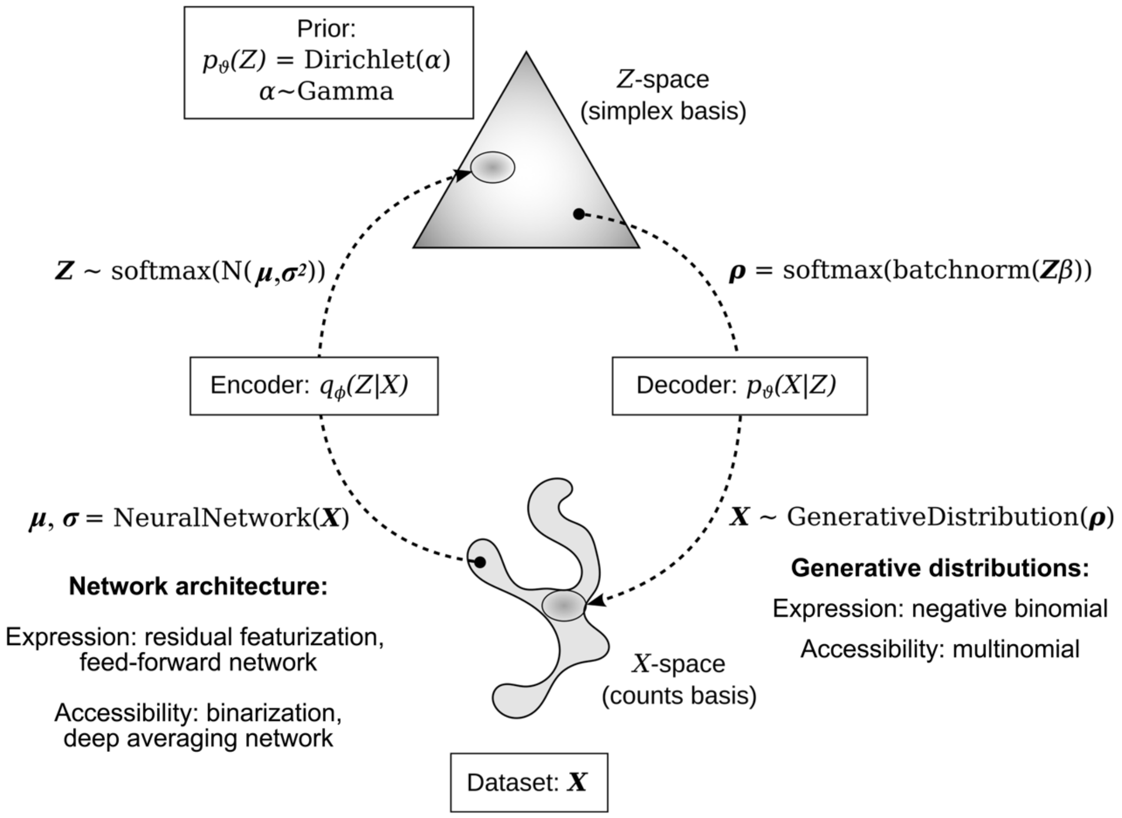


**Extended Data Fig. 1 | Overview of MIRA topic model architecture.** The MIRA topic model uses a variational autoencoder (VAE) approach to learn stochastic mappings between observations in $\boldsymbol{X}$-space, gene-counts or peak-counts in a cell, which are high-dimensional and noisy, and a simpler latent $\boldsymbol{Z}$-space or topic space, which exists on the simplex basis with a Dirichlet prior. (*bottom right*) The generative model relates the observations $\boldsymbol{X}$ to the estimated composition $\boldsymbol{\rho}$ over features (genes or peaks), sampling a negative binomial distribution for RNA counts and a multinomial distribution for ATAC peaks. (*top right*) The composition over features is given by the topic matrix $\boldsymbol{\beta}$ encoding topic-feature associations and the latent topics $\boldsymbol{Z}$ of a cell, which are sampled from the distribution $\boldsymbol{q_\phi(Z|X)}$, the variational approximation of $\boldsymbol{p_\vartheta(Z|X)}$. (*top* left) The distribution of $\boldsymbol{Z}$ is parameterized by $\boldsymbol{\mu}$ and $\boldsymbol{\sigma}^2$, outputs from the encoder neural network given the $\boldsymbol{X}$-space observations as inputs. (*bottom left*) The encoder neural network for RNA data performs deviance residual featurization of counts which are passed through feed-forward layers. The ATAC data encoder passes binarized peak accessibility features through a deep averaging network. (Illustration adapted from Kingma and Welling, *Foundations and Trends in Machine Learning*, 2019).

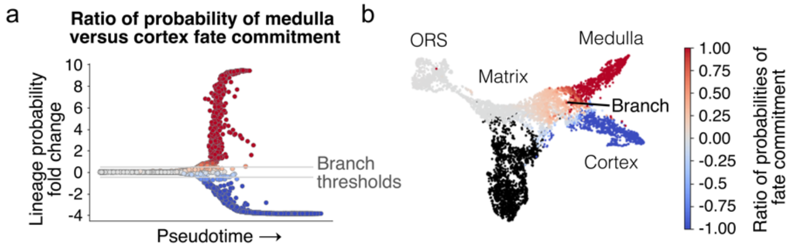


**Extended Data Fig. 2 | MIRA defines branch points between lineages where fate commitment probabilities diverge. a,** Ratio of probability of medulla fate commitment versus cortex commitment of each cell in the hair follicle, arranged by pseudotime. MIRA defines branch points between lineages where probabilities of differentiating into one terminal state diverges from another. **b,** MIRA joint representation UMAP colored by ratio of probability of medulla fate commitment within the ORS, matrix, medulla, and cortex populations. Differentiation in the hair follicle proceeds from ORS to progenitor matrix cells, which then specify into the medulla or cortex fate. (IRS cells indicated in black are not included in this trajectory).

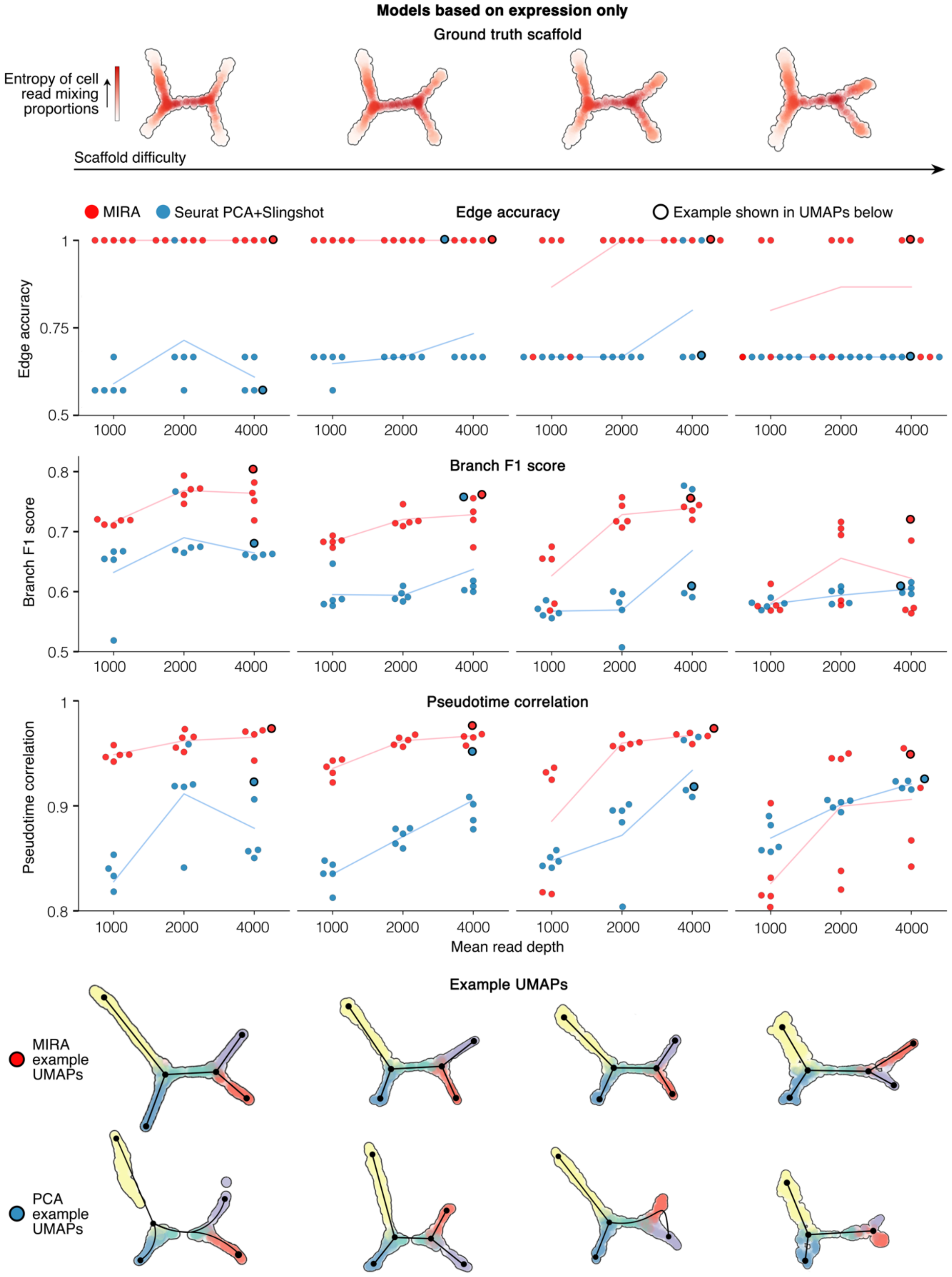


**Extended Data Fig. 3 | MIRA outperforms standard methodology for resolving lineage trajectories using expression data alone.** Benchmarking results comparing MIRA to standard methodology of Seurat PCA+Slingshot in the indicated metrics of lineage trajectory inference using expression data alone. Top row shows ground truth scaffolds, which are computationally synthesized by mixing reads from distinct populations of single cells from a 10x Genomics dataset[1] of peripheral blood mononuclear cells (PBMCs). Scaffold difficulty increases from left to right, where more difficult scaffolds contain lineages where mixture components are more similar. Line plots indicate MIRA (red) versus Seurat PCA+Slingshot (blue) performance in each of the four scaffold difficulties with trials for three different mean read depths (lower read depth further increases the difficulty of solving the topology). For each trial, 5 replicates were tested for each modeling approach. *Edge accuracy* measures the accuracy of the inferred edges compared to ground truth (dynverse's edge flip score[2]). *Branch F1 score*[2] measures the precision and recall of the inferred branches compared to ground truth. *Pseudotime correlation*[2] measures the correlation between inferred versus ground truth pseudotime for each cell. The bottom rows show example UMAPs for MIRA or Seurat PCA+Slingshot for each scaffold difficulty with black edges showing lineage parsing from each algorithm. Cells colored by ground truth branch assignment where blue cells are the origin state. In the line plots above, black outlines indicate the points for the models shown in the example UMAPs.

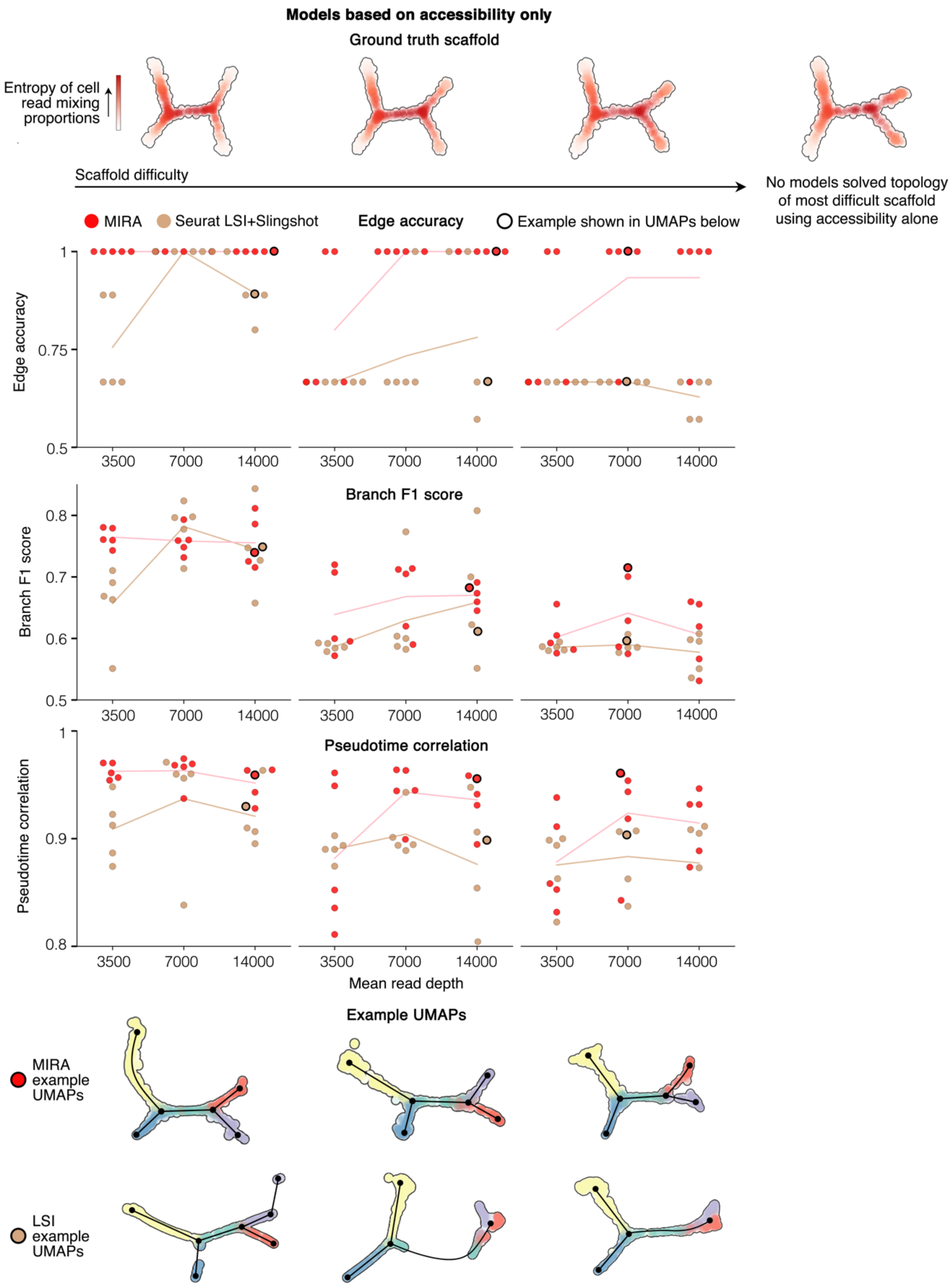


**Extended Data Fig. 4 | MIRA outperforms standard methodology for resolving lineage trajectories using accessibility data alone.** Benchmarking results comparing MIRA to standard methodology of Seurat LSI+Slingshot in the indicated metrics of lineage trajectory inference using accessibility data alone. Top row shows ground truth scaffolds with scaffold difficulty increasing from left to right. No models solved the topology of the most difficult scaffold using accessibility alone so metric comparisons are shown for the other three scaffolds. See Extended Data Fig. 3 for description of metrics.

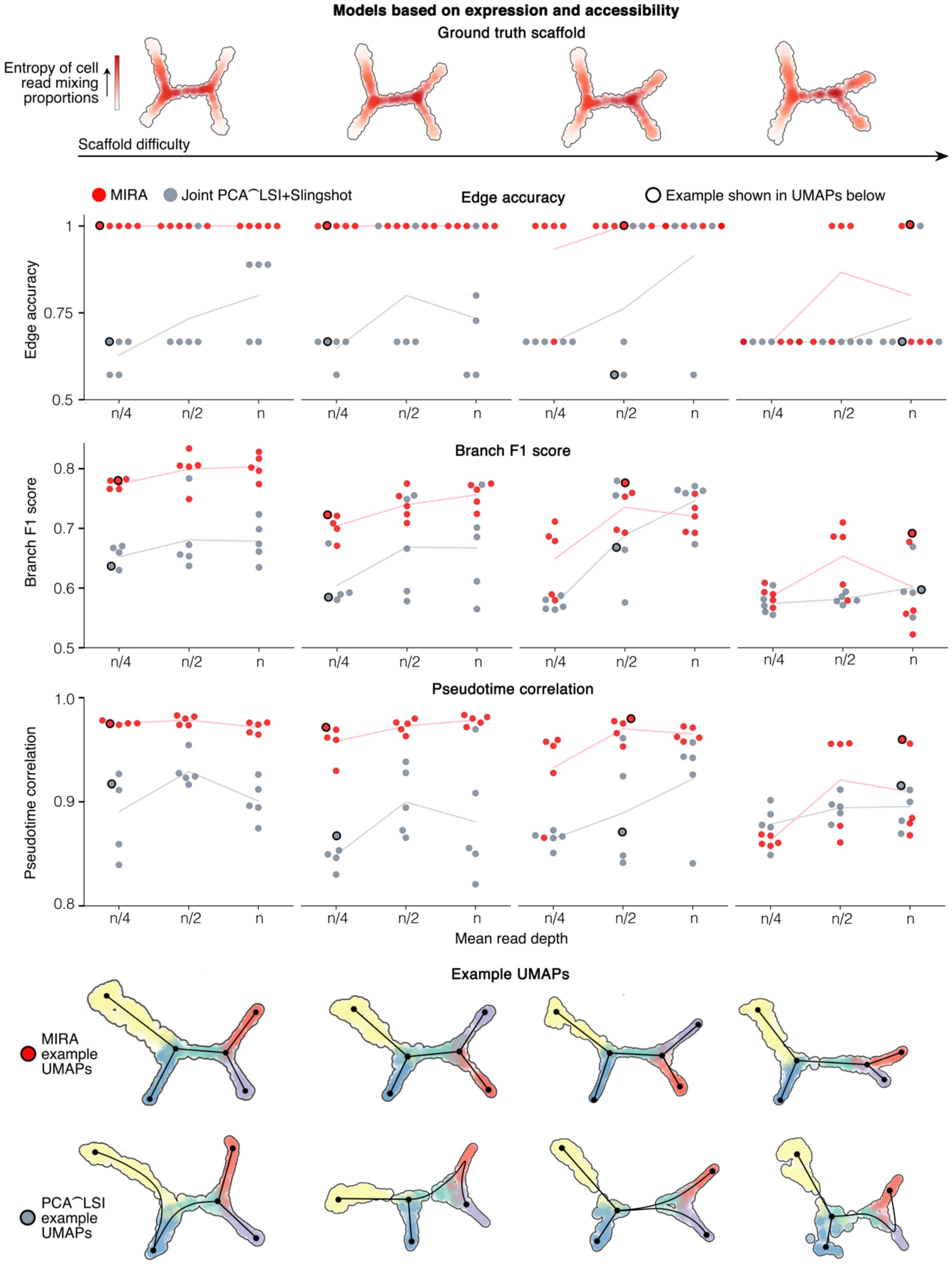


**Extended Data Fig. 5 | MIRA outperforms standard methodology for resolving lineage trajectories using both expression and accessibility data jointly.** Benchmarking results comparing MIRA joint representation to standard methodology of joint representation combining Seurat PCA of expression data and Seurat LSI of accessibility data followed by Slingshot. See Extended Data Fig. 3 for description of metrics. For expression data, mean read depth n=4000; for accessibility data, mean read depth n=14000.

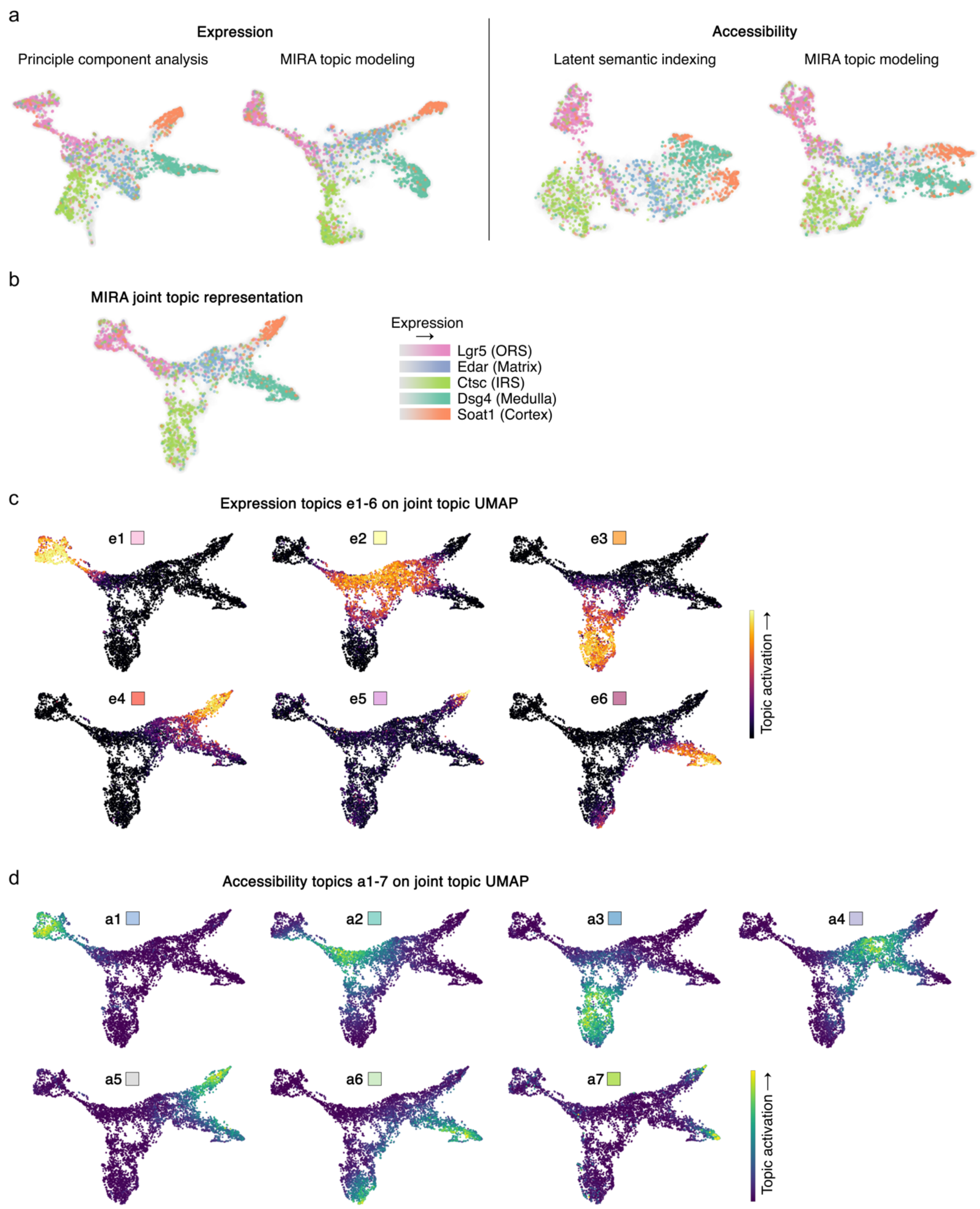


**Extended Data Fig. 6 | MIRA topics describing hair follicle cells were sparse and nonredundant. a,** UMAP based on standard methodology versus MIRA topic modeling for expression or accessibility. Standard PCA-based representation of expression shows matrix population as shifted away from its predecessor ORS and descendant IRS, medulla, and cortex cells. However, MIRA topic modeling of expression appropriately represents matrix cells as an intermediate population between the aforementioned lineages. Standard LSI-based representation of accessibility shows ORS cells interjected between matrix and its descendant IRS and shows medulla situated between two separate cortex populations. Conversely, MIRA topic modeling of accessibility appropriately represents matrix cells as continuous with its descendant IRS and better separates medulla and cortex into two distinct branches. **b,** MIRA joint topic representation of expression and accessibility. In (a-b), colors demonstrate expression of marker genes of indicated lineages**. c,** MIRA expression topics e1-6 and **d,** MIRA accessibility topics a1-7 on joint representation UMAP. In (c-d), colored boxes correspond to topic colors as on stream graphs in Fig. 2c and Extended Data Fig. 7a.

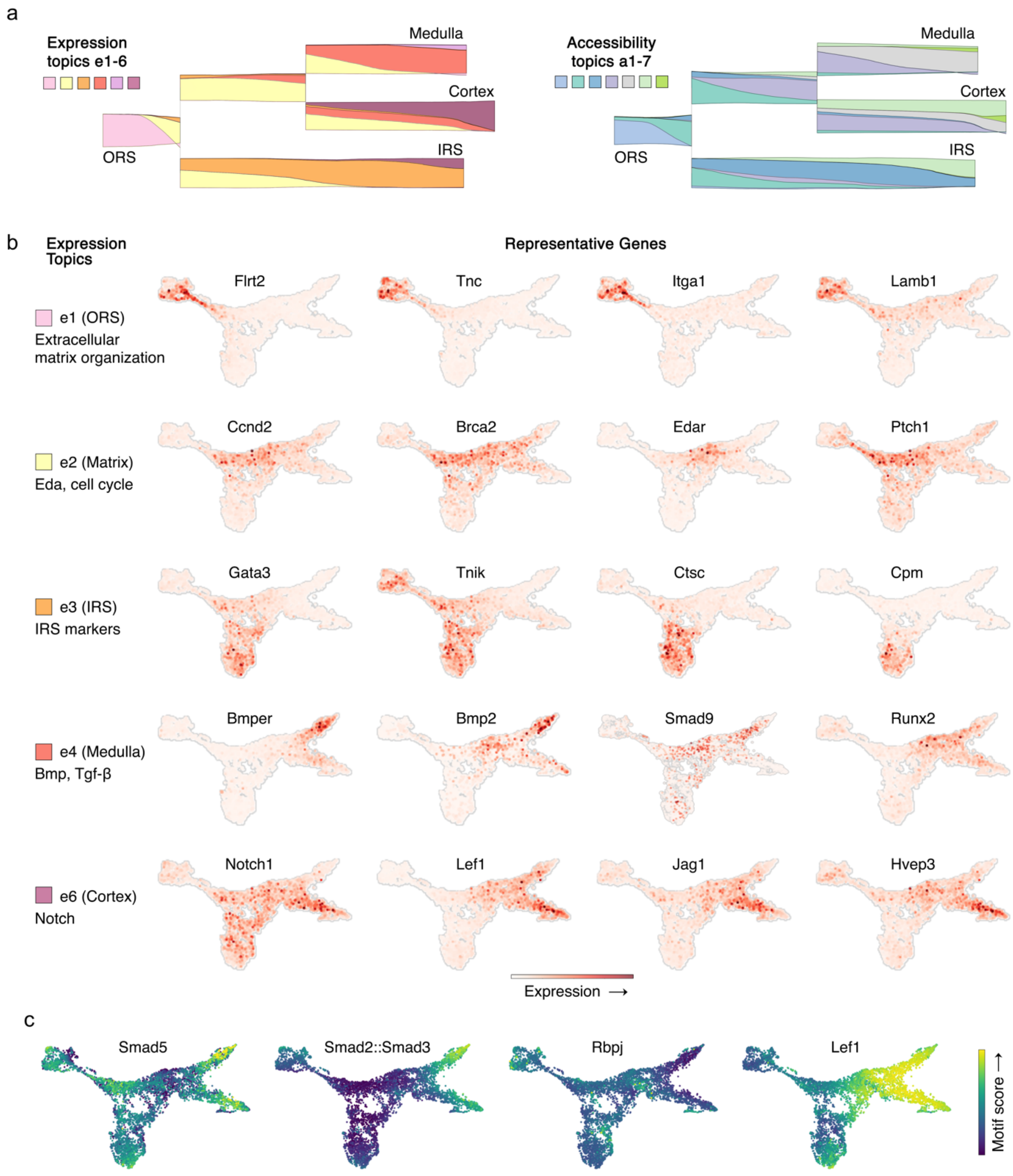


**Extended Data Fig. 7 | MIRA topics described gene modules activated in each lineage. a,** Stream graph of window-averaged cell-topic compositions starting from ORS cell state, progressing rightward through pseudotime (to facilitate visualization of all lineages concurrently, pseudotime scale is not log-transformed, unlike other presented stream graphs). **b,** MIRA joint topic representation colored by expression of genes highly activated in each of the indicated topics, which described the activated gene modules in each lineage. **c,** MIRA joint topic representation colored by indicated motif scores.

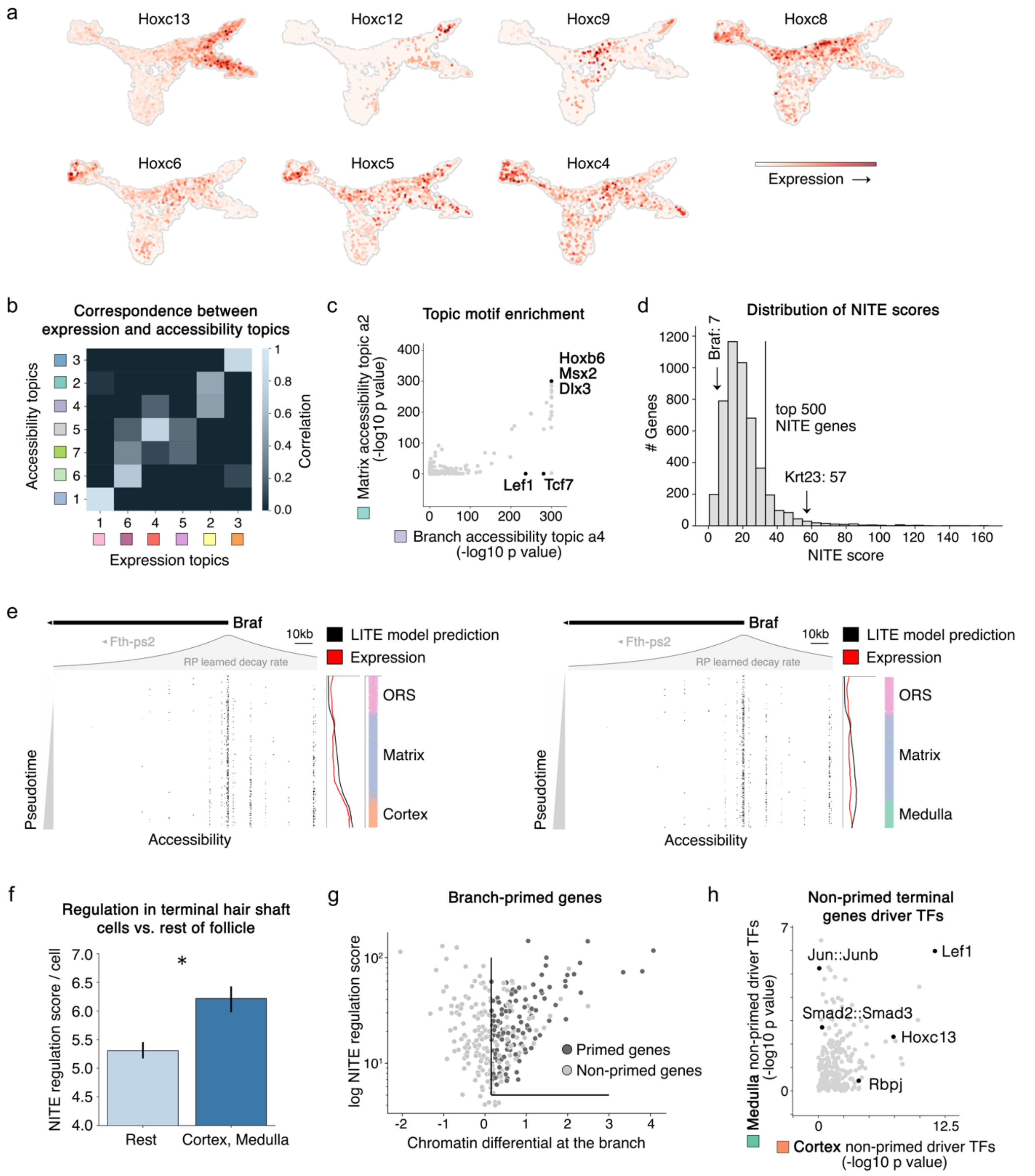


**Extended Data Fig. 8 | Terminal medulla and cortex cells showed significantly higher NITE regulation compared to cells earlier in hair follicle differentiation. a,** MIRA joint topic representation colored by expression of Hoxc genes, indicating that Hoxc motifs activated in both the medulla and cortex accessibility topics (a5 and a6, respectively) were most attributable to Hoxc13 based on its expression in these lineages**. b,** Correlation matrix between expression and accessibility topics. While some topics had a clear one-to-one correlation between modalities (e.g. expression topic e1 with accessibility topic a1), others did not strongly correlate with a single topic from the opposing modality (e.g. branch accessibility topic a4). **c,** Comparison of motif enrichment in top peaks of preceding matrix versus subsequent branch accessibility topics (a2 and a4, respectively). While most motifs were shared between these topics, accessibility of Wnt signaling-related motifs uniquely arose at the branch. **d,** Distribution of NITE scores among genes expressed in the hair follicle. Scores of example LITE gene Braf and NITE gene Krt23 are indicated by arrows. **e,** LITE gene Braf as shown in Fig. 3c but extended to include further downstream region. As described in Fig. 3c, plot shows chromatin accessibility fragments across pseudotime (moving downwards) in trajectories from ORS to matrix to cortex or medulla. Colored bars on the right indicate the identity of cells (colored by clusters in Fig. 2a) within each bin reflected by each row of accessibility fragments. Line plots across pseudotime depict the indicated gene's observed expression (red) and LITE model prediction of expression (black), which is informed by the local accessibility reflected in the fragment plot. **f,** Medulla and cortex cells showed significantly more NITE regulation than other cells in the hair follicle (*$p<0.05$, Wilcoxon rank-sum, error bars=standard deviation). **g,** Genes ultimately expressed in medulla or cortex that were primed at the branch were defined as those with a NITE regulation score above the indicated thresholds that had positive chromatin differential at the branch, indicating that expression was overestimated based on local chromatin accessibility. Branch-primed genes must also be upregulated in the downstream lineage relative to matrix cells. **h,** Driver transcription factor analysis of non-primed medulla versus cortex genes.

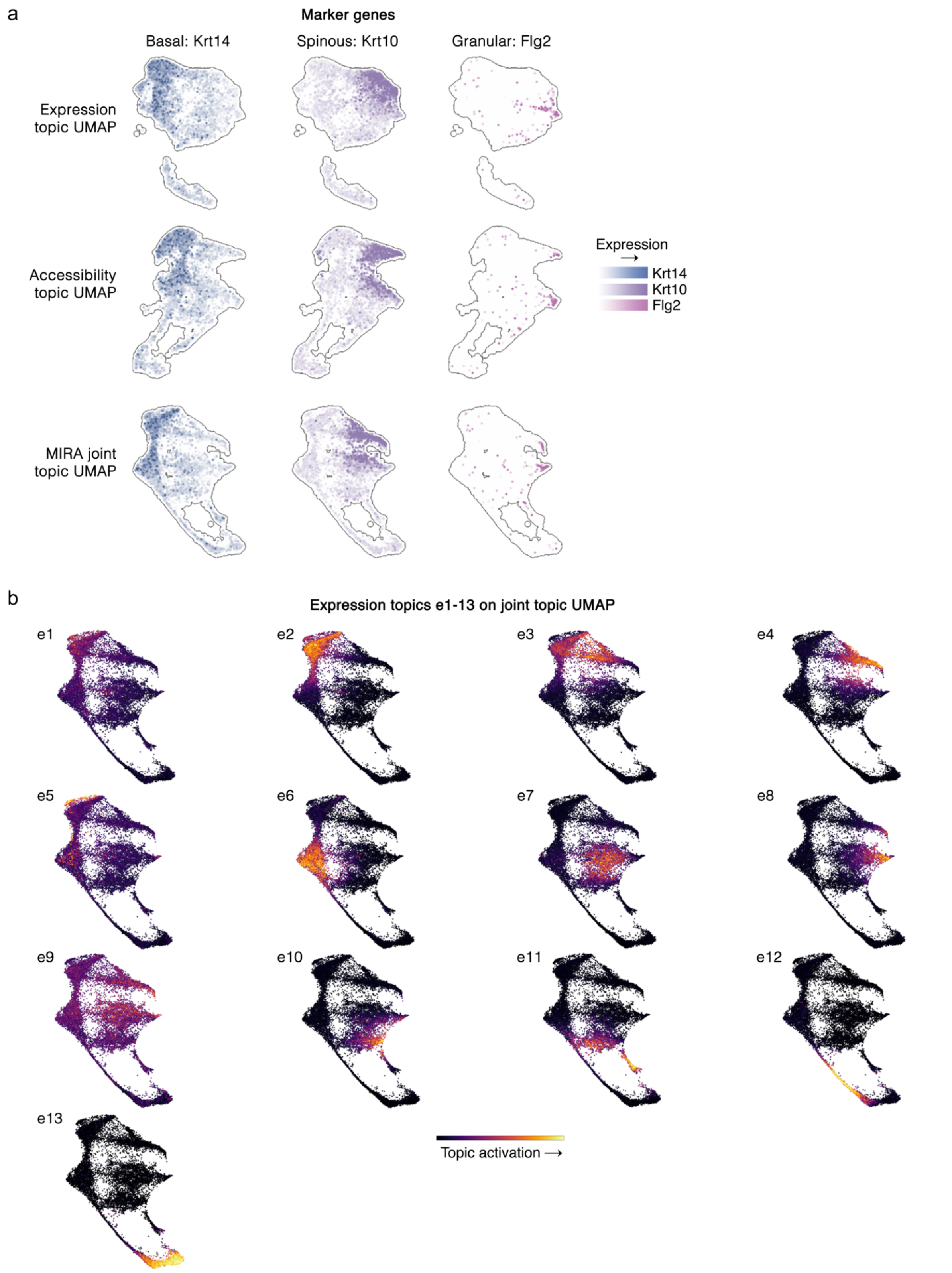


**Extended Data Fig. 9 | MIRA expression topics describing IFE cells captured shared and lineage-specific states. a,** Expression of marker genes of indicated lineages on MIRA expression, accessibility, and joint topic UMAPs. **b,** MIRA expression topics e1-13 on joint representation UMAP.

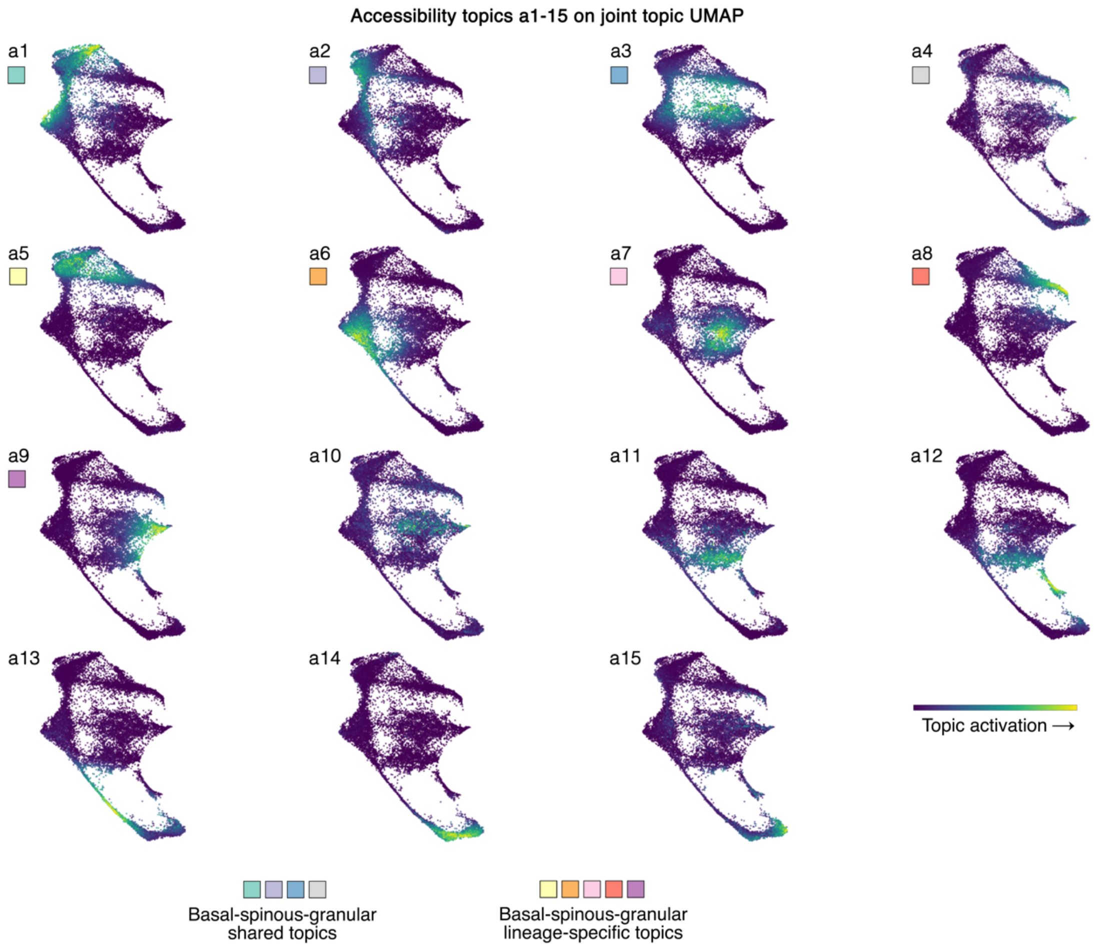


**Extended Data Fig. 10 | MIRA accessibility topics describing IFE cells captured shared and lineage-specific states.** MIRA accessibility topics a1-15 on joint representation UMAP. Colored boxes correspond to topics indicated in Fig. 5h, which are shared or lineage-specific within the basal-spinous-granular or intermediate basal-spinous-granular differentiation trajectories as annotated in Fig. 5a-b.

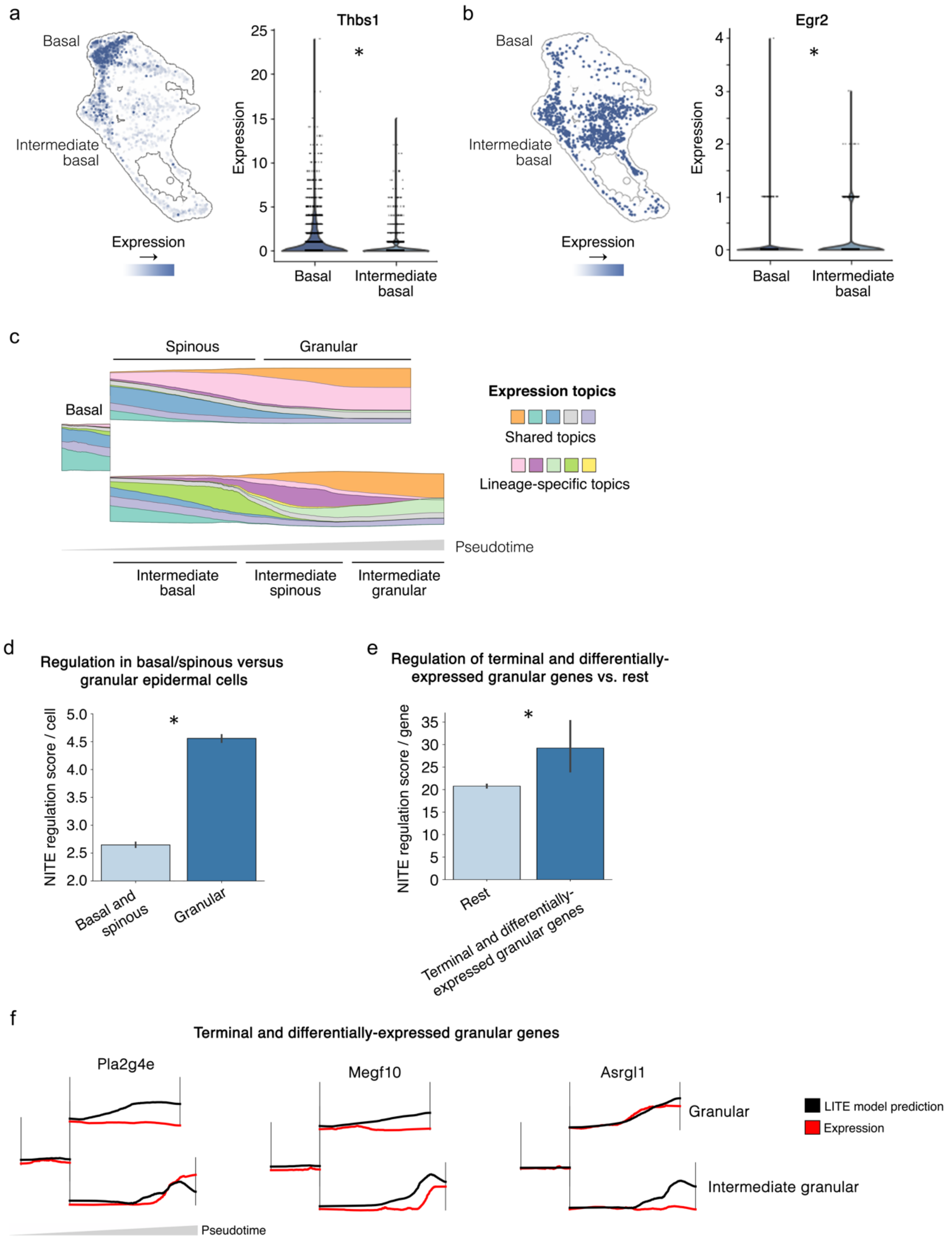


**Extended Data Fig. 11 | Terminal granular cells were enriched for NITE regulation. a,** *Thbs1* and **b,** *Egr2* expression distinguished basal cells distant from the hair follicle from those within the intermediate basal-spinous-granular trajectory near the hair follicle (*p<0.05, Wilcoxon rank-sum, Benjamini-Hochberg corrected). **c,** Stream graph of expression topic compositions of basal-spinous-granular *(top)* and intermediate basal-spinous-granular *(bottom)* lineages. **d,** Terminal IFE granular cells showed significantly more NITE regulation than cells earlier in the differentiation trajectory (basal and spinous cells) (*p<0.05, Wilcoxon rank-sum, error bars=standard deviation). **e,** Genes upregulated in granular cells that were differentially-expressed between granular populations had significantly higher NITE scores than other genes (*p<0.05, Wilcoxon rank-sum, error bars=standard deviation). **f,** Examples of terminally upregulated, differentially-expressed granular genes' local chromatin accessibility (LITE model prediction) and expression. Despite accessibility increasing in both lineages, expression only increased in one lineage.

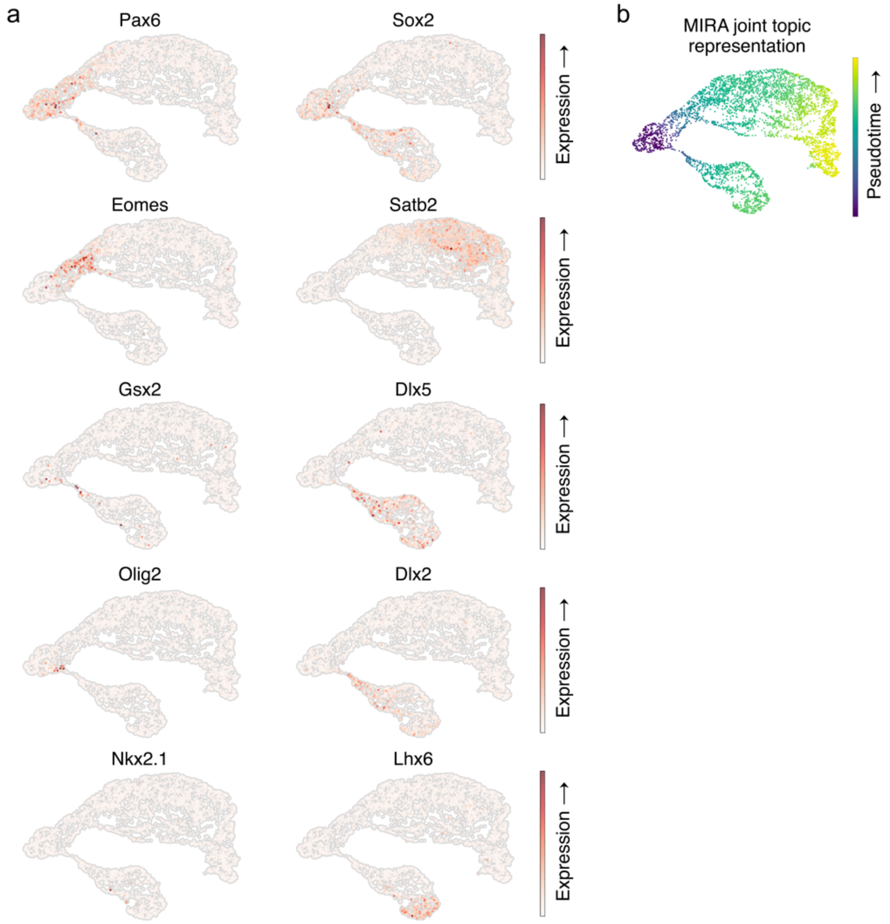


**Extended Data Fig. 12 | MIRA joint representation captured temporal progression of lineage trajectories in embryonic brain development. a,** MIRA topic modeling constructed a joint representation with Pax6+/Sox2+ radial glia-like cells located centrally between the astrocyte, excitatory neuron, and inhibitory neuron branches. Pax6 marks both dorsal progenitors that give rise to astrocytes and excitatory neurons and the anatomically juxtaposed ventral progenitors in the lateral ganglionic eminence that give rise to inhibitory neurons[3,4]. Despite the anatomic separation of excitatory and inhibitory progenitors, they share a common transcriptional state[4], which resulted in their co-location within the UMAP embedding. MIRA's joint representation appeared to appropriately capture the temporal progression towards excitatory neurons including by the activation of Eomes followed by Satb2 and towards inhibitory neurons by the expression of Gsx2 followed by Dlx5, consistent with prior studies[4]. The inhibitory progenitors in the 10x Genomics dataset likely reflect those within the lateral ganglionic eminence given their expression of Pax6, Dlx2, Gsx2, and Olig2 without concurrent Lhx6 or Nkx2.1 expression[3]. The 10x Genomics dataset did not appear to include inhibitory progenitors from the medial (Dlx2+/Gsx2+/Olig2+/Lhx6+/Nkx2.1+) and caudal (Pax6+/Dlx2+/Gsx2+/Olig2+/Lhx6+) ganglionic eminences, which are anatomically further from the dorsal region and less likely to have been collected alongside the excitatory progenitors[3]. **b,** MIRA joint expression and accessibility topic UMAP, colored by pseudotime of inferred trajectory.

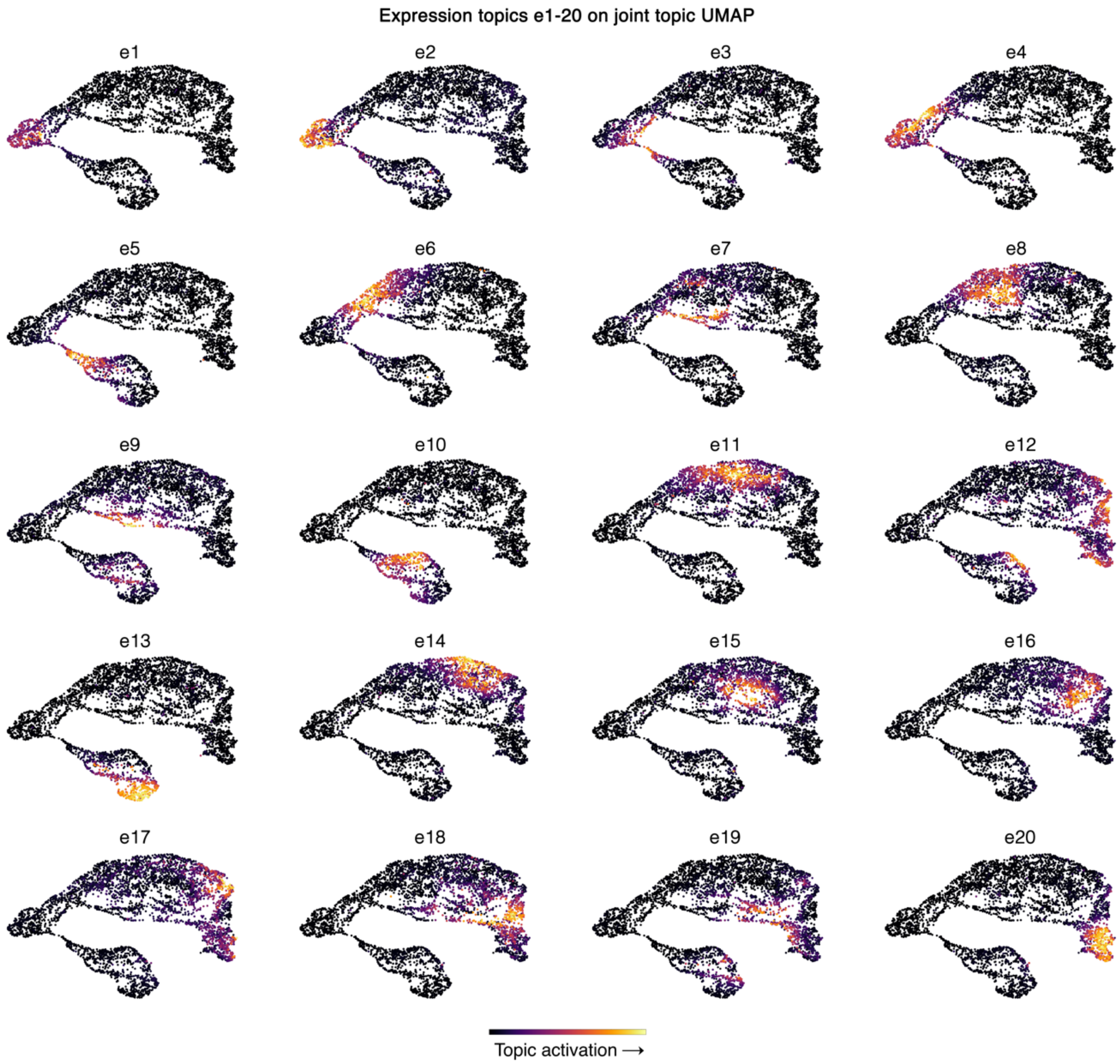


**Extended Data Fig. 13 | MIRA expression topics describing embryonic brain cells were sparse and nonredundant.** MIRA expression topics e1-20 on joint representation UMAP.

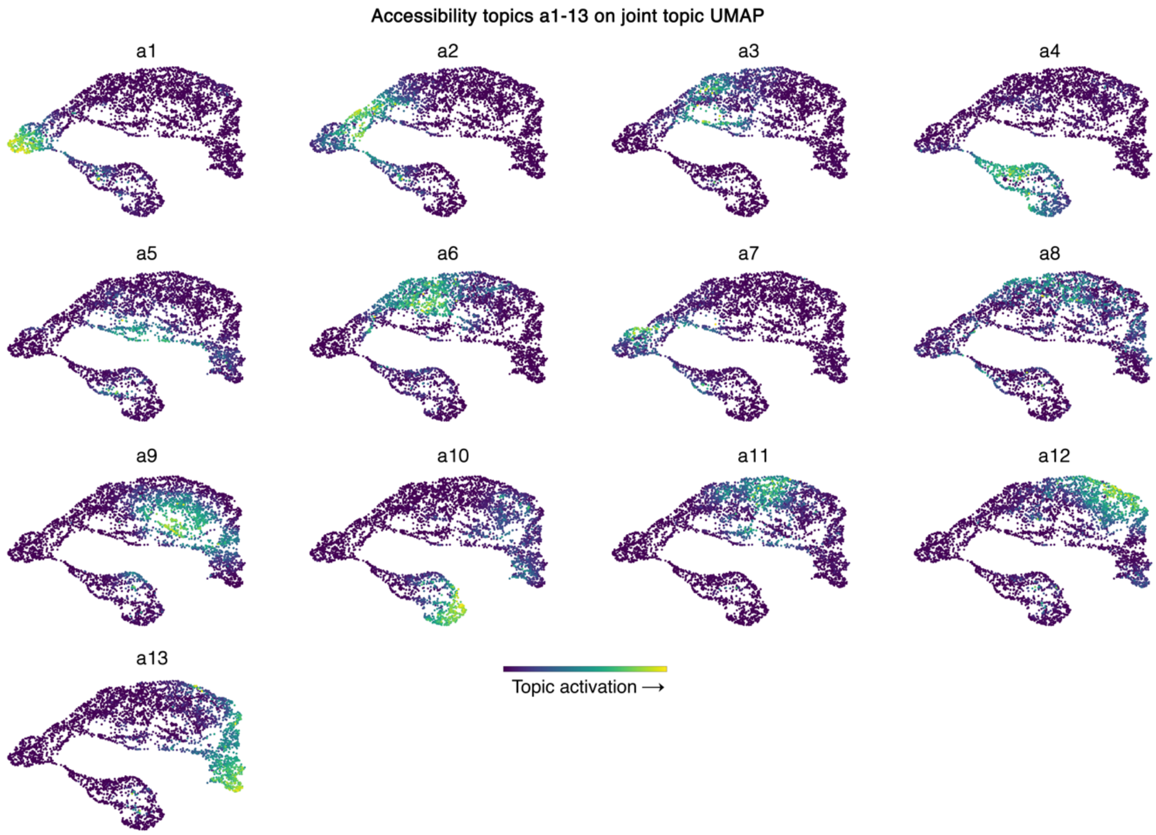


**Extended Data Fig. 14 | MIRA accessibility topics describing embryonic brain cells were sparse and nonredundant.** MIRA accessibility topics a1-13 on joint representation UMAP.

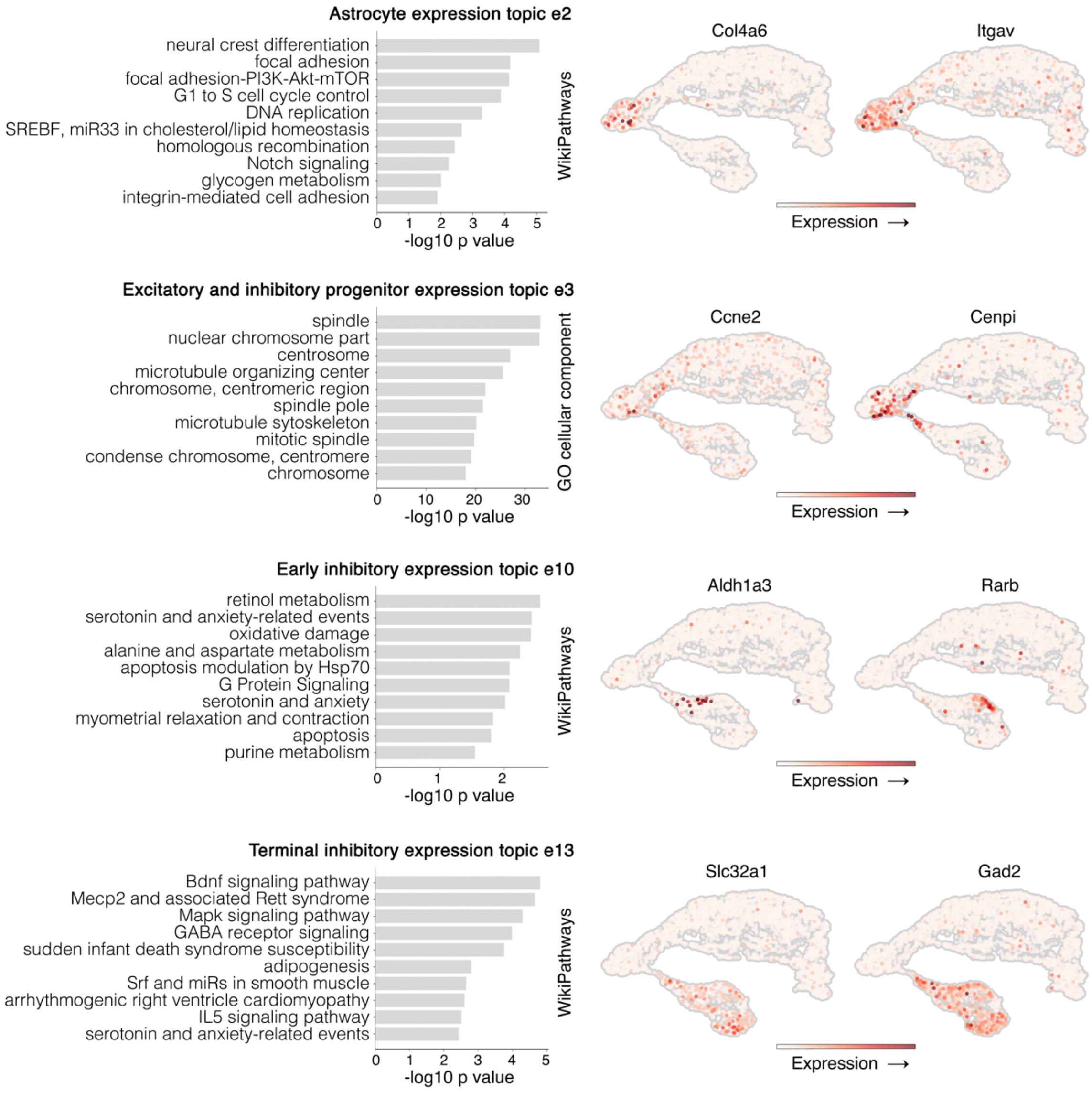


**Extended Data Fig. 15 | MIRA expression topics illuminated pathways key to each stage of differentiation in astrocytes, excitatory and inhibitory progenitors, and inhibitory neurons.** Pathway enrichment of MIRA expression topics describing astrocytes (focal adhesion), excitatory and inhibitory mixed progenitors (proliferation), early inhibitory neurons (retinoic acid signaling), and terminal inhibitory neurons (Bdnf signaling, GABA receptors) (expression of example genes shown to the right).

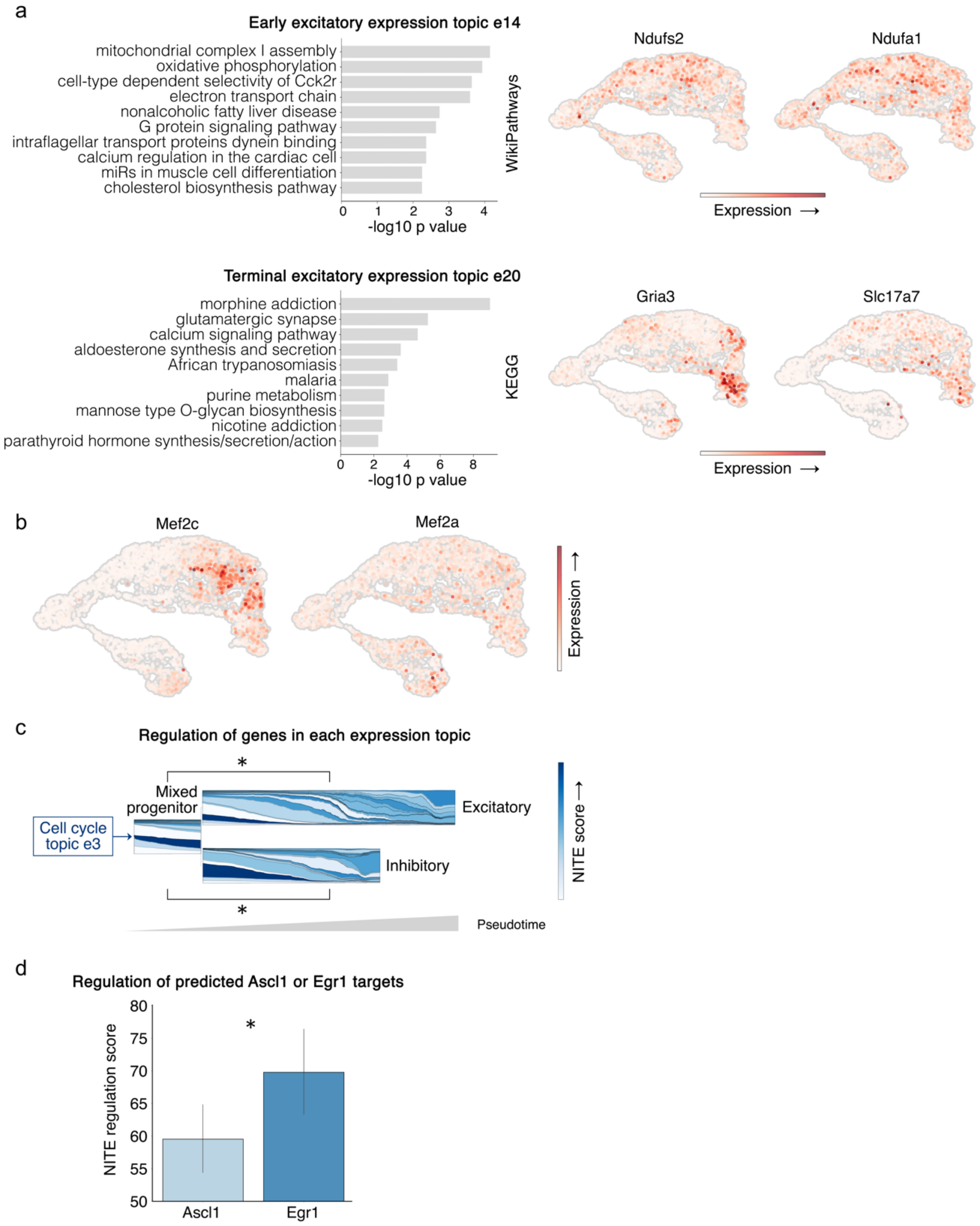


**Extended Data Fig. 16 | MIRA expression topics illuminated pathways key to each stage of differentiation in excitatory neurons. a,** Pathway enrichment of MIRA expression topics describing early excitatory (mitochondrial machinery) or terminal excitatory neurons (glutamatergic neurotransmission) (expression of example genes shown to the right). **b,** *Mef2c* was more highly expressed in excitatory neurons, indicating that Mef2 motifs enriched in the terminal excitatory neuron topic were likely attributable to Mef2c. **c,** Stream graphs of expression topics across pseudolineage trajectory colored by NITE versus LITE regulation of the top genes in each topic. Topics describing earlier states tended towards LITE regulation with the notable exception of topic e3, which is composed of cell cycle genes that have been previously described to be regulated with minimal influence of local chromatin accessibility state[5]. Topics describing terminal states tended more towards NITE regulation, including the major terminal excitatory and inhibitory neuron topics that are composed of neurotransmitter genes. Overall, expression topics describing the excitatory and inhibitory progenitor states (labeled mixed progenitor) were significantly enriched for LITE regulation, whereas after commitment to either the excitatory or inhibitory fate, topics were significantly enriched for NITE regulation (*p<0.05, Wilcoxon rank-sum, Benjamini-Hochberg corrected). **d,** Genes predicted by MIRA pISD modeling to be regulated by pioneer transcription factor Ascl1 showed significantly more LITE regulation compared to genes predicted to be regulated by non-pioneer-like Egr1 (*p<0.05, Wilcoxon rank-sum, error bars=standard deviation).

### References


1. 10x Genomics Datasets. https://www.10xgenomics.com/resources/datasets/pbmc-from-a-healthy-donor-granulocytes-removed-through-cell-sorting-10-k-1-standard-2-0-0.
2. Saelens, W., Cannoodt, R., Todorov, H. & Saeys, Y. A comparison of single-cell trajectory inference methods. *Nat. Biotechnol.* **37**, 547–554 (2019).
3. Hernández-Miranda, L. R., Parnavelas, J. G. & Chiara, F. Molecules and mechanisms involved in the generation and migration of cortical interneurons. *ASN Neuro* **2**, e00031 (2010).
4. La Manno, G. *et al.* Molecular architecture of the developing mouse brain. *Nature* **596**, 92–96 (2021).
5. Ma, S. *et al.* Chromatin Potential Identified by Shared Single-Cell Profiling of RNA and Chromatin. *Cell* **183**, 1103–1116.e20 (2020).

### Extended Methods

#### MIRA input data

The input data for MIRA is expression (raw gene count) and accessibility (binary peak count) matrices from multimodal RNA-sequencing (scRNA-seq) and Assay for Transposase-Accessible Chromatin-sequencing (scATAC-seq) in the same single cells.

#### MIRA topic model

*Model architecture*

The MIRA topic model is a generative probabilistic model where the cell's observed features (transcript counts or accessible genomic intervals) are explained by hidden latent variables. Inspired by topic modeling methods such as Latent Dirichlet Allocation (LDA)[1], we assume that the latent variables describing a cell's state are sparse and compositional. As such, only a few latent variables are active at a time to define each state, and that the latent variables relate linearly to changes in the cell's observed attributes. This constrains the model such that the latent variables decompose expression and accessibility into coherent, interpretable patterns of covarying features. Each latent variable thereby describes a "topic" of coregulated genes or co-accessible genomic loci and suggests that the genes and loci influenced by that topic share some underlying facet of regulation.

MIRA uses a variational autoencoding neural network[2] (Extended Data Fig. 1) to discover latent topics from expression or accessibility data, which enables faster and more flexible inference than classic Gibbs sampling-based solutions like LDA[3]. The central part of the model is the same for expression and accessibility data, although the numbers of topics may differ. The input data is either a gene counts matrix for expression data or a binary peak-count matrix for accessibility data. From either of these inputs, the topic model learns a latent representation for cells $Z \in \mathrm{I}^{N_{\text{cells}} \times N_{\text{topics}}}$ where $\mathrm{I}$ is the unit interval [0,1] and where:

$$\sum_{t=1}^{N_{\text{topics}}} Z_{it} = 1, \forall\, i \in \{1, \dots, N_{\text{cells}}\}$$

For matrices $M_{xy}$, let the notation $M_{x\cdot}$ indicate the matrix row indexed by $x$ and $M_{\cdot y}$ indicate the matrix column indexed by $y$. We specify a sampling procedure such that $Z_{i\cdot} \in \mathrm{I}^{N_{\text{topics}}}$ is Dirichlet-distributed with a hierarchical prior controlling the pseudocounts allotted to each topic:

$$Z_{i\cdot} \sim \text{Dirichlet}\left(\alpha_1, \dots, \alpha_{N_{\text{topics}}}\right), \forall\, i \in \{1, \dots, N_{\text{cells}}\}$$

$$\alpha_t \sim \text{Gamma}\left(2, \tfrac{2N_{\text{topics}}}{\mathcal{J}}\right), \forall\, t \in \{1, \dots, N_{\text{topics}}\}$$

where $\mathcal{J}$ is the initial pseudocount allotted to the Dirichlet distribution, and $\boldsymbol{\alpha}$ is the random variable controlling the sparsity of $Z$. The gamma hyperprior, parameterized as $(\text{shape}, \text{rate})$, controls the sparsity of each topic, allowing for the data-driven tuning of sparsity to fit different patterns and modalities. Density of the gamma hyperprior is concentrated below the mean at $\mathcal{I}/N_{\text{topics}}$, prioritizing the capture of very sparse topics but still enabling flexibility.

The latent representation of each cell represents the composition of topics that describe the expression or accessibility observations measured from that cell. We adapt the generative process of the model to account for the distinct statistical properties of each modality[4,5]. We denote the gene expression data matrix as $X^{\text{RNA}} \in \mathbb{Z}_{\geq 0}^{N_{\text{cells}} \times N_{\text{genes}}}$, and specify a model such that each observation is independently drawn from the following generative process:

$$X_{ij}^{\text{RNA}} \sim \text{NegativeBinomial}(\, n_i \rho_{ij}, \theta_j \,), \forall\, i \in \{1, \dots, N_{\text{cells}}\}, \forall\, j \in \{1, \dots, N_{\text{genes}}\}$$

$$\rho_{i\cdot} = \text{softmax}\big(\, \text{batchnorm}(\, Z_{i\cdot}\beta \,)\big), \forall\, i \in \{1, \dots, N_{\text{cells}}\}$$

$$n_i \sim \text{LogNormal}\big(\log\, \hat{n}_i^{\text{RNA}}, 1\big), \forall\, i \in \{1, \dots, N_{\text{cells}}\}$$

$$\hat{n}_i^{\text{RNA}} = \textstyle\sum_{j=1}^{N_{\text{genes}}} X_{ij}^{\text{RNA}}, \forall\, i \in \{1, \dots, N_{\text{cells}}\}$$

where $\rho \in \mathrm{I}^{N_{\text{cells}} \times N_{\text{genes}}}$ is the predicted composition of expression across all genes in each cell and $\sum_{j=1}^{N_{\text{genes}}} \rho_{ij} = 1, \forall\, i \in \{1, \dots, N_{\text{cells}}\}$; $\beta$ is the $\mathbb{R}^{N_{\text{topics}} \times N_{\text{genes}}}$ matrix linking gene expression to the influence of topics; and $n_i$ is the effective read depth of cell $i$. $\boldsymbol{\theta} \in \mathbb{R}_{\geq 0}^{N_{\text{genes}}}$ is a global variable determining the overdispersion of the negative binomial distribution for each gene across all cells.

For chromatin accessibility data, we model observations of accessibility $X^{\text{ATAC}} \in \{0,1\}^{N_{\text{cells}} \times N_{\text{peaks}}}$ across all regions given a cell using the multinomial distribution:

$$X_{i\cdot}^{\text{ATAC}} \sim \text{Multinomial}\big(\, \rho_{i\cdot}, \hat{n}_i^{\text{ATAC}}\big), \forall\, i \in \{1, \dots, N_{\text{cells}}\}$$

$$\rho_{i\cdot} = \text{softmax}\big(\, \text{batchnorm}(\, Z_{i\cdot}\beta \,)\big), \forall\, i \in \{1, \dots, N_{\text{cells}}\}$$

$$\hat{n}_i^{\text{ATAC}} = \sum_{k=1}^{N_{\text{peaks}}} X_{ik}^{\text{ATAC}}, \forall\, i \in \{1, \dots, N_{\text{cells}}\}$$

where $\rho \in \mathrm{I}^{N_{\text{cells}} \times N_{\text{peaks}}}$ is the predicted composition of accessibility across all regions in each cell; $\beta$ is the $\mathbb{R}^{N_{\text{topics}} \times N_{\text{peaks}}}$ matrix linking accessibility to the influence of topics; and $\hat{n}_i^{\text{ATAC}}$ is the observed number of accessible peaks in cell $i$. Thus, accessibility in a cell is generated by $\hat{n}_i^{\text{ATAC}}$ independent samples from the categorical distribution over regions. This is the same assumption underlying the generative process of sparse wordcount compositions in a document used in natural language topic modeling. The likelihood function of the multinomial distribution given observed data $X_{i\cdot}^{\text{ATAC}}$ and region composition $\rho_{i\cdot}$ does not depend on the read depth parameter, so we do not learn a variable for effective scATAC-seq read depth.

For both modalities, MIRA takes $\rho_{i\cdot}$ to be the imputed value of the features, representing its estimated rate of occurrence relative to other features (genes or regions) in the same cell.

Notably, $\rho_{i\cdot}$ does not depend on the read depth of a cell, enabling normalized comparisons of feature magnitude across cells with heterogenous capture rates.

MIRA applies batch normalization[6] to the activation of each gene or accessible region given the latent topics of a cell, $Z_{i\cdot}\beta_{\cdot j}$ or $Z_{i\cdot}\beta_{\cdot k}$, respectively. For genes $j$ (or congruently peaks $k$), batch normalization standardizes that activation using summary statistics tracked from previous activation scores across many cells, the batch mean $\mu_j^{\mathrm{bn}}$ and batch standard deviation $\sigma_j^{\mathrm{bn}}$, then projects that quantity to the mean, $b_j$, and standard deviation, $\gamma_j$, of that feature's space:

$$\mathrm{batchnorm}(\ Z_{i\cdot}\beta_{\cdot j}\ ) = \gamma_j \left( \frac{Z_{i\cdot}\beta_{\cdot j} - \mu_j^{\mathrm{bn}}}{\sigma_j^{\mathrm{bn}}} \right) + b_j$$

This normalizes the topic-feature matrix $\beta$ such that the topic-feature activation strengths are dependent on the strength of association and decoupled from the mean variance of the feature counts. This is critical to the analysis of the topics so that the most activated features correspond with the strongest associations rather than the most highly expressed genes or most accessible loci.

*Stochastic variational inference*

Given the observations from cells, $X^{\mathrm{RNA}}$ or $X^{\mathrm{ATAC}}$ as $X$, MIRA finds topics $\beta$; feature means and variances, $\gamma$ and $b$; dispersions $\theta$ (only for expression data); as well as cell-level latent representations $Z$, such that the probability of observing the data is maximized given those parameters $\vartheta$ and conditioned on the latent space:

$$\vartheta_{max} = \mathrm{argmax}_{\vartheta} \log p_{\vartheta}(X)$$

$$p_{\vartheta}(X) = \int p_{\vartheta}(X \mid Z)\, p_{\vartheta}(Z)\, dZ$$

$$\vartheta = (\beta, \gamma, b, \theta)$$

The integral for the marginal likelihood of the model is intractable, so the values of the parameters cannot be solved analytically. Instead of using Monte Carlo sampling-based methods, MIRA employs the variational autoencoder approach[2] which is based on a variational approximation of the distribution $p_{\vartheta}(Z \mid X)$, and the observation that the marginal distribution is related to the posterior predictive distribution of $Z$ by Bayes rule.

The variational distribution $q$, conditioned on the observations $X$, is represented by an encoder neural network with weights $\phi$:

$$q_{\phi}(Z \mid X) \approx\ p_{\vartheta}(Z \mid X)$$

$$Z \sim\ q_{\phi}(Z \mid X) =\ \mathrm{Encoder}_{\phi}(X)$$

to approximate the marginal likelihood of the model. The encoder neural network uses the observations of $X$ to parameterize the distribution from which $Z$ is sampled. MIRA provides the generative model and its parameters $\vartheta$, and the variational sampling method and its parameters $\phi$, to Pyro's stochastic variational inference function[7]. Pyro then jointly estimates the parameter values by maximizing the evidence lower bound (ELBO) objective[3] using stochastic gradient ascent, which maximizes the probability of the observed data given the variational approximation $q_\phi(Z \mid X)$, while minimizing the Kullback-Leibler (KL) divergence between variational distribution $q_\phi(Z \mid X)$ and the prior distribution $p_\vartheta(Z)$. We assume the latent variables are independent, which satisfies Pyro's mean field condition and enables Pyro's use of analytical expressions for KL divergence. During inference, Pyro learns point estimates for all parameters $\vartheta$ and $\phi$.

*Variational reparameterization*

To train the variational parameters of the model using gradient ascent, MIRA reparametrizes the latent variable sampling scheme in terms of normal distributions, enabling Pyro to find unbiased Monte Carlo estimates of the ELBO expectation's gradient[2]. MIRA recasts the Dirichlet prior as implemented by ProdLDA[8]:

$$Z_{i\cdot} \sim \mathrm{softmax}\big(\mathrm{Normal}(\,\mu^Z_{\cdot}, \mathrm{diag}((\sigma^Z_{\cdot})^2)\,)\big) \approx \mathrm{Dirichlet}\left(\alpha_1, \dots, \alpha_{N_{\text{topics}}}\right), \forall\, i \in \{1, \dots, N_{\text{cells}}\}$$

$$\mu_t^Z = \log \alpha_t - \frac{1}{N_{\text{topics}}} \sum_{\tau=1}^{N_{\text{topics}}} \log \alpha_\tau, \forall\, t \in \{1, \dots, N_{\text{topics}}\}$$

$$\left(\sigma_t^Z\right)^2 = \frac{1}{\alpha_t}\left(1 - \frac{2}{N_{\text{topics}}}\right) + \frac{1}{N_{\text{topics}}\alpha_t}, \forall\, t \in \{1, \dots, N_{\text{topics}}\}$$

$$\alpha_t \sim \mathrm{LogNormal}\left(\alpha_\mu, \alpha_\sigma^2\right) \approx \mathrm{Gamma}(2, \frac{2N_{\text{topics}}}{\mathcal{J}}), \forall\, t \in \{1, \dots, N_{\text{topics}}\}$$

$$\alpha_\mu = \log \frac{\mathcal{J}}{\sqrt{\frac{3}{2}}\, N_{\text{topics}}}, \qquad \alpha_\sigma^2 = \log \frac{3}{2}$$

The $\alpha_\mu$ and $\alpha_\sigma$ parameters specify a log-normal distribution with the same mean and variance as the generative gamma distribution, and softmax of samples from the normal distribution parameterized by $\mu_t^Z$ and $\sigma_t^Z$ approximates the Dirichlet prior over topics. Using the output layer of the encoder neural network (see MIRA Topic Model: *Stochastic variational inference* section) conditioned on the observations from the cell, MIRA defines the variational distributions of the latent topics for each cell:

$$v_{i\cdot}^{\text{output}} = \mathrm{Encoder}_\phi(X_{i\cdot})$$

$$\mu_{i\cdot} = (v_{i1}^{\text{output}}, \dots,\ v_{iN_{\text{topics}}}^{\text{output}})$$

$$\sigma_{i\cdot} = \text{softplus} \circ (v^{\text{output}}_{i,(N_{\text{topics}}+1)}, \dots, \ v^{\text{output}}_{i,(2N_{\text{topics}})})$$

For expression model only:

$$\mu_i^{n_i} = v^{\text{output}}_{i,(2N_{\text{topics}}+1)}$$

$$\sigma_i^{n_i} = \text{softplus}(v^{\text{output}}_{i,(2N_{\text{topics}}+2)})$$

For cell $i$, the neural network output $v_{i\cdot}^{\text{output}} \in \mathbb{R}$ gives a $(2N_{\text{topics}} + 2)$-dimensional vector for expression or a $2N_{\text{topics}}$-dimensional vector for accessibility, which provides estimates of the mean $\mu_{i\cdot} \in \mathbb{R}^{N_{\text{topics}}}$ and standard deviation $\sigma_{i\cdot} \in \mathbb{R}^{N_{\text{topics}}}$ parameters for the variational distribution of $Z_{i\cdot}$ for that cell. For expression data, the encoder network also parameterizes the distribution of effective read depth $n_i \sim \text{LogNormal}(\mu_i^{n_i}, (\sigma_i^{n_i})^2)$. When specifying standard deviation parameters for the variational distribution, we found the non-negative softplus transformation[9] to be more numerically stable than the exponential transformation. Above, $\circ$ refers to the composition of the softplus function over the vector output of the encoder network.

*Encoder network architecture*

The encoder neural network takes the observations of a given modality as features and outputs a parameterization for the latent representation for the cell. MIRA adapts the architecture of the encoder neural network to fit the properties of that modality. For gene expression data, MIRA first transforms raw count observations to normalized quantities using deviance residual featurization[10] of highly variable genes, which are then passed through the neural network. The deviance residuals $r_{ij}$ of the raw counts $X_{ij}^{\text{RNA}}$ regress out the effects of count variation and circumvent count distortions induced by traditional log-plus-one featurization of expression count data, providing a better initial representation of cell state for decomposition into topics by the encoder network.

MIRA passes the deviance residuals through two hidden layers of a feed-forward neural network and an output layer. Each layer consists of a fully-connected layer, batch normalization, ReLU activation[11], then dropout[12]. The hidden layers have 128 nodes. Thus, the encoder network conditioned on expression of cell $i$ is given by:

$$f^l(v) = \text{dropout}(\ \text{ReLU}(\ \text{batchnorm}(\ W^l v + b^l)))$$

$$r_{i\cdot} = \text{DevianceResiduals}(X_{i\cdot}^{\text{RNA}})$$
$$v_{i\cdot}^0 = f^0(r_{i\cdot} \oplus \log \hat{n}_i^{\text{RNA}})$$
$$v_{i\cdot}^1 = f^1(v_{i\cdot}^0)$$
$$v_{i\cdot}^{\text{output}} = \text{batchnorm}(W^2 v_{i\cdot}^1 + b^2)$$

where $f^l$ is the function of the $l^{th}$ layer of the encoder network, $W^l$ and $b^l$ are the weights associated with that layer, and $v_{i\cdot}^l \in \mathbb{R}^{N_{\text{nodes}}}$ is the output of the $l^{th}$ layer for the $i^{th}$ cell. The output layer of the encoder network specifies parameters $v_{i\cdot}^{\text{output}}$ for the latent variational distribution (see MIRA topic model, *Variational reparameterization* section) and is not subject to ReLU nonlinearity or dropout. We inject the observed read depth of the cell $\hat{n}_i^{\text{RNA}}$ into the first feed-forward layer of the neural network by concatenating it ($\oplus$) to the deviance residual features.

The encoder network for chromatin accessibility data requires a different model architecture due to the large number of peaks with high degree of sparsity. MIRA uses a Deep Averaging Network (DAN)[13], which averages embedding vectors of all features found in the sample before passing that resultant vector through successive feed-forward layers. Applied to accessibility data, each site is associated with a 128-dimensional vector, and those vectors are averaged for every accessible site in a cell. The averaged vector passes through a hidden layer of the same specification as the expression encoder, then an output layer. The output of the DAN network for cell $i$ is given by:

$$v_{i\cdot}^{0} = \frac{1}{|\Omega_i|}\sum_{k\,\in\,\Omega_i} W_{k\cdot}^{0}$$
$$v_{i\cdot}^{1} = f_1\left(v_{i\cdot}^{0} \oplus \log \hat{n}_i^{\text{ATAC}}\right)$$
$$v_{i\cdot}^{\text{output}} = \text{batchnorm}(W^2 v_{i\cdot}^{1} + b^2)$$

where $W_{k\cdot}^{0} \in \mathbb{R}^{N_{\text{nodes}}}$ denotes the embeddings for each peak, and $v_{i\cdot}^{0}$ is the average of the embedding vectors in $\Omega_i$, which is the set of accessible peaks $X_{ik}^{\text{ATAC}}$ in the cell $i$ regularized by leaving out peaks at a rate given by Bernoulli trials with parameter $d$. Again, we inject the read depth of the cell $\hat{n}_i^{\text{ATAC}}$ into the first feed-forward layer of the neural network.

*Feature selection and training procedure*

To increase training speed, the number of input features used by the encoder can be limited by selecting highly variable genes in expression data and optionally randomly down-sampling peaks in ATAC-seq data for samples with a large number (>200,000) of peaks. On the other hand, topic enrichments may be more relevant when including additional genes and peaks that may not have met arbitrary feature selection cutoffs. For this reason, MIRA topic models may learn patterns in a superset of features while only utilizing a subset as features for the encoder network.

MIRA maximizes the ELBO objective by gradient ascent using the ADAM optimizer[14]. We adapt the learning rate of the optimizer during training using the one-cycle learning rate policy[15]. In two phases, the learning rate starts small and peaks one-third of the way through training, then slowly diminishes over the remainder of training. We set the initial and maximum learning rates using the learning rate range test[15].

To prevent node collapse (when topics settle into insurmountable local minima early in training) we employ KL annealing of the ELBO objective[16] (see MIRA Topic Model: *Stochastic variational inference* section). The KL term exerts a strong regularizing influence through the prior $p_{\vartheta}(Z)$, which can dominate the gradient early in training and reduce expressivity of the model. Initially, the KL term weight is set to zero and increases linearly until plateauing at one, which occurs half-way through training.

*Hyperparameter optimization*

MIRA includes a rigorous hyperparameter tuning scheme to ensure the model captures informative, non-redundant topics in the data. The most influential parameter on downstream analysis is $N_{\text{topics}}$, the number of topics, which is tuned along with ε, the smoothing parameter for the ADAM optimizer steps, the dropout rate of the encoder neural network, batch size, and the number of epochs trained. We evaluate a given specification of the model using the negative ELBO as the loss on a held-out set of cells. Empirically, the model loss appears to be stochastically convex and separable with respect to each of these parameters, meaning they may be jointly tuned using zero-order optimization of the model loss with respect to the hyperparameter values to approach the most optimal model for a given dataset.

For each iteration of hyperparameter optimization, MIRA uses a Tree of Parzen estimator[17] (TPE) implemented by Optuna[18] to suggest a new combination of hyperparameters that may improve on the previous best model. TPE is a Bayesian method for hyperparameter selection that uses pre-defined priors over the parameter space and evidence from previous trials to inform the next suggested hyperparameter combination.

To evaluate a set of hyperparameters recommended by TPE, MIRA performs five-fold cross-validation on a training set of cells and reports the average loss across all folds. To prevent excessive time spent on poorly performing models, each fold's loss is compared to previous trials, and the trial is terminated early if the current iteration's model does not meet the criteria of a successive halving bandit[19] with a reduction factor of three. The scores of early-terminated trials are penalized by the addition of a trial penalty factor, $P$, to the average loss:

$$P = P_0 2^{1-n}$$

where $n \in \{1, \ldots, 4\}$ is the cross-validation fold at the point of early termination. The penalty decreases for each fold tested and encourages TPE to explore the parameter spaces of trials that survived for more folds. Tuning may be run for a set number of iterations, 32 for large datasets with lower variance model performance estimates or 75 for small datasets, or until TPE converges and repeatedly suggests a similar number of topics.

After the tuning phase, the top five models are trained on the entire training set of cells, and performance compared on a held-out test set of cells. The best performing model from this phase is selected as the final model of the data and retrained on all available cells. MIRA repeats these optimization steps for each modality.

*Topic Analysis*

Given a trained topic model, the $\beta$ matrix encodes the linear associations between topics and expression or accessibility features. To get the normalized activation $\psi_{tj} \in \mathbb{R}$ of a gene $j$ (or congruently peak $k$) given topic $t$, we scale the value of the $\beta$ matrix using the learned batch normalization function's feature-specific variance and bias parameters:

$$\psi_{tj} = \operatorname{sign}(\gamma_j) \frac{\beta_{tj} - \mu_j^{\text{bn}}}{\sigma_j^{\text{bn}}}$$

The distribution of activations across all genes and topics is roughly standard normal and is not skewed by the variance and mean levels of the feature. The top $n$ features most strongly associated with a topic are given by the top $n$ activation scores.

To annotate expression topics, MIRA extracts the top $n$ genes and passes the geneset to Enrichr[20] for comparison to precompiled ontologies. To annotate accessibility topics, MIRA extracts the top $c$ percentile of most activated peaks in a topic, then finds transcription factors (TFs) with predicted binding sites (by either motif analysis as described below or occupancy if provided chromatin immunoprecipitation-sequencing (ChIP-seq) data) enriched in the most activated peaks versus the remaining peaks using the Fisher exact test[21], implemented by scipy[22]. The Fisher exact test gives a fast approximation of Monte Carlo-based simulations of the null distribution of intersection between two sets of genomic regions[23].

*Joint representation*

The topic composition of cell $i$ is given by the expected value of the variational approximation of the posterior of $Z_{i\cdot}$, denoted $\hat{Z}_{i\cdot}$:

$$\hat{Z}_{i\cdot} = \mathbb{E}\left[q_\phi(Z_{i\cdot} \mid X_{i\cdot})\right] \approx \mathrm{softmax}(\mu_{i\cdot}),$$

where $q$ is the variational distribution parameterized by the encoder neural network conditioned on the observed features of cell $i$ and mean $\mu_{i\cdot} \in \mathbb{R}^{N_{\text{topics}}}$ is given by the output layer of the network (see MIRA topic model: *Variational reparameterization* section). MIRA projects the $N_{\text{topics}}$-dimensional simplex space topic compositions for each cell to $(N_{\text{topics}} - 1)$-dimensional real space using the isometric log-ratio transformation (ILR)[24]:

$$\mathrm{ILR}(\hat{Z}_{i\cdot}) = \left(\log\frac{\hat{Z}_{i1}}{g(\hat{Z}_{i\cdot})}, \dots, \log\frac{\hat{Z}_{iN_{\text{topics}}}}{g(\hat{Z}_{i\cdot})}\right) \cdot G$$

$$g(\hat{Z}_{i\cdot}) = \exp\left(\frac{1}{N_{\text{topics}}}\sum_{t=1}^{N_{\text{topics}}} \log \hat{Z}_{it}\right)$$

$$G_{t\tau} = \begin{cases} \dfrac{\sqrt{\tau/(\tau+1)}}{\tau} & \text{if } t < \tau + 1 \\ -\sqrt{\tau/(\tau+1)} & \text{if } t = \tau + 1 \\ 0 & \text{if } t > \tau + 1 \end{cases}$$

$$\text{for } t \in \{1, \dots, N_{\text{topics}}\}$$
$$\text{and } \tau \in \{1, \dots, (N_{\text{topics}} - 1)\}$$

where $g(\hat{Z}_{i\cdot})$ is the geometric mean of the composition of $\hat{Z}_{i\cdot}$, and $G \in \mathbb{R}^{N_{\text{topics}} \times (N_{\text{topics}} - 1)}$ is a Gram-Schmidt orthonormalized basis matrix derived from an arbitrary hierarchical relationship between topic compositions[25]. Transformation to $(N_{\text{topics}} - 1)$-dimensional space by the $G$ matrix aligns topic activations along an orthogonal basis. To create a joint representation encoding information from both modalities, MIRA concatenates the isometric log-ratio transformed vectors for expression and accessibility topics into one vector representing the multimodal cell state, $J_{i\cdot} \in \mathbb{R}^{N_{\text{topics}}^{\text{RNA}} + N_{\text{topics}}^{\text{ATAC}} - 2}$ :

$$J_{i\cdot} = \text{ILR}(\hat{Z}_i^{\text{RNA}}) \oplus \text{ILR}(\hat{Z}_i^{\text{ATAC}}), \text{for } i \in \{1, \dots, N_{\text{cells}}\}$$

Using the Manhattan distance between cells in the joint space, MIRA constructs a k-nearest neighbors (KNN) graph where edges represent cells with similar transcriptional and accessibility states. Assuming transitions between topics capture major biological state changes, those changes would be aligned along the axes in orthonormal ILR-transformed space. Therefore, the Manhattan distance represents the distance between cells as the transitions required to traverse the axes along topics to arrive at the other cells' biological state. In addition, the Manhattan distance has also been shown to preserve nearest-neighbor relationships in high-dimensional space better than Euclidean distance[26].

MIRA implements an extension of the ILR transformation, where the log ratio of the topic composition and its geometric mean may be substituted for any box-cox transformation, and the geometric mean is substituted for the generalized mean[27]. This introduces a parameter to the construction of the joint representation, the box-cox power transformation parameter $a$. As $a$ approaches zero, this extension produces the classic ILR transformation.

The joint KNN graph may be used for clustering by the Leiden algorithm[28] and low-dimensional visualization using UMAP[29]. To generate UMAP visualizations, we use the default parameters given by the umap-learn Python package.

*Motif score*

Using the JASPAR CORE collection[30], we call motifs hits within scATAC-seq peaks with the MOODS3 algorithm[31]. The adjusted p-value threshold is set to p<1e-5. Then, we calculate motif scores $Q \in \mathbb{R}^{N_{\text{cells}} \times N_{\text{factors}}}$ for each cell and each factor using the query likelihood model[32]. The score for TF $h$ in cell $i$ is given by the log-probability of sampling the set of regions predicted to be bound by $h$, $\mathfrak{C}_h$ (the cistrome of $h$), from the distribution of regions $K$ given by the ATAC topic model:

$$Q_{ih} = \sum_{k \in (\mathfrak{C}_h \cap K)} \log \hat{\rho}_{ik}^{\text{ATAC}} \quad \text{for } i \in \{1, \dots, N_{\text{cells}}\} \text{ and } h \in \{1, \dots, N_{\text{factors}}\}$$

where $\hat{\rho}_{i\cdot}^{\text{ATAC}}$ is the composition of peaks in a cell given by the mean variational estimate of the latent topics $\hat{Z}_i$. The $Q$ matrix is first normalized such that the factor scores in a cell have a Euclidean norm of 1, then each factor's scores are standardized to the standard normal distribution across all cells for comparability.

**Pseudotime trajectory inference**

*Transport map construction*

A transport map, or Markov chain model $\pi \in \mathrm{I}^{N_{\text{cells}} \times N_{\text{cells}}}$ where $\mathrm{I}$ is the unit interval [0,1], describes the transition probabilities between cells progressing through a differentiation system:

$$\sum_{\zeta=1}^{N_{\text{cells}}} \pi_{i\zeta} = 1\,, \forall\ i \in \{1, \dots, N_{\text{cells}}\}$$

where $\pi_{i\zeta}$ is the probability of transitioning from cell $i$ to cell $\zeta$ after an arbitrary discrete time step. MIRA uses the Palantir algorithm[33] to transform the undirected joint KNN graph describing cells in similar states into a directed transport map $\pi$ representing the stochastic differentiation process based on multimodal transition probabilities. First, Palantir assigns each cell a pseudotime describing its progress through the differentiation process. Pseudotime $s$ is taken to be the shortest path distance of traversing the joint KNN graph from the origin cell $O$ to each cell $i$. Then, Palantir transforms the undirected joint KNN graph into a directed transport map by pruning edges in the joint KNN graph that travel "backwards" relative to the pseudotemporal flow of cells progressing from the user-chosen origin cell $O$.

From the transport map, MIRA identifies terminal cells where the forward progress of the differentiation reaches a stationary state at the end of each lineage. MIRA finds the left eigenvectors of the transport map whose eigenvalues are approximately one. The cells with the maximum value for each associated eigenvector mark the terminal states[34].

Lastly, MIRA again uses the Palantir algorithm to assign to each cell a probability of reaching each lineage's terminal state following a random walk through the transport map. We denote the probability of reaching the $z^{th}$ terminal state from cell $i$ following a random walk through the joint space derived transport map as $p(J_z \mid J_i)$.

*Lineage tree inference*

Here we describe a novel extension of the Palantir algorithm which uses the cell terminal state probabilities to construct a bifurcating tree structure representation of the data. MIRA determines lineages and branch points using the terminal fate probabilities found by Palantir. First, a lineage $\ell_{Oz}$ is defined as the set of all cells for which the probability of reaching that lineage's terminal state $z$ is greater than or equal to the probability of reaching that terminus from the origin state $O$:

$$\ell_{Oz} = \{\, i \subseteq \{1, \dots, N_{\text{cells}}\} \mid p(J_z \mid J_i) \ \geq p(J_z \mid J_O)\}$$

The branch time $s^*$ between two lineages with terminal states $a$ and $b$ is defined by:

$$s^*(O, a, b) \ = \min\{\, s(i) \mid \ \operatorname{abs}(F_i^{ab}) > \varepsilon, i \in \ \ell_{Oa} \cup \ell_{Ob}\}$$

$$F_i^{ab} = \log \frac{p(J_a \mid J_i) / p(J_b \mid J_i)}{p(J_a \mid J_O) / p(J_b \mid J_O)}, \text{for } i \ \in \ \ell_{Oa} \cup \ell_{Ob}$$

First, all cells in lineages $a$ and $b$ are merged into a combined set of cells, $\ell_{Oa} \cup \ell_{Ob}$, then MIRA calculates $F_i^{ab}$, the log fold change of the ratios between the probability of reaching lineage terminus $a$ versus lineage terminus $b$ at cell $i$ relative to the probability at the start cell $O$. Intuitively, before the branch between two lineages, the ratios of the probabilities of differentiating down two different trajectories is constant, and after the branch point, these probabilities diverge from the initial balance as cells become more likely to reach one terminal state rather than the other. The branch time between two lineages is taken to be the pseudotime $s$ of the first cell where $F_i^{ab}$ exceeds some threshold $\varepsilon$.

To construct a bifurcating lineage tree using these definitions, MIRA starts with all terminal states as disconnected leaves. MIRA first finds the branch times between all lineages, and the lineages which branch latest in the differentiation are merged to create a new super-lineage, where each cell's probability of differentiating into the super-lineage is $p(J_a | J_i) + p(J_b \mid J_i)$. A node is added upstream connecting these lineages' terminal states with a branch point, and all cells in the lineages with a pseudotime greater than the branch time are assigned to the appropriate child of the branch node depending on which lineage they have more affinity to, determined by $\mathrm{sign}(F_i^{ab})$. Then, MIRA recomputes branch times between the lineages to account for the super-lineage and again merges the last-branching trajectories. This process is repeated until all lineages have been connected to the root node and all cells have been assigned to a node.

### MIRA regulatory potential (RP) model

*Model architecture*

The MIRA RP model relates changes in local accessible chromatin to gene expression by learning upstream and downstream distances of perceived regulatory influence that maximize the probability of observing the expression data given the accessibility state in the same single cells. MIRA models the generative process of sampling expression counts for gene $j \in \{1, \dots, N_{\text{genes}}\}$ in cell $i \in \{1, \dots, N_{\text{cells}}\}$ given the accessibility state $A_{i\cdot}$ of the cell as:

$$X_{ij}^{\text{RNA}} \sim \text{NegativeBinomial}\left( n_i \rho_{ij}, \theta_j \right)$$

$$\rho_{ij} = \frac{e^{\lambda_{ij}}}{\sum_{g=1}^{N_{\text{genes}}} \exp\left( \text{batchnorm}_{\text{g}}\left( \hat{z}_{i\cdot}^{\text{RNA}} \beta_{\cdot g} \right)\right)}$$

$$\lambda_{ij} = \gamma_j \left( \frac{c_{ij} - \mu_j^{\text{bn}}}{\sigma_j^{\text{bn}}} \right) + b_j$$

$$c_{ij} = R\left(\mathfrak{D}_{j\cdot}, A_{i\cdot}, a_{j\cdot}, \delta_{j\cdot}, \Delta_{j\cdot}\right) = \sum_{\eta \in \{\text{U,D,P}\}} a_{j\eta} \sum_{\varsigma \in \mathfrak{D}_{j\eta}} A_{i\varsigma} 2^{-\delta_{j\varsigma}/\Delta_{j\eta}}$$

$$A_{i\varsigma} = \hat{\rho}_{i\varsigma}^{\text{ATAC}}$$

For each cell $i$ and gene $j$, the $R$ function takes as arguments: the genomic interval sets $\mathfrak{D}_{j\eta}$ for $\eta \in \{\mathrm{U}, \mathrm{D}, \mathrm{P}\}$ which filters peaks based on strand-oriented positional relationships upstream $(\mathrm{U})$, downstream $(\mathrm{D})$, or proximal $(\mathrm{P})$ to the gene transcription start site (TSS); the accessibility state $A_{i\cdot} \in \mathbb{R}_{\geq 0}^{|\mathfrak{D}_{j\mathrm{U}}|+|\mathfrak{D}_{j\mathrm{D}}|+|\mathfrak{D}_{j\mathrm{P}}|}$ of each locus in a cell; non-negative $a_{\mathrm{U}}, a_{\mathrm{D}}$, and $a_{\mathrm{P}}$ parameters that scale the relative effects of upstream $(\mathrm{U})$, downstream $(\mathrm{D})$, or proximal $(\mathrm{P})$ accessibility ($a_{\mathrm{U}}, a_{\mathrm{D}}, a_{\mathrm{P}} \sim \mathrm{HalfNormal}(0, 1)$), respectively; the distances $\delta_{j\cdot} \in \mathbb{R}_{\geq 0}^{|\mathfrak{D}_{j\mathrm{U}}|+|\mathfrak{D}_{j\mathrm{D}}|+|\mathfrak{D}_{j\mathrm{P}}|}$ from the TSS of gene $j$ to the loci in the specified genomic interval set; and the decay rate parameters $\Delta_{i\mathrm{D}}$ and $\Delta_{i\mathrm{U}}$.

The accessibility of each region in $\mathfrak{D}_{j\eta}$ is weighted by its distance from the TSS in terms of the learned decay rate parameter $\Delta_{i\eta}$, and the effects of all loci are summed together to summarize the *cis*-regulatory effect on gene expression. The accessibility state $A_{i\cdot}$ of loci in cell $i$ is taken to be the predicted compositional distribution $\hat{\rho}_{i\cdot}^{\mathrm{ATAC}}$ given by the chromatin accessibility topic model, to reduce noise and normalize for differences in read depth of ATAC observations between cells. The upstream and downstream region sets encompass regions between 1.5 and 600 kilobases from the TSS; the proximal region is within 1.5 kilobases from the TSS. Regions within 1.5 kilobases of other genes are masked.

The $\Delta_{j\mathrm{D}}$ and $\Delta_{j\mathrm{U}}$ parameters affect the respective downstream $(D)$ and upstream $(U)$ decay rates of local chromatin accessibility's influence on gene expression. The value of the parameter is the estimated distance, in kilobases, over which the influence of accessible sites on gene expression is halved, $\Delta_{\mathrm{U}}, \Delta_{\mathrm{D}} \sim \mathrm{LogNormal}(\log(15),\ 1.44)$. The prior distribution reflects *a priori* information about the likely ranges of regulatory influence[35,36], placing the mean decay distance at 15 kilobases and penalizing extreme ranges which suggest spurious long-range correlations. The variance given by $1.44$ places the 90% and 99% quantiles of the prior over regulatory distances at 69 and 242 kilobases, respectively. Influence of accessibility in the promoter region is not decayed, thus $\Delta_{j\mathrm{P}}$ is set to ∞.

The model relates the *cis*-regulatory relationship $c_{ij}$ to the observed expression data $X_{ij}^{\mathrm{RNA}}$ following the same generative statistical method as the expression topic model. Parameters $\gamma_j$ and $b_j$ form a Bayesian batch normalization function, which disconnects the magnitude of change in accessibility from that of gene expression and reduces the variance of gradient updates: $\gamma_j \sim \mathrm{LogNormal}(0,1)$, $b_j \sim \mathrm{Normal}(\,0, 25\,)$. $\theta_j$ regulates overdispersion of the negative binomial count observations, $\theta_j \sim \mathrm{Gamma}\left(2, \frac{1}{2}\right)$.

The compositional rate of expression $\rho_{ij}$ is estimated by approximating the softmax function of the RNA topic model using the RP model activation $\lambda_{ij}$ for the numerator and topic model activations across all genes $\hat{Z}_{i\cdot}^{\mathrm{RNA}}\beta$ for the denominator. This ensures that the RP model is learning compositional relationships of gene expression consistent with those learned by the RNA topic model and that we may use the same estimated read depth random variable $n_i$ estimated by the RNA topic model.

Notably, this model adjusts for technical variation and noise between both assays to learn regulatory distances describing the *cis*-regulatory relationship between local chromatin and expression.

*Parameter estimation*

The objective is to find parameters $\vartheta$ that maximize the probability of the observed expression $X^{\mathrm{RNA}}$ given the accessibility state $A$:

$$\vartheta_{\max} = \mathrm{argmax}_{\vartheta} \log p_{\vartheta}\left(X^{\mathrm{RNA}} \,\middle|\, A\right)$$

$$\vartheta = \{\, a_{j\cdot},\ \Delta_{j\cdot},\ \gamma_j, b_j,\ \theta_j\}$$

MIRA employs variational inference[3] to estimate the posterior predictive distribution of the parameters given the data, using the variational distribution $q$ and maximizing the ELBO objective. Point estimates for each parameter in the variational distribution are estimated using delta distributions. MIRA takes gradient steps to maximize the ELBO using the 2nd order Limited-memory Broyden–Fletcher–Goldfarb–Shanno algorithm (L-BFGS)[37]. Because the batch normalization parameters $\mu_j^{\mathrm{bn}}$ and $\sigma_j^{\mathrm{bn}}$ are updated after each batch but are not tuned by the optimizer, the gradient history may cause updates to become unstable. To prevent update instability, we implemented Frozen-batch L-BFGS[38], a variant of L-BFGS that improves the algorithm's performance in stochastic settings. MIRA trains until the loss does not decrease by more than a given threshold for three iterations.

*NITE model architecture*

The RP model discussed above is defined as the local chromatin accessibility-influenced transcriptional expression (LITE) model. The LITE model learns a *cis*-regulatory relationship relating expression to local chromatin accessibility. The non-local chromatin accessibility-influenced transcriptional expression (NITE) model augments the LITE model to additionally include knowledge of cell-wide chromatin state through the incorporation of the MIRA latent accessibility topics as features. The specification of the NITE model mirrors the LITE model (see Regulatory Potential Modeling: *Model architecture* section) except for the inclusion of coefficients describing the relationship between cell-wide chromatin topics and expression:

$$c_{ij} = R(\mathfrak{D}_{j\cdot}, A_{i\cdot}, a_{j\cdot}, \delta_{j\cdot}, \Delta_{j\cdot}) + \sum_{t=1}^{N_{\mathrm{topics}}} a_t^{\mathrm{topics}} \hat{Z}_{it}^{\mathrm{ATAC}}$$

$$a_t^{\mathrm{topics}} \sim \mathrm{Normal}(0, 1), \quad \text{for} \quad t \in \{1, \ldots, N_{\mathrm{topics}}\}$$

$$\theta_j^{\mathrm{NITE}} \leftarrow \theta_j^{\mathrm{LITE}}$$

The dispersion parameter $\theta^{\mathrm{NITE}}$ is fixed as the value learned by the LITE model for the same gene so performance differences between the LITE and NITE models are not driven by the effect of dispersion on the distribution of expression. For a given gene, MIRA first trains a LITE model, then seeds the variational distribution of the NITE model with the point estimates from the LITE model. NITE model training proceeds in the same manner as LITE model training and learns new values for each parameter.

### LITE vs. NITE regulation test

To test the ability for local chromatin to predict expression of a gene, we perform a likelihood ratio test[39] between the LITE and NITE models, where the null hypothesis is that the LITE model, based only on local chromatin features, is sufficient to predict expression:

$$\Lambda_{\mathrm{j}} = -2 \log \frac{\mathcal{L}^{\mathrm{LITE}}\left(\rho_{\cdot j}^{\mathrm{LITE}} \middle| X_{\cdot j}^{\mathrm{RNA}}\right)}{\mathcal{L}^{\mathrm{NITE}}\left(\rho_{\cdot j}^{\mathrm{NITE}} \middle| X_{\cdot j}^{\mathrm{RNA}}\right)}, \text{for } j \in \{1, \ldots, N_{\mathrm{genes}}\}$$

Here, $\mathcal{L}^{\mathcal{M}}\left(\rho_{\cdot j}^{\mathcal{M}} \middle| X_{\cdot j}^{\mathrm{RNA}}\right)$ is the likelihood of the expression predictions of model $\mathcal{M}$, the LITE or NITE model for that gene, given the observations of the expression of gene $j$ across all cells, where $X_{\cdot j}^{\mathrm{RNA}} \in \mathbb{Z}_{\geq 0}^{N_{\mathrm{cells}}}$. The LITE and NITE models parameterize a negative binomial distribution of expression given the accessibility state $A_{i\cdot}$ of the cell. Thus, for model $\mathcal{M}$:

$$\mathcal{L}^{\mathcal{M}}\left(\rho_{\cdot j}^{\mathcal{M}} \middle| X_{\cdot j}^{\mathrm{RNA}}\right) = \prod_{i=1}^{N_{\mathrm{cells}}} p\left(X_{ij}^{\mathrm{RNA}} = \mathrm{NegativeBinomial}\left(n_i \rho_{ij}^{\mathcal{M}}, \theta_j^{\mathrm{LITE}}\right)\right)$$

If the expression predictions given the NITE model parameters are more likely given the observed data than the LITE model predictions, this increases the test statistic. The test statistic $\Lambda_{\mathrm{j}}$ is not directly comparable between genes due to differences induced by count variability, so we normalize all genes' test statistics to remove this effect:

$$\text{NITE score}_j = \frac{\Lambda_j}{1 + \frac{\sum_{i=1}^{N_{\mathrm{cells}}} \mathbb{I}(X_{ij} > 0)}{\underset{g \in \{1, \ldots, N_{\mathrm{genes}}\}}{\mathrm{median}} \left(\sum_{i=1}^{N_{\mathrm{cells}}} \mathbb{I}(X_{ig} > 0)\right)}}, \text{for } j \in \{1, \ldots, N_{\mathrm{genes}}\}$$

$$\text{where } \mathbb{I}(\mathrm{True}) = 1 \text{ and } \mathbb{I}(\mathrm{False}) = 0.$$

Due to the properties of expression counts and the negative binomial distribution, both the LITE and NITE models predict zero counts for a gene with high probability. Thus, cells with no reads observed for a given gene are not as informative to the test, and genes which have a smaller fraction of zero counts have larger test statistics. Above, we scale the test statistic for each gene based on the number of nonzero counts relative to the median nonzero counts across all genes tested. When the number of nonzero counts for a gene is greater than the median the penalty to the test statistic increases. This procedure yields a comparable NITE score for each gene.

**Cell NITE score**

The cell NITE score is calculated similarly to gene NITE score, except the test is performed on rows of the expression matrix $X^{\mathrm{RNA}}$ instead of columns:

$$\Lambda_i = -2\log\frac{\mathcal{L}^{\mathrm{LITE}}\left(\rho_{i\cdot}^{\mathrm{LITE}} \middle| X_{i\cdot}^{\mathrm{RNA}}\right)}{\mathcal{L}^{\mathrm{NITE}}\left(\rho_{i\cdot}^{\mathrm{NITE}} \middle| X_{i\cdot}^{\mathrm{RNA}}\right)},\ \text{for } i = 1, \ldots, N_{\mathrm{cells}}$$

$$\text{NITE score}_i = \frac{\Lambda_i}{1 + \dfrac{\sum_{j=1}^{N_{\mathrm{genes}}} \mathbb{I}(X_{ij} > 0)}{\underset{k\in\{1,\ldots,N_{\mathrm{cells}}\}}{\mathrm{median}} \left(\sum_{j=1}^{N_{\mathrm{genes}}} \mathbb{I}(X_{kj} > 0)\right)}},\ \text{for } i = 1, \ldots, N_{\mathrm{cells}}$$

**Chromatin differential**

The chromatin differential $\chi$ in cell $i \in \{1, \ldots, N_{\mathrm{cells}}\}$ for gene $j \in \{1, \ldots, N_{\mathrm{genes}}\}$ is given by:

$$\chi_{ij} = \log\frac{\rho_{ij}^{\mathrm{LITE}}}{\rho_{ij}^{\mathrm{NITE}}}$$

which is the log-ratio of the compositional prediction of expression given by the LITE and NITE models.

**Probabilistic *in silico*-deletion**

*Gene-TF associations*

MIRA makes use of the LITE model and probabilistic *in silico*-deletion (pISD) to predict the TFs that regulate a gene or set of genes[40]. This method assesses the strength of association between a gene and the observed or predicted TF binding sites by probing how the LITE model performance is affected by masking out scATAC-seq reads from that TF's binding sites. TFs that severely degrade the LITE model's predictive strength are more likely to be regulators of the gene than TFs for which binding site masking has no effect on the model's predictive strength. This method reveals TFs that bind regions where accessibility correlates with a given gene's expression.

As such, the pISD test compares the ability of the LITE model to predict expression $X_{\cdot j}^{\mathrm{RNA}}$ given the local chromatin accessibility around a gene, relative to its predictive ability after masking the set of accessible sites $\mathfrak{C}_h$ predicted to be bound by a given TF $h$. If the TF $h$ is predicted to bind regulatory regions that degrade the ability of the model to predict expression, this increases the value of the association score $\mathcal{A}_{jh}$.

MIRA's association score between gene $j$ and TF $h$ is given by the likelihood ratio test:

$$\mathcal{A}_{jh} = -2\log\frac{\mathcal{L}^{\hbar}\left(\rho_{\cdot j}^{\hbar} \middle| X_{\cdot j}^{\mathrm{RNA}}\right)}{\mathcal{L}\left(\rho_{\cdot j} \middle| X_{\cdot j}^{\mathrm{RNA}}\right)}$$

The denominator describes the likelihood $\mathcal{L}$ of the expression predictions $\rho_{\cdot j}$ of the LITE model given $X_{\cdot j}^{\mathrm{RNA}}$ using all nearby accessible regions. The numerator describes the likelihood $\mathcal{L}^{\hbar}$ of the LITE model prediction of the expression of gene $j$ when the regions $\mathfrak{C}_h$ predicted to bind $\mathrm{TF}_h$ are masked (modeling the $\mathrm{TF}_h$'s deletion $\hbar$), and is given by:

$$\mathcal{L}^{\hbar}\left(\rho_{\cdot j}^{\hbar} \middle| X_{\cdot j}^{\mathrm{RNA}}\right) = \prod_{i=1}^{N_{\mathrm{cells}}} p\left(X_{ij}^{\mathrm{RNA}} = \mathrm{NegativeBinomial}\left( n_i \rho_{ij}^{\hbar}, \theta_j^{\mathrm{LITE}} \right)\right)$$

$$\rho_{ij}^{\hbar} = \frac{e^{\lambda_{ij}^{\hbar}}}{\sum_{g=1}^{N_{\mathrm{genes}}} \exp\left( \mathrm{batchnorm}_g\left( \hat{Z}_{i\cdot}^{\mathrm{RNA}} \beta_{\cdot g} \right)\right)}, \mathrm{for}\ i \in \{1, \dots, N_{\mathrm{cells}}\}$$

$$\lambda_{ij}^{\hbar} = \gamma_j \left( \frac{c_{ij}^{\hbar} - \mu_j^{\mathrm{bn}}}{\sigma_j^{\mathrm{bn}}} \right) + b_j, \mathrm{for}\ i \in \{1, \dots, N_{\mathrm{cells}}\}$$

$$c_{\cdot j}^{\hbar} = \mathrm{QuantileNorm}\left(R\left(\mathfrak{D}_{j\cdot} \backslash \mathfrak{C}_h, A_{\cdot\cdot}, a_{j\cdot}, \delta_{j\cdot}, \Delta_{j\cdot}\right), c_{\cdot j}\right)$$

$$R\left(\mathfrak{D}_{j\cdot} \backslash \mathfrak{C}_h, A_{\cdot\cdot}, a_{j\cdot}, \delta_{j\cdot}, \Delta_{j\cdot}\right) = \sum_{\eta \in \{\mathrm{U,P,D}\}} a_{j\eta} \sum_{\varsigma \in \mathfrak{D}_{j\eta} \backslash \mathfrak{C}_h} A_{i\varsigma} 2^{-\delta_{j\varsigma}/\Delta_{j\eta}}$$
$$\mathrm{for}\ i \in \{1, \dots, N_{\mathrm{cells}}\}$$

$$A_{i\varsigma} = \hat{\rho}_{i\varsigma}^{\mathrm{ATAC}}$$

The same values for the learned parameters $a_{j\cdot}$, $\gamma_j$, $b_j$, $\Delta_{j\cdot}, \mu_j^{\mathrm{bn}}$, and $\sigma_j^{\mathrm{bn}}$ are applied to the masked and the unmasked models and are determined from the maximum *a posteriori* estimate of the unmasked model. Masking regions around the gene reduces the amount of observed accessible chromatin in the LITE model and induces a downward shift in the value of the RP, which may confound detection of binding in binding sites that support driving of expression. To compensate for the shift, we perform quantile normalization[41] of the *cis*-regulatory prediction of the masked model to the distribution of the prediction from the unmasked model, mapping the distribution of $c_{\cdot j}^{\hbar}$ to $c_{\cdot j}$. This ensures that when the predictions are passed through the generative statistical model of expression, the difference in probability of observed expression is not influenced by the mean shift. Instead, the difference in probability is driven by differences in the ordering of predictions. For example, if a given gene's expression is solely defined by accessibility of a single upstream enhancer, masking that region will render the accessibility states indistinguishable whether the gene is expressed or not expressed. This drives an increase in the test statistic.

Because the predicted binding sites of TFs, whether given by motifs or by ChIP-seq occupancy, can be noisy and inaccurate, driver TF analysis gains statistical power from testing many genes against many TFs. However, testing every gene in every cell against every TF quickly becomes computationally intensive. Therefore, MIRA down-samples the cells used for each gene in the pISD test based on stratified sampling of its expression quanta. First, MIRA takes the log of expression in each cell and adds a pseudocount equal to mean log expression of the gene across all cells. Then the cells are sorted based on their expression level and divided into quanta such that the first group are the cells with the highest expression that comprise expression proportional to $\frac{1}{N_{\text{bins}}}$ worth of the total expression, the second group are the cells with the next-highest expression comprising $\frac{1}{N_{\text{bins}}}$ of the total expression, *et cetera*. An equal number of cells are taken from each bin so that a diverse array of expression states are sampled, but more informative highly-expressed states are prioritized. By default, 1500 cells are selected.

*Gene set driver TF test*

Since associations between individual genes and TFs are noisy due to the inability to ascertain a TF's true binding sites and regulatory influence in a particular cell, we instead test for shared influence of a TF across multiple genes with similar dynamics or properties. We predict TFs driving expression of a query set of genes using a one-sided Mann-Whitney U test[42] over the association scores $\mathcal{A}$. For each TF, MIRA compares the query gene set's association scores with that TF versus a background gene set's association scores to find TFs with significantly higher association with the query set. By default, the background gene set is taken to be all genes for which association scores were estimated which are not in the query set.

**Stream graphs**

MIRA renders stream graphs using Matplotlib[43]. In stream mode, the value of a feature at each pseudotime point is calculated by Savitzky-Golay[44] filter over a user-defined window size. Features are ordered by the pseudotime of their maximum value, which roughly layers features in the order in which they appear on the stream. In heatmap mode, each box represents the average value of cells within that window.

**Benchmarking**

We performed benchmarking to compare MIRA to standard methodologies in tasks of manifold construction and lineage inference using established metrics. First, we wrote a Python program, Frankencell (https://github.com/AllenWLynch/frankencell-dynverse), to generate a series of synthetic differentiation trajectories by mixing reads from individual cells sampled from distinct, well-defined cell populations from real multiomic single cell RNA-seq and ATAC-seq data. This approach provides us with systems that maintain the complexity of real single cell data jointly represented by expression and accessibility modes, but for which we know the ground truth state of cells and can objectively and quantitatively compare analysis approaches. We utilized a 10x Genomics Multiome dataset of peripheral blood mononuclear cells (PBMCs)[45] where we identified four clearly distinguishable cell populations corresponding to B cells, T cells, natural killer (NK) cells, and monocytes.

For the expression data, we used the Cell Ranger[46] gene expression count matrix provided by 10x Genomics. For the accessibility data, we used MACS2[47] to identify peaks from pseudobulk data representing each of the four distinct cell populations described above. We called peaks for each population using fragments with a length of less than 150 base pairs, which are more likely to be associated with nucleosome depleted cis-regulatory regions[48,49].

Peaks were then merged using the iterative overlap peak merging procedure[50] and peaks common between populations were filtered, resulting in a test set of 63,074 genomic regions. All fragments were intersected with this peak set to aggregate a peak-count matrix for use in downstream analyses.

We constructed a tree-structured scaffold of artificial cell states where the root origin is primarily composed of B cell reads and the three terminal leaf nodes are primarily composed of T cell, monocyte, or NK cell reads, respectively. “Monocytes”, or synthetic cells primarily containing reads from monocytes, branch from the root population first, then “T cells” and “NK cells” bifurcate later in the synthetic differentiation. The root is labeled blue in Fig. 1e and Extended Data Fig. 3-5. We specified the lineage tree as a directed acyclic graph, where trajectories followed edges marking transitions between cell state nodes, and nodes represented ground truth branch points in the synthetic scaffold. For each node in the lineage tree, we defined the proportions of reads to be mixed from each of the reference single cell populations, which we refer to as mixing proportions. Transitions between nodes were modeled as sigmoidal transformations from the mixing proportions of one node to the next with respect to time. We use sigmoidal rather than linear transformations to simulate the nature of cell state transitions in biological systems[51].

To generate a synthetic cell, we first sampled a path through the lineage tree according to a Markov chain model where, starting from the root node and stepping through the nodes in the tree until reaching a terminus, the probability of transitioning to any child of the current node was uniform. We then sampled a progress value between 0 and 1 representing a position along that path, according to a beta (0.5,1) distribution. A progress value of 0 defines a cell at the start of the path or root of the tree, while 1 defines a cell at the terminus of the sampled path. Using the sampled progress to place the cell on an edge of the lineage tree, we calculated the cell’s read mixing proportions based on the sigmoidal interpolation of node-defined mixing proportions. Given the mixing proportions of the nodes defining the start and end of a directed edge of the lineage graph, $\pi_{\text{start}}$ and $\pi_{\text{end}}$, respectively, the mixing proportion of a cell progressed $p$ fraction down that edge was calculated as:

$$\pi_{\text{cell}} = \pi_{\text{start}}(1-\delta) + \pi_{\text{end}}\delta$$

$$\delta = \sigma\big(s(p-0.5)\big)$$

where $\sigma$ denotes the sigmoid function and $s$ governs aggression of the sigmoid transformation. Small values of $s$ create a more gradual transformation between nodes, while larger values specify a more rapid transition (or a depletion of cells with an intermediate phenotype). For all scaffolds, we set $s$ to 6.

We termed a collection of synthetic cells and their associated mixing proportions a “scaffold”, which represents a construction plan for sampling reads from the real single cell dataset. Using this generation process, we specified four different scaffolds of 1000 cells each. We varied the difficulty of each scaffold by adjusting the cell type proportions at the leaf and internal nodes of the lineage tree such that more difficult scaffolds have more similar terminal nodes. We defined the following node structure for the scaffolds. The root node, 0, has one child, node 1. Node 1 has nodes 2 and 3 as children, and node 2 has nodes 4 and 5 as children.

For each scaffold, we adjusted the mixing proportions of each node, $\pi_{\text{node}}$, according to the scaffold-specific difficulty parameter $k$:

$$\pi_0 = [1, 0, 0, 0]$$
$$\pi_1 = \left[0.5, 0.25 + k, 0.125 - \frac{k}{2}, 0.125 - \frac{k}{2}\right]$$
$$\pi_2 = [0.1 + \ k, 0.1 + k, 0.4 - k, 0.4 - k]$$
$$\pi_3 = [k, 1 - k, 0, 0]$$
$$\pi_4 = [k, 0, 1 - 3k \ , 2k],$$
$$\pi_5 = [k, 0, 2k \ , 1 - 3k],$$

$$\text{for k} = 0,\ 0.05,\ 0.075,\ 0.1$$

$$\pi_{\text{node}} = [P_{\text{B cell}}, P_{\text{monocyte}}, P_{\text{NK cell}}, P_{\text{T cell}}], \qquad \sum \pi_{\text{node}} = 1$$

where $P$ is the proportion of reads contributed from the denoted cell cluster to synthetic cells at that node. Scaffolds with larger values of $k$ mix greater proportions of interfering populations' reads into the nodes, making manifold learning and lineage inference more difficult.

To generate a multimodal dataset from a scaffold, read depths of the synthetic cells were sampled from a lognormal distribution. To sample reads to represent each synthetic cell, we randomly selected one real single cell from each population in the reference dataset and hypergeometrically sampled reads from those cells to fulfill their respective contributions according to the mixing proportions.

For each of the four scaffolds, we tested performance for three different mean read depths by sampling reads to mean depths of 1000, 2000, or 4000 RNA reads per cell and 3500, 7000, and 14000 ATAC reads per cell. Lower read depth further increases the difficulty of solving the topology. For each scaffold and read depth condition, we generated 5 replicates to find the variance of model performance. Each dataset and associated ground truth scaffold was stored in dynverse's[52] common trajectory model format to facilitate analysis of inferred lineage inference using established metrics. For all tests, we provided the ground-truth start and terminal cells to the inference methods. On Zenodo (https://doi.org/10.5281/zenodo.6390740), we provide the generated data matrices for all tests, along with the reference single cell datasets and seed configuration files needed to regenerate the tests using the Frankencell software.

We evaluated model performance on three key metrics defined in the dynverse package [52]: edge accuracy, branch F1 score, and pseudotime correlation. Edge accuracy (dynverse's edge flip score) measures the minimal number of edge additions or subtractions needed to convert the test model's inferred trajectory graph into the ground truth graph, divided by the total number of edges in both graphs. Pseudotime correlation measures the correlation of inferred pseudotemporal distances between cells versus ground truth distances. Branch F1 score quantifies the closeness of the predicted cell lineage assignment compared to the ground truth. All metrics were calculated using the dynverse package. The overall score presented in Fig. 1c-d represents the geometric mean of each of the three metrics, standardized across all tests and taken to (0,1) by the sigmoid function (consistent with dynverse[52] methodology).

We compared MIRA using expression data alone, MIRA using accessibility data alone, and MIRA jointly using both expression and accessibility data with standard alternative methodologies. We utilized dynverse's Slingshot wrapper[53] shown in Saelens et al. to perform favorably against all contemporaneous competitors, as the benchmark trajectory inference method. We compared MIRA against Slingshot paired with standard dimensionality reduction techniques as discussed below.

For expression data, we followed conventional highly variable gene selection, then computed reduced dimensions via PCA as implemented by irlba[54] (used by Seurat[55]). dynverse's Slingshot wrapper selects the number of PCA dimensions to use in lineage inference using the "elbow method"[56] on the sorted magnitude of the eigenvalues. To ensure that this selection method did not underestimate the dimensions needed to construct an informative manifold, we tested enforcing the minimum number of dimensions at 3, 4, and 5. We found larger numbers of dimensions did not improve results.

For accessibility data, we compared MIRA against Slingshot paired with two forms of latent semantic indexing (LSI): 1) Seurat standard term frequency-inverse document frequency (TF-IDF) featurization of scATAC-seq peak counts followed by singular value decomposition (SVD), and 2) modified TF-IDF implemented by Signac[57] followed by SVD. Again, we tested various floors of the number of LSI dimensions and tested whether dropping the first eigenvector, noted in Cusanovich et al.[58] to be influenced by read depth variation, gave improved results. Ultimately, Seurat standard TF-IF featurization, fixing the minimum number of LSI dimensions at 4, and dropping the first eigenvector gave the most competitive model and was subsequently shown in all plots.

We compared MIRA's joint representation with a joint representation of Seurat PCA and LSI embeddings computed by concatenating standardized dimensions from both modes. In the joint test, PCA and LSI were calculated using the best methodology from the independent expression and accessibility tests. Slingshot was subsequently employed on the concatenated embeddings.

For both expression and accessibility modes, MIRA's topic model encoder architecture was allocated one hidden layer of 64 nodes, MIRA's topic models were tuned for 32 iterations using our Bayesian hyperparameter optimization scheme, and the number of topics were constrained to a minimum of 5 and a maximum of 15. The joint KNN graph was calculated with a box-cox transformation parameter of 0.2. Both MIRA and Slingshot lineage inference algorithms are guided by specification of the terminal cell states. Cells that were either the start or end of a lineage were taken to have a read depth equal to the mean of the distribution. Slingshot includes an initial coarse-grain clustering step, and user input indicates which of those clusters represents terminal states. In the automated test, the inclusion of one of the provided ground-truth terminal cells in a cluster designated that cluster as terminal. MIRA determines a subset of cells that are close to a terminal state and the user selects the terminal cell from these possibilities. In the automated test, the user input process was approximated by harmonizing provided terminal cells with the MIRA-determined subset of terminal state cells by finding mutual nearest neighbor pairs between those groups. Provided terminal cells that were not mutual nearest neighbors with any MIRA-determined terminal cell were dropped, as this suggested the provided cell was not placed near the edge of the trajectory topology and did not designate a distinct lineage.

We caution that one cannot draw conclusions of a biological nature based on the synthetically generated benchmark data sets. For example, general conclusions about the relative importance of chromatin accessibility and gene expression in defining cell identity will require investigation of diverse multiomics data sets from diverse biological systems.

In sum, we generated 60 synthetic multimodal datasets with reads derived from a real 10x Genomics single cell multiomics experiment. We systematically varied read depth and scaffold difficulty to assess MIRA's performance and robustness compared to standard approaches, and we evaluated the quality of trajectory inference predictions against a quantitative gold standard using established metrics.

**Software**

Models were implemented using Pyro[7] and PyTorch[43], numerical calculations were implemented using Numpy[44], and statistical tests were conducted with Scipy[22]. Data is stored in the AnnData structure for interoperability with Scanpy[45]. The MIRA analysis package is freely available at https://github.com/cistrome/MIRA.

**Normalized expression**

Raw RNA-seq counts were normalized for streamplot, UMAP, and LITE vs. NITE prediction plot visualization using Scanpy's *normalize_total* function, with *target_sum* set to 10000. In the Share-seq skin dataset, this was followed by nearest-neighbor smoothing with the joint connectivity graph, calculated with the UMAP kernel as implemented by Scanpy.

**Feature selection for expression topic model**

We calculated dispersions and mean counts from log-normalized expression using Scanpy. Genes with mean expression greater than 0.0125 and dispersion greater than 0 were selected as exogenous features. Genes from that group that had dispersion greater than 0.5 were selected as endogenous features for the encoder network.

**Feature selection for accessibility topic model**

All peaks identified for the 10X brain dataset were used as endogenous and exogenous features. All peaks identified for the SHARE-seq skin dataset were used as exogenous features, and 100,000 peaks were randomly selected as endogenous features.

**Data availability**

The authors of the SHARE-seq skin study[62] provide the RNA-seq count matrix at https://www.ncbi.nlm.nih.gov/geo/query/acc.cgi?acc=GSM4156608 and the ATAC-seq peak count matrix at https://www.ncbi.nlm.nih.gov/geo/query/acc.cgi?acc=GSM4156597. 10x Genomics provides the brain dataset[63] RNA-seq count matrix and ATAC-seq peak count matrix at https://www.10xgenomics.com/resources/datasets/fresh-embryonic-e-18-mouse-brain-5-k-1-standard-2-0-0. RNA-seq and ATAC-seq count matrices used for the benchmarking study may be found at https://www.10xgenomics.com/resources/datasets/pbmc-from-a-healthy-donor-granulocytes-removed-through-cell-sorting-10-k-1-standard-2-0-0.

**Data preprocessing**
We used the count matrices provided by the authors of the SHARE-seq skin study[62] for our analysis. CellRanger count matrices were used for the 10x Genomics brain dataset[63].

**Skin dataset cell type selection**
We calculated one expression and one accessibility topic model describing all cells in the SHARE-seq skin data including the hair follicle, interfollicular epidermis (IFE), and mesenchymal cell populations. The joint KNN graph was defined by topics across all cell types and was constructed with a box-cox power transformation parameter of 0.2. Cells were clustered using the Leiden algorithm on the joint KNN graph with a resolution of 2.5. Then, clusters were merged and assigned cell type labels using known skin marker gene expression. Cell type labels were cross-referenced with those provided by the authors of the original SHARE-seq skin study[62]. We then used labeled cells corresponding to the hair follicle or IFE for downstream analyses. Mesenchymal cell populations were not further analyzed. For the hair follicle, we re-calculated the UMAP representation from the joint KNN graph subset. When training RP models, we used all cells in the hair follicle and the IFE, excluding the mesenchymal cells.

**Skin dataset representation comparisons**
We compared representations generated by standard methods using either expression or accessibility data to those generated by MIRA topic modeling. Standard expression-based representations were calculated following Scanpy's recommended workflow. First, count matrices were normalized for read depth by the *normalize_total* function with *target_sum* set to 10000, followed by log-plus-one transformation. Genes with mean expression greater than 0.0125 and dispersion greater than 0.5 (default values) were taken to be highly variable. Next, each genes' log-normalized expression was standardized, and principal component analysis (PCA) was performed on the standardized expression of highly variable genes. A neighborhood graph of cells was then calculated using the PCA representation of the data, which was used to generate a UMAP representation.

Standard accessibility-based representations were calculated based on latent semantic indexing. Latent semantic indexing of ATAC-seq data was calculated using scikit-learn's46 TF-IDF transformation, followed by truncated singular value decomposition (SVD) of all peaks. This was used to calculate a KNN graph with $k$=15 using Euclidean distance on the first 50 eigenvectors, which then was used to generate a UMAP using the default parameters of the umap-learn[29] Python package.

**Skin branch dynamics analysis**
To classify medulla and cortex gene regulation based on branch dynamics, we first found genes with a NITE score > 5 that were differentially-expressed between the two lineages using Scanpy's *rank_gene_groups*. In Fig. 4c, the colors of the bars indicate whether the genes were significantly more highly expressed in the medulla (green) or the cortex (orange) (adjusted p value < 0.1 by Wilcoxon with Benjamini-Hochberg correction and log fold change (LFC) > 1).

We next classified these medulla- or cortex-specific genes based on their expression or accessibility dynamics at the branch:

Expression: we compare the genes' expression in pre-branch matrix cells to post-branch cells (medulla cells for genes in the green bar; cortex cells for genes in the orange bar)

- "High" refers to genes that were already expressed before the branch and did not significantly increase expression after the branch (adjusted p value > 0.1 or LFC < 1)
- "Low" refers to genes that increased in expression after the branch (adjusted p value < 0.1, LFC > 1)

Accessibility: we quantify the chromatin differential at 200 branch point cells

- "High" refers to genes with accessibility at the branch that is higher than expected based on their expression at the branch (chromatin differential > 0.15)
- "Low" refers to genes with accessibility at the branch that is *not* higher than expected based on their expression at the branch (chromatin differential < 0.15)

Therefore, the High Expression-High Accessibility group is composed of medulla- or cortex-specific genes that are already expressed and accessible at the branch.

The Low Expression-High Accessibility group, which we refer to as "branch-primed genes", are medulla- or cortex-specific genes that are more accessible at the branch than would be expected based on their expression at the branch. They subsequently increase in expression levels after the branch in one of the two lineages.

Finally, the Low Expression-Low Accessibility group, which we refer to as "terminal genes", are medulla- or cortex-specific genes that are not yet expressed nor accessible at the branch. Only after the cells have committed to one of the two fates do these genes become accessible and expressed in that lineage.

**Skin primed gene driver TF analysis**

We identified driver TFs of medulla and cortex fate commitment using probabilistic *in silico* deletion. Query sets encompassing the medulla-primed and cortex-primed genes were compared to the background genes that included all other genes for which RP models were trained. The background gene set thus included all highly-variable genes (see Feature selection for expression topic model section), in addition to the top 200 most-activated genes for any topic.

**IFE differentially-expressed terminal genes**

We identified terminally upregulated genes that were differential-expressed between the granular and intermediate granular cell populations in the IFE. First, differentially-expressed genes between granular populations were identified using Scanpy's *rank_gene_groups* (Wilcoxon with Benjamini-Hochberg correction), with adjusted p-value less than 0.1 and log2 fold change between populations greater than 1. Then we selected genes that were differentially-expressed between granular and spinous cells, or between intermediate granular and intermediate spinous cells, with adjusted p-value less than 0.1 and log2 fold change greater than 1. Therefore, we defined terminally upregulated, differentially-expressed genes as those which were both differentially-expressed between granular lineages and upregulated in the granular populations relative to their precursor spinous populations.

**Brain dataset cell type selection**
In addition to the major cell populations described in the main text, the 10X brain dataset also detected a minimal number of Olig1/2+ oligodendrocytes, VWF+/Cdh5+ endothelial cells, and Reelin-positive cells potentially consistent with Cajal-Retzius cells, a transient molecularly and morphologically distinct neuronal population in the developing cerebral cortex. All cells in the dataset were included for MIRA topic modeling, but these minimally detected cell populations were excluded from trajectory analyses due to being too small in number to reliably determine their state along pseudotime and due to the endothelial and Reelin-positive cells being distinct from the major lineages represented in the data. The expression topics exclusively associated with the endothelial and Reelin-positive clusters were therefore also excluded from the later trajectory analyses.

### References


1. Blei, D. M., Ng, A. Y. & Edu, J. B. Latent Dirichlet Allocation Michael I. Jordan. *Journal of Machine Learning Research* **3**, 993–1022 (2003).
2. Kingma, D. P. & Welling, M. Auto-Encoding Variational Bayes. *arXiv:1312.6114* (2013).
3. Hoffman, M. D. *et al.* Stochastic Variational Inference. *Journal of Machine Learning Research* **14**, 1303–1347 (2013).
4. Lopez, R., Regier, J., Cole, M. B., Jordan, M. I. & Yosef, N. Deep generative modeling for single-cell transcriptomics. *Nature Methods 2018 15:12* **15**, 1053–1058 (2018).
5. Choi, K., Chen, Y., Skelly, D. A. & Churchill, G. A. Bayesian model selection reveals biological origins of zero inflation in single-cell transcriptomics. *Genome Biology* **21**, 1–16 (2020).
6. Ioffe, S. & Szegedy, C. Batch Normalization: Accelerating Deep Network Training by Reducing Internal Covariate Shift. *32nd International Conference on Machine Learning, ICML 2015* **1**, 448–456 (2015).
7. Bingham, E. *et al.* Pyro: Deep Universal Probabilistic Programming. *Journal of Machine Learning Research* **20**, 1–6 (2019).
8. Srivastava, A. & Sutton, C. Autoencoding Variational Inference For Topic Models. *5th International Conference on Learning Representations, ICLR 2017 - Conference Track Proceedings* (2017).
9. Dugas, C., Bengio, Y., Bélisle, F., Nadeau, C. & Garcia, R. Incorporating Second-Order Functional Knowledge for Better Option Pricing. in *Proceedings of the 13th International Conference on Neural Information Processing Systems* 451–457 (MIT Press, 2000).
10. Townes, F. W., Hicks, S. C., Aryee, M. J. & Irizarry, R. A. Feature selection and dimension reduction for single-cell RNA-Seq based on a multinomial model. *Genome Biology* **20**, 1–16 (2019).
11. Deng, L. *et al.* Recent advances in deep learning for speech research at Microsoft. *ICASSP, IEEE International Conference on Acoustics, Speech and Signal Processing - Proceedings* 8604–8608 (2013) doi:10.1109/ICASSP.2013.6639345.

12. Srivastava, N., Hinton, G., Krizhevsky, A. & Salakhutdinov, R. Dropout: A Simple Way to Prevent Neural Networks from Overfitting. *Journal of Machine Learning Research* **15**, 1929–1958 (2014).
13. Iyyer, M., Manjunatha, V., Boyd-Graber, J. & Daumé III, H. Deep Unordered Composition Rivals Syntactic Methods for Text Classification. in *Proceedings of the 53rd Annual Meeting of the Association for Computational Linguistics and the 7th International Joint Conference on Natural Language Processing (Volume 1: Long Papers)* 1681–1691 (Association for Computational Linguistics, 2015). doi:10.3115/v1/P15-1162.
14. Kingma, D. P. & Ba, J. L. Adam: A Method for Stochastic Optimization. *3rd International Conference on Learning Representations, ICLR 2015 - Conference Track Proceedings* (2014).
15. Smith, L. N. & Topin, N. Super-Convergence: Very Fast Training of Neural Networks Using Large Learning Rates. 36 (2017) doi:10.1117/12.2520589.
16. Bowman, S. R. *et al.* Generating Sentences from a Continuous Space. *CoNLL 2016 - 20th SIGNLL Conference on Computational Natural Language Learning, Proceedings* 10–21 (2015) doi:10.18653/v1/k16-1002.
17. Bergstra, J., Bardenet, R., Bengio, Y. & Kégl, B. Algorithms for Hyper-Parameter Optimization. in *Advances in Neural Information Processing Systems* (eds. Shawe-Taylor, J., Zemel, R., Bartlett, P., Pereira, F. & Weinberger, K. Q.) vol. 24 (Curran Associates, Inc., 2011).
18. Akiba, T., Sano, S., Yanase, T., Ohta, T. & Koyama, M. Optuna: A Next-generation Hyperparameter Optimization Framework. *Proceedings of the ACM SIGKDD International Conference on Knowledge Discovery and Data Mining* 2623–2631 (2019) doi:10.1145/3292500.3330701.
19. Jamieson, K. & Talwalkar, A. Non-stochastic Best Arm Identification and Hyperparameter Optimization. *Proceedings of the 19th International Conference on Artificial Intelligence and Statistics, AISTATS 2016* 240–248 (2015).
20. Chen, E. Y. *et al.* Enrichr: Interactive and collaborative HTML5 gene list enrichment analysis tool. *BMC Bioinformatics* **14**, 1–14 (2013).
21. Fisher, R. A. On the Interpretation of χ 2 from Contingency Tables, and the Calculation of P. *Journal of the Royal Statistical Society* **85**, 87 (1922).
22. Virtanen, P. *et al.* SciPy 1.0: fundamental algorithms for scientific computing in Python. *Nature Methods 2020 17:3* **17**, 261–272 (2020).
23. Layer, R. M. *et al.* GIGGLE: a search engine for large-scale integrated genome analysis. *Nature Methods* **15**, 123–126 (2018).
24. Egozcue, J. J., Pawlowsky-Glahn, V., Mateu-Figueras, G. & Barceló-Vidal, C. Isometric Logratio Transformations for Compositional Data Analysis. *Mathematical Geology* **35**, 279–300 (2003).
25. Silverman, J. D., Washburne, A. D., Mukherjee, S. & David, L. A. A phylogenetic transform enhances analysis of compositional microbiota data. *eLife* **6**, (2017).
26. Aggarwal Charu C. and Hinneburg, A. and K. D. A. On the Surprising Behavior of Distance Metrics in High Dimensional Space. in *Database Theory — ICDT 2001* (ed. den Bussche Jan and Vianu, V.) 420–434 (Springer Berlin Heidelberg, 2001).

27. Tsagris, M. T., Preston, S. & Wood, A. T. A. A data-based power transformation for compositional data. (2011) doi:10.48550/arxiv.1106.1451.
28. Traag, V. A., Waltman, L. & van Eck, N. J. From Louvain to Leiden: guaranteeing well-connected communities. *Scientific Reports 2019 9:1* **9**, 1–12 (2019).
29. McInnes, L., Healy, J. & Melville, J. UMAP: Uniform Manifold Approximation and Projection for Dimension Reduction. (2018).
30. Fornes, O. *et al.* JASPAR 2020: update of the open-access database of transcription factor binding profiles. *Nucleic Acids Research* **48**, D87–D92 (2020).
31. Korhonen, J. H., Palin, K., Taipale, J. & Ukkonen, E. Fast motif matching revisited: high-order PWMs, SNPs and indels. *Bioinformatics* **33**, 514–521 (2017).
32. Ponte, J. M. & Croft, B. W. A language modeling approach to information retrieval. *Proc. of the 21st annual ACM SIGIR conference on Research and development in information retrieval* 275–281 (1998).
33. Setty, M. *et al.* Characterization of cell fate probabilities in single-cell data with Palantir. *Nature Biotechnology 2019 37:4* **37**, 451–460 (2019).
34. Bergen, V., Lange, M., Peidli, S., Wolf, F. A. & Theis, F. J. Generalizing RNA velocity to transient cell states through dynamical modeling. *Nature Biotechnology* **38**, 1408–1414 (2020).
35. Chen, C. H. *et al.* Determinants of transcription factor regulatory range. *Nature Communications 2020 11:1* **11**, 1–15 (2020).
36. Avsec, Ž. *et al.* Effective gene expression prediction from sequence by integrating long-range interactions. *Nature Methods 2021 18:10* **18**, 1196–1203 (2021).
37. Liu, D. C. & Nocedal, J. On the limited memory BFGS method for large scale optimization. *Mathematical Programming 1989 45:1* **45**, 503–528 (1989).
38. Yadav, A., Goldstein, T. & Jacobs, D. Making L-BFGS Work with Industrial-Strength Nets. *BMVC* (2020).
39. Pearson, E. S. & Naymon, J. On the Use and Interpretation of Certain Test Criteria for Purposes of Statistical Inference. *Biometrika* **20**, 275–240 (1928).
40. Qin, Q. *et al.* Lisa: inferring transcriptional regulators through integrative modeling of public chromatin accessibility and ChIP-seq data. *Genome Biology* **21**, 1–14 (2020).
41. Amaratunga, D. & Cabrera, J. Analysis of Data From Viral DNA Microchips. *Journal of the American Statistical Association* **96**, 1161–1170 (2011).
42. Wilcoxon, F. Individual Comparisons by Ranking Methods. *Biometrics Bulletin* **1**, 80 (1945).
43. Hunter, J. D. Matplotlib: A 2D Graphics Environment. *Computing in Science  Engineering* **9**, 90–95 (2007).
44. Savitzky, A. & Golay, M. J. E. Smoothing and Differentiation of Data by Simplified Least Squares Procedures. *Analytical Chemistry* **36**, 1627–1639 (2002).
45. PBMC from a healthy donor - granulocytes removed through cell sorting (10k) Single Cell Multiome ATAC + Gene Expression Dataset by Cell Ranger ARC 2.0.0. (2021).
46. Zheng, G. X. Y. *et al.* Massively parallel digital transcriptional profiling of single cells. *Nature Communications 2017 8:1* **8**, 1–12 (2017).
47. Zhang, Y. *et al.* Model-based analysis of ChIP-Seq (MACS). *Genome Biology* **9**, 1–9 (2008).

48. Buenrostro, J. D., Giresi, P. G., Zaba, L. C., Chang, H. Y. & Greenleaf, W. J. Transposition of native chromatin for fast and sensitive epigenomic profiling of open chromatin, DNA-binding proteins and nucleosome position. *Nature Methods 2013 10:12* **10**, 1213–1218 (2013).
49. He, H. H. *et al.* Refined DNase-seq protocol and data analysis reveals intrinsic bias in transcription factor footprint identification. *Nature methods* **11**, 73–78 (2014).
50. Corces, M. R. *et al.* The chromatin accessibility landscape of primary human cancers. *Science* **362**, (2018).
51. Alon, U. An Introduction to Systems Biology; Design Principles of Biological Circuits; Second Edition.
52. Saelens, W., Cannoodt, R., Todorov, H. & Saeys, Y. A comparison of single-cell trajectory inference methods. *Nature Biotechnology 2019 37:5* **37**, 547–554 (2019).
53. Street, K. *et al.* Slingshot: Cell lineage and pseudotime inference for single-cell transcriptomics. *BMC Genomics* **19**, 1–16 (2018).
54. Lewis, B. W. & Baglama, J. The irlba Package. (2012).
55. Hao, Y. *et al.* Integrated analysis of multimodal single-cell data. *Cell* **184**, 3573-3587.e29 (2021).
56. Thorndike, R. L. Who belongs in the family? *Psychometrika 1953 18:4* **18**, 267–276 (1953).
57. Stuart, T., Srivastava, A., Madad, S., Lareau, C. A. & Satija, R. Single-cell chromatin state analysis with Signac. *Nature Methods 2021 18:11* **18**, 1333–1341 (2021).
58. Cusanovich, D. A. *et al.* A Single-Cell Atlas of In Vivo Mammalian Chromatin Accessibility. *Cell* **174**, 1309-1324.e18 (2018).
59. Paszke, A. *et al.* PyTorch: An Imperative Style, High-Performance Deep Learning Library. in *Advances in Neural Information Processing Systems 32* (eds. Wallach, H. et al.) 8024–8035 (Curran Associates, Inc., 2019).
60. Harris, C. R. *et al.* Array programming with NumPy. *Nature 2020 585:7825* **585**, 357–362 (2020).
61. Wolf, F. A., Angerer, P. & Theis, F. J. SCANPY: Large-scale single-cell gene expression data analysis. *Genome Biology* **19**, 1–5 (2018).
62. Ma, S. *et al.* Chromatin Potential Identified by Shared Single-Cell Profiling of RNA and Chromatin. *Cell* **183**, 1103-1116.e20 (2020).
63. Fresh embryonic E18 mouse brain (5k). Single Cell Multiome ATAC + Gene Expression Dataset by Cell Ranger ARC 2.0.0. (2021).
64. Pedregosa, F. *et al.* Scikit-learn: Machine Learning in Python. *Journal of Machine Learning Research* **12**, 2825–2830 (2011).
